# HIV broadly neutralizing antibody escape dynamics drive the outcome of AAV vectored immunotherapy in humanized mice

**DOI:** 10.1101/2024.07.11.603156

**Authors:** Nicolas M.S. Galvez, Adam D. Nitido, Seo Bin Yoo, Yi Cao, Cailin E. Deal, Christine L. Boutros, Scott W. MacDonald, Yentli E. Soto Albrecht, Evan C. Lam, Maegan L. Sheehan, Dylan Parsons, Allen Z. Lin, Martin J. Deymier, Jacqueline M. Brady, Benjamin Moon, Christopher B. Bullock, Serah Tanno, Amarendra Pegu, Xuejun Chen, Cuiping Liu, Richard A. Koup, John R. Mascola, Vladimir D. Vrbanac, Daniel Lingwood, Alejandro B. Balazs

## Abstract

Broadly neutralizing antibodies (bNAbs) have shown promise for prevention and treatment of HIV. Potency and breadth measured *in vitro* are often used as predictors of clinical potential; however, human studies demonstrate that clinical efficacy of bNAbs can be undermined by both pre-existing and *de novo* resistance. Here we find that HIV-infected humanized mice receiving bNAbs delivered via AAV as Vectored ImmunoTherapy (VIT) can be used to identify antibody escape paths, which are largely conserved for each bNAb. Path selection, and consequent therapeutic success, is driven by the fitness cost and resistance benefit of emerging mutations. Applying this framework, we independently modulated bNAb resistance or the fitness cost of escape mutants, resulting in enhanced efficacy of VIT. This escape path analysis successfully explains the therapeutic efficacy of bNAbs, and enables a tractable means of quantifying and comparing the potential for viral escape from therapeutics *in vivo*.

**Graphical Abstract:** 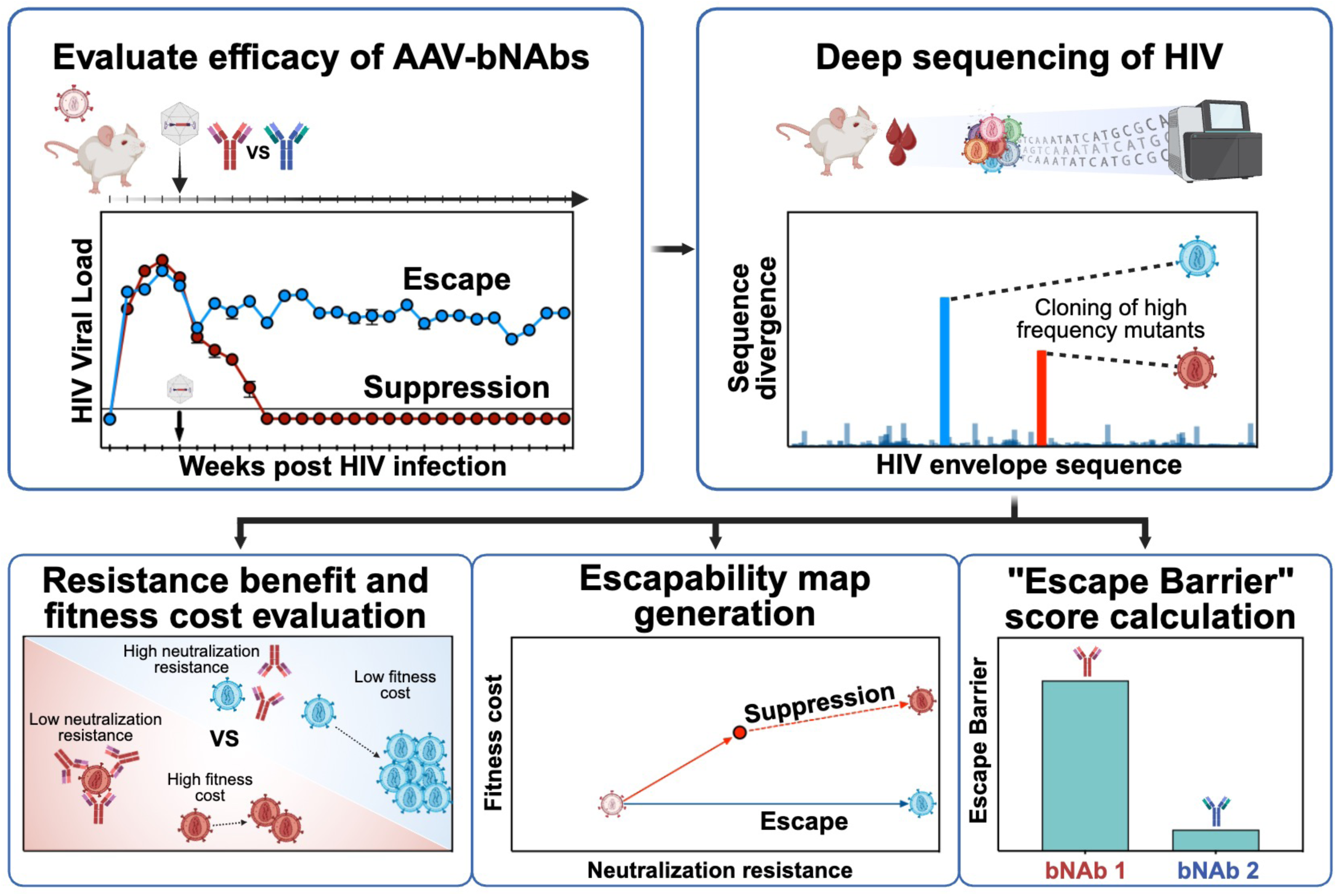

## Introduction

Despite substantial efforts to develop therapies and vaccines, HIV-1 remains a global pandemic. Current combination antiretroviral therapy (cART) regimens can effectively prevent and treat HIV infections, and emerging long-acting agents such as Lenacapavir show remarkable efficacy in preventing viral acquisition^1,2^. However, their therapeutic success still requires strict adherence to the regimen, as viral rebound is swiftly observed upon treatment interruption and may lead to the development of cART resistance^1^. This is of particular relevance when access to healthcare is interrupted^3^. Over the past decade, broadly neutralizing antibodies (bNAbs) have been extensively described and characterized^4,5^. These antibodies target different sites of vulnerability on the HIV envelope (Env) protein and can prevent infection across HIV clades^4^. Their clinical success is often predicted by using metrics determined *in vitro* across large panels of diverse viral isolates^6^. These measurements consist of potency, the average inhibitory concentration needed to block 50% infection (IC_50_) across all tested isolates, as well as breadth, the fraction of isolates neutralized at a given antibody concentration^7^. Recent advances in antibody isolation have identified new bNAbs with median potencies as low as 0.003 μg/mL^8,9^ and breadth capable of neutralizing up to 96% of a panel of 208 viruses at an IC_50_ of <1 μg/mL^10^.

Given their potential, significant efforts have been made to advance bNAb-based therapies, with multiple studies demonstrating their success in preventing HIV infection in mouse^11–16^ and non-human primate (NHP) models^17–21^. These findings paved the way for the Antibody Mediated Protection (AMP) studies (NCT02716675 and NCT02568215), two international harmonized phase 2b randomized controlled trials assessing the capacity of the CD4 binding site (CD4bs)-targeting bNAb VRC01 to prevent HIV-1 acquisition^22^. While VRC01 prevented transmission of neutralization-sensitive strains, overall efficacy was limited by a large fraction of circulating strains with mutations conferring resistance to neutralization^23,24^. Numerous additional studies have also demonstrated the potential for bNAb administration to transiently suppress HIV viremia in animal models^25–32^ as well as in people living with HIV (PLWH) who harbor sensitive viruses^33–40^. However, these studies also revealed that even combinations of two or three bNAbs targeting different sites of vulnerability can fail to control viremia due to the emergence of escape mutations^38,41,42^.

The clinical translation of bNAbs is also hindered by the need for repeated infusions to maintain serum concentrations above a therapeutic threshold^43^. To overcome this, we and others have described Vectored Immunoprophylaxis (VIP)^12^, which utilizes a single intramuscular (IM) injection of recombinant adeno-associated viral (AAV) vectors engineered to encode a bNAb transgene ^44–47^. VIP induces the production of durable and protective concentrations of antibodies *in vivo*^48^. Studies of VIP with VRC07, a CD4bs-targeting bNAb clonally related to VRC01 but with greater potency and breadth^8,17^, have demonstrated that delivery via AAV8 prevents intravenous and mucosal transmission of HIV in humanized mice^11,12,16^ and of SIV/SHIV in NHP models^49,50^. Recently, the VRC603 trial administered AAV8-VRC07^11,51^ to cART-suppressed patients (NCT03374202). In this study, volunteers reported up to 3 µg/mL of VRC07 in serum for at least three years following a single IM administration^51^. Despite these promising results, anti-drug antibodies (ADAs) emerged in a subset of participants, highlighting one of the challenges of this approach. However, recent studies suggest that ADAs may be less likely to emerge in settings of increased immune tolerance, such as transient immunosuppression of adult NHPs^52^ or in pediatric settings^53^.

The therapeutic potential of bNAbs delivered via AAV as Vectored ImmunoTherapy (VIT) to cure an established HIV infection is an area of active investigation, given its potential use in resource-limited settings where ART adherence is suboptimal^53^. In particular, the success of VIT can be hindered by the selection of escape variants, as seen during previous trials of passively transferred bNAbs^54^. Here, we present the use of VIT in HIV-infected humanized mice as an experimental model to evaluate viral suppression and profile the emergence of Env mutations leading to viral escape from individual vectored antibodies. We chose two widely used Clade B, Tier 2 isolates for this: HIV_JR-CSF_, a chronic virus cloned from the CSF of a person who died of severe AIDS encephalopathy^55^; and HIV_REJO.c_, a transmitter/founder isolate that was computationally derived as the likely strain that initiated a heterosexual transmission^56^. Both of these isolates represent clinically relevant viruses, as they were cloned soon after isolation and have not been extensively cultured *in vitro* unlike laboratory adapted strains (e.g., HIV_NL4-3_)^57^. To our surprise, AAV8-VRC07 treatment drove viral evolution towards a high fitness cost escape path that often failed to emerge, enabling long-term suppression of HIV_REJO.c_. Interestingly, while necessary for activity, traditional breadth and potency metrics were insufficient to predict therapeutic outcome; instead, the specific fitness cost and resistance benefit tradeoffs of the primary escape paths best explained the success of VIT. To understand this dynamic, we adapted the fitness landscape^58^ framework to create an ‘escapability’ map for each antibody-strain combination, effectively explaining VIT outcomes by identifying the most traversed escape paths during vectored antibody therapy. Our data show that escape pathway dynamics are a key component to understanding the *in vivo* efficacy of antibody-mediated therapies against HIV, and that future therapeutic strategies that increase the fitness cost of the primary escape paths are essential for clinical success.

## Results

### VRC07, but neither N6 nor PGDM1400, suppresses HIV_REJO.c_ replication in humanized mice

To explore whether bNAbs delivered via AAV8 as VIT could be used as a long-lived therapeutic strategy for suppressing ongoing HIV replication, we employed the Bone marrow-Liver-Thymus (BLT) humanized mouse model^59^. This model exhibits robust engraftment of human lymphocytes across tissues, enabling sustained infection with HIV^60,61^. BLT mice were infected with HIV_REJO.c_, a clade B, Tier 2, CCR5-tropic, transmitted/founder infectious molecular clone (IMC) of HIV^56^. Prior to treatment, the virus was allowed to propagate and diversify in the host over a period of four weeks (Figure 1A). Groups of BLT mice harboring established HIV_REJO.c_ infections were given a single IM injection of 5×10^11^ genome copies (GC) AAV8 expressing either VRC07, N6, or PGDM1400 bNAbs or Luciferase as a negative control. Importantly, all the tested bNAbs potently neutralized HIV_REJO.c_ *in vitro* (Figure S1A, Table S1). After vector administration, antibody expression levels in all mice increased over a period of six weeks (Figure 1B).

**Figure 1.**
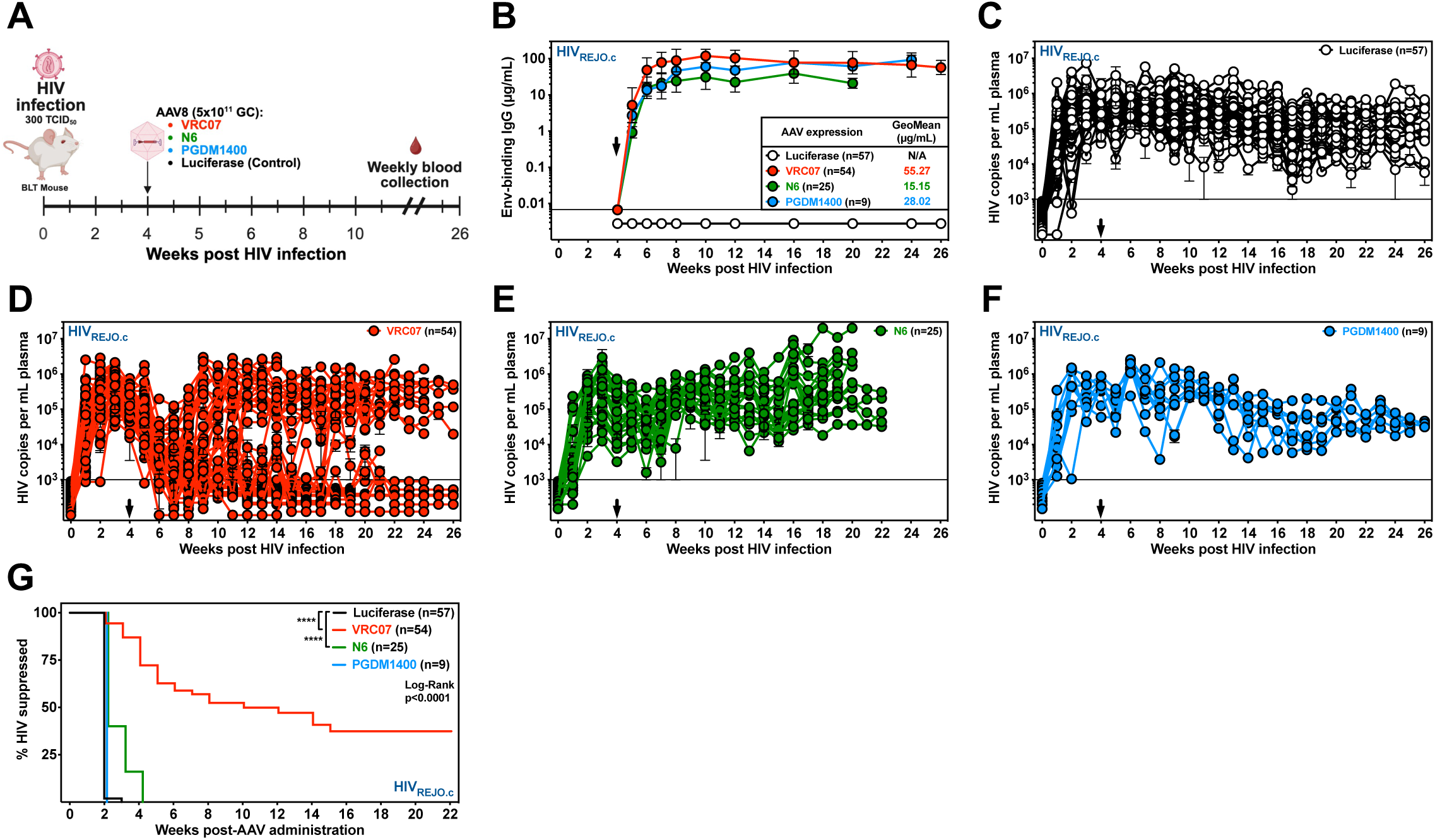
Vectored delivery of VRC07, but not PGDM1400 or N6, can suppress established HIV_REJO.c_ infection. **(A)** BLT humanized mice were infected with 300 TCID_50_ of HIV_REJO.c_ and the viral population was allowed to replicate for 4 weeks. Mice were then IM injected with 5×10^11^ genome copies (GC) of AAVs encoding for either VRC07, N6, PGDM1400, or Luciferase as control. Mice were followed for 6 months, and blood samples were collected weekly. **(B)** ELISA-based quantitation of gp120-binding antibodies in the serum of HIV_REJO.c_-infected humanized mice following administration of AAV8-Luciferase, AAV8-VRC07, AAV8-N6, or AAV8-PGDM1400 vectors. Black arrow denotes vector administration. Data are plotted as geometric mean ± geometric SD. **(C-F)** HIV viral load in plasma of HIV_REJO.c_-infected mice injected with AAV8-Luciferase as control **(C)**, AAV8-VRC07 **(D)**, AAV8-N6 **(E)**, or AAV8-PGDM1400 **(F)**. Black arrows denote vector administration. Each colored line depicts an individual mouse tracked over time. The sensitivity of qPCR was 1 genome copy per µL of plasma, and 5 µL were used in the reaction, resulting in a 1000 copy per mL limit of detection (solid line). Data are presented as mean ± S.E.M. **(G)** Kaplan-Meier plot of viral suppression in BLT humanized mice infected with HIV_REJO.c_ given the indicated bNAb-expressing vector. The total model significance (p<0.0001) and pairwise comparison against the Luciferase control (****: p<0.0001) were assessed independently using Log-rank (Mantel-Cox) tests. The percentage of HIV suppressed was defined as the fraction of mice that were not escaped as described in the methods.

For AAV8-VRC07-treated mice, the bNAb expression level plateaued at a geometric mean of ∼55 μg/mL in plasma, which was sustained for the remainder of the study; a value approximately 2,000 times over its *in vitro* IC_50_ value (Figure S1B). Infected mice exhibited mean plasma viral loads averaging 5.34×10^5^ copies per mL of plasma by week four and sustained viremia following AAV8-Luciferase treatment for the remaining 22 weeks of the study (Figure 1C). In contrast, following AAV8-VRC07 treatment, viral loads decreased precipitously over a three-week period, which diverged into two outcomes: viral rebound in 56% (30/54) or sustained control in 44% (24/54) of mice throughout the study (Figure 1D, Figure S1C). Notably, this trend was observed across five independent experiments, each using a unique donor for each batch of BLT mice (Table S2).

To explore whether another CD4bs-directed bNAb could achieve long-lived viral suppression we tested N6^62^, an exceptionally broad bNAb with an *in vitro* IC_50_ against HIV_REJO.c_ similar to that of VRC07 (Figure S1A). In AAV8-N6 treated mice, the plasma concentrations of N6 achieved a geometric mean of ∼15 μg/mL, over 300 times above the *in vitro* IC_50_ against HIV_REJO.c_ (Figure 1B, Figure S1B). However, AAV8-N6 treatment only resulted in a transient decline in viral load over the first two weeks, which returned to pre-treatment levels within four weeks of vector injection (Figure 1E).

Finally, we also tested PGDM1400^63^, a somatic variant of the PGT145 antibody family that recognizes the HIV envelope trimer V1/V2 apex with exceptional potency. This bNAb neutralized HIV_REJO.c_ with approximately five-fold greater potency than those measured for VRC07 and N6 (Figure S1A, Table S1). Following AAV8-PGDM1400 administration, expression of PGDM1400 rapidly increased, reaching a geometric mean plasma concentration of ∼28 μg/mL, or nearly 3,700 times above the *in vitro* IC_50_ against HIV_REJO.c_ (Figure 1B, Figure S1B). However, all mice treated with AAV8-PGDM1400 displayed no significant change in viremia over the 26-week period of observation (Figure 1F). Comparison of the geometric mean viral loads of all evaluated mice demonstrated that only those animals receiving AAV8-VRC07 exhibited a significant decline in viral load compared to the AAV8-Luciferase control. (Figure S1D).

To determine whether differences in viral population structure prior to antibody treatment might explain the observed outcome, we compared the average diversity (Shannon entropy) of the HIV *env* gene three weeks after infection, but prior to vector administration, and noted no significant differences between any of the different groups (Figure S1E). When comparing rebounding AAV8-VRC07 treated mice to suppressed mice, we found that mice in each group had indistinguishable expression levels of VRC07 antibody (Figure S1F), but rebounding mice had a 1.6 times higher viral load compared to suppressed mice at the time of AAV administration (Figure S1F).

Together, these studies show that only AAV8-VRC07 could suppress actively replicating HIV_REJO.c_ for an extended duration. In contrast, AAV8-N6 treatment resulted in a small but significant delay in viral escape as compared to the AAV8-Luciferase control; while AAV8-PGDM1400 was unable to significantly delay viral escape, despite comparable viral heterogeneity and higher bNAb potency (Figure 1G).

### Antibody escape paths are conserved for each broadly neutralizing antibody

To understand the basis for the observed variation in bNAbs effectiveness against HIV_REJO.c_, we mapped the escape paths traversed by the virus to acquire resistance to each individual bNAb. Plasma viral RNA was obtained from each mouse endpoint and used to amplify viral Env for deep sequencing to analyze non-synonymous mutations that arose during viral escape. Individual HIV-infected mice receiving AAV8-Luciferase exhibited mutations distributed throughout the HIV_REJO.c_ envelope (Figure S2A, Control); however, these mutations followed no discernable pattern on the HIV_REJO_ *env* gene (Figure S2B) and did not significantly impact the distribution of potential N-linked glycosylation sites (PNGs) on this protein (Figure S2C).

Most Env sequences obtained from HIV_REJO.c_-infected animals that escaped from VRC07 exhibited two pairs of potential escape mutations: either two D-loop mutations (N276D and D279A) or a D-loop and a V5-loop mutation (D279A and N460D, respectively) (Figure S2A, VRC07). When assessed collectively across all escaped mice, these three mutations encompassed the majority of amino acid divergence from the parental (WT) sequence (Figure 2A). Notably, the N460D mutation also resulted in a PNGs loss in the V5-loop of HIV_REJO.c_ (Figure S2D).

**Figure 2.**
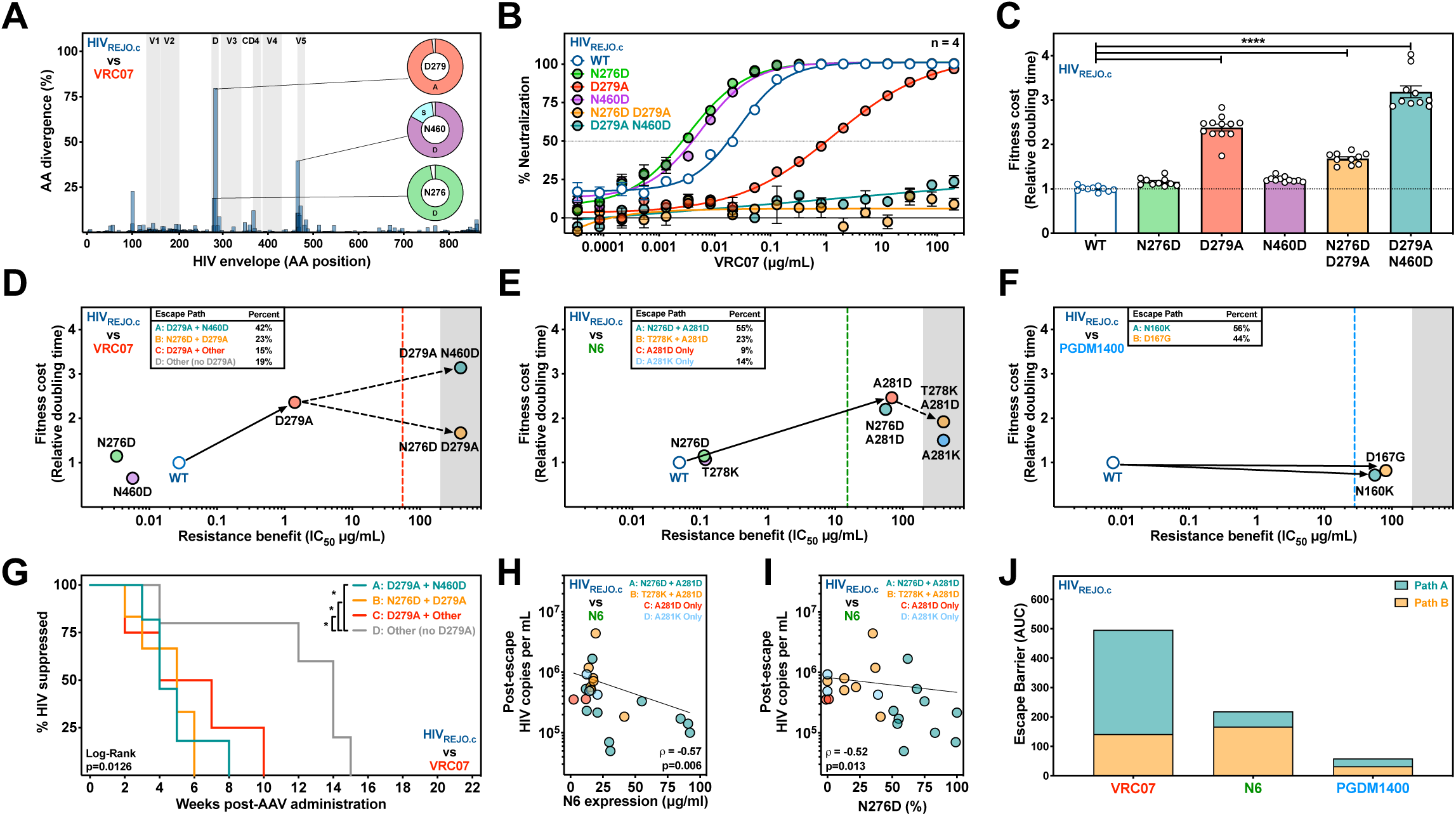
Fitness cost and resistance benefit accrued along escape paths determine the difficulty of HIV_REJO._ escape from each bNAb. **(A)** Amino acid divergence from the envelope gene of the HIV_REJO.c_ parental strain across all AAV8-VRC07 treated mice. Sequences were determined by Illumina Deep Sequencing of the viral envelope isolated from plasma at the final experimental timepoint. The X-axis represents the envelope protein amino acid position relative to HIV_HXB2_ numbering. The Y-axis represents the percentage of average amino acid divergence from the parental strain, corrected for divergence observed in control mice. Pie charts represent the most common amino acid mutations for sites with the highest divergence. See also Figure S2. **(B)** *In vitro* neutralization assays of each HIV_REJO.c_ mutant identified in **(A)** against VRC07. Data are plotted as mean ± S.E.M. Each data point was evaluated in quadruplicate. **(C)** Relative viral growth of each HIV_REJO.c_ mutant identified in **(A)**. Growth rates were determined in activated CD4^+^ T cells performing *QuickFit* assays and normalized to the parental strain. Data are plotted as mean ± S.E.M. and statistical differences were assessed by a Kruskal-Wallis non-parametric ANOVA with Dunn’s *post hoc* test to correct for multiple comparisons (****: p<0.0001). See also Figure S2. **(D-F)** Escapability maps denoting fitness cost (Y-axis, relative doubling time shown in (**C**)) and resistance benefits (X-axis, neutralization resistance shown in (**B**)) for each HIV_REJO.c_ mutant observed during escape from VRC07 (**D**), N6 (**E**), or PGDM1400 (**F**). Dashed vertical line denotes the geometric mean antibody serum concentration after vectored bNAb administration. Shaded areas represent the neutralization assay limit of detection. Solid arrows represent the likely initial path taken with dashed arrows representing the likely second step paths to escape. In the map legend, each escape path is sorted based on the relative frequency observed in the sequencing data. See also Figures S2 and S3. **(G)** Survival analysis of HIV_REJO.c_-infected mice treated with AAV8-VRC07 by escape path haplotype. The total model significance (p=0.0126) and pairwise comparisons of the non-D279A escape paths to each other escape path (*: p<0.05) were assessed independently using Log-rank (Mantel-Cox) tests. See also Figure S3. **(H)** Correlation plot of N6 geometric mean-expression vs post-escape HIV_REJO.c_ viral load. The line represents the semi-log least squares regression, ρ represents the Spearman correlation value determined for the data, with the associated p-value below. See also Figure S3. **(I)** Correlation plot of the post-N6 escape viral load of HIV_REJO.c_ vs the frequency of N276D determined by viral envelope sequencing. The line represents the semi-log least squares regression, ρ represents the Spearman correlation value determined for the data, with the associated p-value below. See also Figure S3. **(J)** Escape Barrier (AUC) Score denoting the aggregate fitness cost for each escape path as determined in the escapability maps. This score was calculated by adding the area under the escapability plot for escape paths A and path B viral escapes from VRC07, N6, and PGDM1400. See also Figure S4.

To determine the impact of these mutations on VRC07 resistance, each individual mutation and combination of mutations observed in haplotypes were engineered into HIV_REJO.c_ IMC plasmids to produce replication-competent Env mutant viruses. Each mutant virus was tested for neutralization sensitivity against VRC07 *in vitro* (Figure 2B). When assessed individually, neither N276D nor N460D conferred any resistance to VRC07 neutralization relative to the WT HIV_REJO.c_. However, D279A led to partial escape by mediating a 51-fold increase in the IC_50_ to 1.4 μg/mL and a reduction of the dose-response curve slope (Table S1) ^64^. Of note, this new IC_50_ still remained well below the geometric mean serum concentration of VRC07 achieved *in vivo*. However, complete escape from VRC07 was only observed for envelopes containing both D279A and either N460D or N276D (Figure 2B).

To measure the fitness cost associated with VRC07 escape mutations emerging in HIV_REJO.c_, we determined the growth rate of each IMC envelope mutant using QuickFit, an RT-qPCR-based assay that quantifies viral growth in activated primary CD4^+^ T cells across large numbers of replicates (Figure S2E)^65^. The doubling time of each IMC harboring escape mutations was then normalized to the WT virus to determine specific fitness costs (or benefits) (Figure 2C). The D279A mutation incurred a significant fitness cost, whereas no changes were observed for the N276D or N460D mutations alone. Strains containing paired mutations also exhibited a higher fitness cost compared to the WT strain. Altogether, these results show that HIV_REJO.c_ follows conserved evolutionary paths to achieve complete escape from VRC07 with differential impact on fitness cost and antibody resistance benefit.

The escape paths traversed by HIV_REJO.c_ to acquire resistance to N6 were also evaluated (Figure S2A, N6). Three mutations in or around the D-loop were the most frequent (Figure S2F), with one of them resulting in loss of a PNGs (Figure S2G). Unlike the HIV_REJO.c_ escape paths from VRC07, one mutation on the D-loop, namely A281D, was sufficient to escape from levels of N6 achieved *in vivo* (Figure S2H, Table S1). Moreover, when A281D was paired with T278K, resistance increased, whereas pairing with N276D modestly decreased resistance. Additionally, A281K led to complete escape from N6, although it required three nucleotide mutations from the parental sequence (Figure S2H). Using the aforementioned QuickFit assay, the fitness cost of all identified individual and haplotype combinations of mutations were evaluated (Figure S2I,J). Similar to the HIV_REJO.c_ VRC07 escape paths, the initial A281D and the combinations with N276D or T278K, resulted in a higher fitness cost relative to the WT strain (Figure S2I,J).

Lastly, the escape paths traversed by HIV_REJO.c_ to acquire resistance to PGDM1400 were evaluated (Figure S2A, PGDM1400). Mutations were observed surrounding the V1/V2 trimer apex region, with N160K and D167G being the most prevalent (Figure S2K), with the former resulting in the loss of a PNGs (Figure S2L). As expected, both of these mutations were sufficient to confer complete escape from PGDM1400, reaching IC_50_ values above the average level of expression achieved *in vivo* (Figure S2M, Table S1). N160K and D167G resulted in a fitness benefit compared to WT strain (Figure S2N,O).

Together, these results show that *in vivo* escape from bNAbs in humanized mice follows reproducible paths, each with distinct fitness costs and resistance benefits. In light of the abundance of escape paths, we sought to understand why a given pathway was traversed as opposed to another.

### The Escape Barrier of each bNAb is determined by the fitness landscape of escape paths

To create an escapability map of the fitness landscape in the context of VIT, we projected individual escape mutations onto a two-dimensional axis of replicative fitness cost and resistance benefit (Figure S2P). This was replicated for each evaluated bNAb, with each point representing an identified escape mutation and arrows representing the potential escape paths (Figure 2D-F). We categorized each mouse sample into an escape path by using the viral haplotype frequencies of the dominant escape mutations (Figure S3A-F).

To escape VRC07 *in vivo*, 81% of the HIV_REJO.c_ sequences harbored the D279A mutation, conferring partial escape from VRC07, albeit with a high fitness cost (Figure 2D). These escaped viruses also acquired a secondary mutation, typically N460D (Path A) or N276D (Path B), leading to complete escape from VRC07. Path A escape variants (D279A + N460D) incurred an even higher fitness cost than the D279A-only escape variants; whereas path B escape variants (N276D + D279A) improved their fitness relative to the D279A-only variants. Notably, 19% of the escapes did not harbor the D279A amino acid change as a partial escape (Path D), but rather found an alternative escape pathway. These non-canonical escape paths exhibited a significant delay in time to escape (Figure 2G), suggesting that these viral populations incurred a higher fitness cost than D279A-based escape paths. To identify the potential forces driving a given population into a specific path, we compared the VRC07 expression levels (Figure S3G), the post-escape viral load (Figure S3J), and the total number of accumulated mutations (Figure S3M) for each path, but failed to find any significant differences across the paths.

The same analysis was performed for N6 (Figure 2E), where 86% of the escape variants harbored an A281D mutation, conferring complete escape from N6, with an IC_50_ 4.5 times greater than the average *in vivo* expression. These A281D variants typically had a second mutation, either N276D (Path A) or T278K (Path B). Compared to A281D-only variants, path A escapes were marginally more fit and less resistant to N6; while path B escapes were both more fit and more resistant to N6 (Figure 2E). A mix of both path A and path B haplotypes was found within most samples (Figure S3E). Notably, samples with higher serum concentrations of N6 were associated with lower post-escape viral loads (Figure 2H), suggesting that N6 may impair viral replication. This was particularly evident for samples with higher frequencies of N276D (Path A) (Figure 2I). A small number of N6 escape sequences had three nucleotide mutations resulting in A281K, which completely escaped N6 with a minor fitness cost (Figure 2E). There were also some escaped samples harboring only the A281D variant, with no other prevalent mutations; however, the N6 serum levels of these samples were 6.4-fold lower compared to those of path A samples (Figure S2H). Of note, given that these were only two samples, statistical differences could not be assessed. The post-escape viral load (Figure S2K) and the total number of accumulated mutations (Figure S2N) were similar across the different paths.

Finally, the PGDM1400 escape paths were clustered based on the haplotypes of N160K and D167G (Figure 2F, Figure S3C and S3F). N160K (Path A) was found in 56% of the escape variants and the remaining 44% contained D167G (Path B). Both paths led to complete escape from PGDM1400, with an IC_50_ 2.0- or 2.9-times greater than the average *in vivo* expression (Figure 2F). We did not observe any significant difference between the PGDM1400 expression levels and post-escape viral loads for the two escape paths (Figure S3I and 3L, respectively). However, Path B escapes accumulated significantly more mutations compared to Path A escapes (Figure S3O).

In order to understand the cumulative fitness cost of HIV_REJO.c_ escape from VRC07, N6, and PGDM1400, we used the escapability maps to determine Escape Barrier scores. Briefly, for each escape path on the map, we calculated the area under the curve (AUC) summing the fitness cost across the IC_50_ values from the WT to the escaped strain (Figure S4A-C). We then combined the AUC of the top two escape paths, which accounted for the majority of the observed haplotypes, into a final Escape Barrier score (Figures S4D-F). Interestingly, the ranking of the Escape Barrier scores for VRC07, N6, and PGDM1400 corresponded with *in vivo* experimental outcomes (Figure 2J).

### HIV_JR-CSF_ rapidly escapes from bNAbs due to low fitness cost escape mutations

Given the ability of AAV8-VRC07 to reproducibly suppress HIV_REJO.c_ in a subset of mice, we sought to determine whether this could be replicated with another HIV isolate. HIV_JR-CSF_ is a Tier 2, Clade B, CCR5-tropic primary isolate originally obtained from the cerebrospinal fluid of a person living with HIV^55^. Groups of humanized mice were infected with an HIV_JR-CSF_ IMC viral stock four weeks prior to receiving AAV8-Luciferase or AAV8-bNAbs (Table S2). Antibody expression stabilized within four weeks of AAV administration, achieving geometric mean plasma concentrations tens to hundreds of times above the *in vitro* IC_50_ (Figure 3A, Figure S5A,B). In contrast to HIV_REJO.c_, HIV_JR-CSF_-infected mice exhibited no changes in viral load following administration of any of the AAV8-bNAbs evaluated and were indistinguishable from the AAV8-Luciferase controls (Figure 3B, Figure S5C-F).

**Figure 3.**
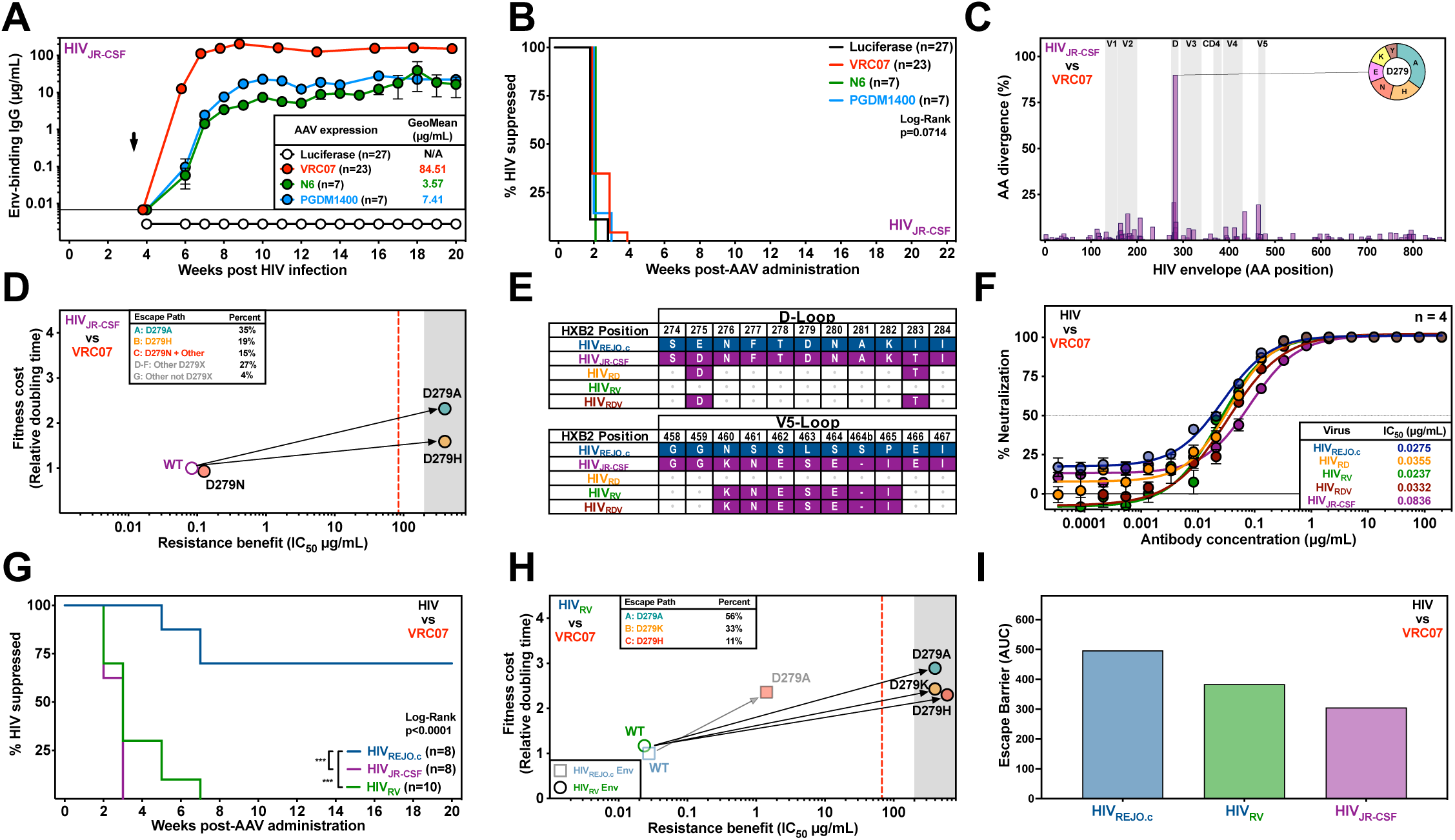
HIV_REJO.c_ V5-loop restricts escape from VRC07, by decreasing the resistance benefit of D279 mutations. **(A)** ELISA-based quantitation of gp120-binding antibodies in the serum of HIV_JR-CSF_-infected humanized mice following administration of 5×10^11^ genome copies (GC) of AAV8-Luciferase, AAV8-VRC07, AAV8-N6, or AAV8-PGDM1400 vectors. Black arrow denotes vector administration. Data are plotted as geometric mean ± geometric SD. **(B)** Kaplan-Meier plot of viral suppression in humanized mice infected with HIV_JR-CSF_ given the indicated bNAb-expressing vector. The total model significance was assessed using a Long-rank (Mantel-Cox) test (p=0.0714). See also Figure S5. **(C)** Amino acid divergence from the envelope gene of the HIV_JR-CSF_ parental strain across all AAV8-VRC07 treated mice. Sequences were determined by Illumina Deep Sequencing of the viral envelope isolated from plasma at the final experimental timepoint. The X-axis represents the envelope protein amino acid position relative to HIV_HXB2_ numbering. The Y-axis represents the percentage of average amino acid divergence from the parental strain, corrected for divergence observed in control mice. Pie chart represents the most common amino acid mutations for the site with the highest divergence. See also Figure S6. **(D)** Escapability map of HIV_JR-CSF_ escape from VRC07. Dashed vertical line denotes the geometric mean antibody serum concentration after AAV8-VRC07 administration. Shaded area represents the neutralization assay limit of detection. Arrows represent the likely path taken to escape. See also Figures S6 and S8. **(E)** Alignment of the D-loop and the V5-loop amino acid sequences for HIV_REJO.c_, HIV_JR-CSF_, and the HIV_RD_, HIV_RV_, and HIV_RDV_ chimeras. **(F)** *In vitro* neutralization of HIV_REJO.c_, HIV_RD_, HIV_RV_, HIV_RDV_, and HIV_JR-CSF_ by VRC07 bNAb. Data are plotted as mean ± S.E.M. Each datapoint was evaluated in quadruplicate. **(G)** Kaplan-Meier plot of viral suppression in humanized mice infected with HIV_REJO.c_, HIV_RV_, or HIV_JR-CSF_ following vectored VRC07 administration. The total model significance (p<0.0001) and pairwise comparisons against HIV_REJO.c_ (***: p<0.001) were assessed independently using Log-rank (Mantel-Cox) tests. See also Figure S7. **(H)** Escapability map of HIV_RV_ during escape from VRC07. Lighter square symbols represent the original HIV_REJO.c_ escape path taken against VRC07. Dashed vertical line denotes geometric mean bNAb concentrations after vectored VRC07 administration. Shaded areas represent the limits of detection. Arrows represent the likely path to escape. See also Figures S7 and S8. **(I)** Escape Barrier (AUC) Score quantifying the difficulty of escape from VRC07 for HIV_REJO.c_, HIV_RV_, and HIV_JR-CSF_. See also Figure S8.

We deep sequenced HIV_JR-CSF_ Env from each mouse to identify mutations associated with escape from VRC07, N6, and PGDM1400 (Figure S6A). Interestingly, mice receiving AAV8-Luciferase exhibited some consistent mutations distributed throughout the HIV_JR-CSF_ envelope despite the absence of bNAb selection pressure, suggesting adaptation to the host (Figure S6B). This was most apparent at position N339 that translated into loss of a PNGs near the V3-loop, which was not observed for HIV_REJO.c_ (Figure S6C). Other mutations around the V2- and V4-loops were also found at higher rates than for HIV_REJO.c_.

To escape VRC07, HIV_JR-CSF_ acquired a single D-loop mutation at position D279, which was observed across all samples (Figure 3C), and a minority of samples exhibited a loss of a PNGs at N276 as also seen for HIV_REJO.c_ (Figure S6D). The IC_50_ of HIV_JR-CSF_ harboring D279 escape mutations were evaluated, with D279A and D279H achieving complete escape with an IC_50_ greater than 200 µg/mL, while D279N exhibited a modest increase in IC_50_ relative to the WT strain (Figure S6E, Table S2). D279A exhibited an increased fitness cost relative to the parental strain, while D279H and D279N had no statistically significant fitness cost (Figure S6F,G). Unlike HIV_REJO.c_, a single mutation at site D279 was sufficient to mediate complete escape from VRC07 with a modest fitness cost (Figure 3D), in line with the inability of AAV8-VRC07 to suppress HIV_JR-CSF_.

Next, we evaluated the escape paths following AAV8-N6 treatment, finding selection for a single nucleotide change to yield a A281D mutation (Figure S6H), along with loss of a PNGs at site N276 (Figure S6I). The A281D mutation was sufficient to achieve escape (Figure S6J) albeit at a high fitness cost (Figure S6K,L). We also observed that the combination of A281D with N276D resulted in complete escape at a lower fitness cost (Figure S6M).

Finally, escape from AAV8-PGDM1400 treatment resulted in a variety of V2-loop mutations at sites N160 or T162 (Figure S6N), resulting in loss of a PNGs (Figure S6O). All of the mutations evaluated at sites N160 and T162 resulted in escape from PGDM1400 (Figure S6P), with no fitness cost (Figure S6Q,R). The resulting escapability map recapitulates that of HIV_REJO.c_ (Figure S6S).

Considering the lack of viral suppression observed for some VIT-treated HIV-infected mice, we evaluated the *in vitro* neutralizing activity of sera from a subset of these animals (Figure S7A-H). As expected, sera from all mice expressing bNAbs were able to neutralize both HIV_REJO.c_ and HIV_JR-CSF_ with an IC_50_ value similar to purified proteins. No neutralization activity was seen in sera from control mice. Given prior reports of antiviral activity of bNAbs following passive transfer in HIV-infected humanized mice^31^, we explored whether the kinetics of vectored antibody expression played a role in the lack of antiviral activity seen for N6 and PGDM1400 in our studies. We performed passive transfer of either a control antibody (2A10, a malaria-specific antibody), N6, or PGDM1400 in HIV_JR-CSF_ infected humanized mice (Figure S7I-L and Table S3). As expected, high antibody concentrations were detected starting as early as one week post-infusion, and remained elevated throughout the course of weekly administrations (Figure S7I). Interestingly, we did not observe any change in viral load in any of the mice during the length of the experiment (Figure S7K-L). Despite this, deep sequencing of viral envelopes revealed the selection of escape mutations that largely mirrored those seen for AAV-delivered bNAbs (Figure 7M,N). These results suggest that differences in antibody expression kinetics do not explain the lack of activity seen in our studies. Instead, it is possible that HIV infection in the BLT humanized mouse model may be particularly challenging to suppress relative to other models^31^.

### Increasing HIV_REJO.c_ mutation resistance benefit enables complete escape from VRC07

Our experiments demonstrate that HIV_REJO.c_ and HIV_JR-CSF_ exhibit differences in escape paths under VRC07 selection. However, these studies could not discern whether the differences in Env sequence or the inherent fitness differences between these isolates contributed to escapability. Notably, in the context of the HIV_REJO.c_ Env, the D279A mutation only resulted in a partial escape, whereas for HIV_JR-CSF_, the same mutation resulted in complete escape. To evaluate whether this could be explained by differences in the diversification of these viruses in humanized mice, relative to those seen in humans, we compared the baseline frequency of each amino acid at each position in env for both isolates to the frequencies observed in the corresponding position in the LANL database. We found that amino acids that were at a high frequency in HIV_REJO.c_ and HIV_JR-CSF_ also tended to have a higher frequency in the LANL database (Figure S8A), suggesting that both isolates generate diverse quasispecies that are consistent with clinical infections. To determine whether the fitness cost of escape mutations observed in both viruses were consistent with conservation in the LANL database, we directly compared the LANL amino acid frequency of each escape mutation to their observed fitness cost in HIV_REJO.c_ and HIV_JR-CSF_ backbones, and found a strong and significant negative correlation (Figure S8B), confirming that high fitness cost mutations in both isolates are also rarely seen in the LANL database.

To dissect the contribution of the resistance benefit to escape VRC07, we engineered chimeric IMCs of HIV_REJO.c_ containing the swaps of amino acid sequences comprising the D-loop (HIV_RD_), the V5-loop (HIV_RV_) or both loops (HIV_RDV_) from HIV_JR-CSF_ (Figure 3E, Table S4). Each of the three chimeric viruses were neutralized by VRC07 with a similar IC_50_ to those of the original isolates (Figure 3F). Importantly, both D-loop chimeras, HIV_RD_ and HIV_RDV_, exhibited a modest increase in fitness cost, whereas the V5-loop-only chimera, HIV_RV_, showed no difference in fitness relative to the HIV_REJO.c_ strain (Figure S9A,B).

Following infection of BLT humanized mice with each chimeric strain, AAV8-VRC07 administration resulted in a geometric mean steady-state plasma antibody concentration of approximately 50 μg/mL for all the groups (Figure S9C). This led to suppression in a subset of HIV_REJO.c_-infected mice, whereas all HIV_JR-CSF_-infected mice escaped (Figure S9D,E). A similar proportion of mice infected with HIV_RD_ and HIV_RDV_ were suppressed by VRC07, as seen for HIV_REJO.c_ (Figure S9F,G). In contrast, all mice infected with HIV_RV_ exhibited rapid escape from VRC07, as seen for HIV_JR-CSF_ (Figure 3G, Figure S9H,I).

We focused on the HIV_RV_ chimera to understand why this swap resulted in a suppression outcome similar to that of HIV_JR-CSF_. In contrast to HIV_REJO.c_, mutations arising in HIV_RV_-infected animals receiving AAV8-VRC07 were confined to the D-loop (Figure S9J-N). HIV_RV_ mutants harboring single amino acid changes at D279 were completely escaped from VRC07, albeit with substantial fitness costs (Figure S9O-Q).

Taken together, these data suggest that the sequence context of the HIV envelope directly influences the path taken to escape from VRC07 (Figure 3H). The escapability map reveals that D279 mutations result in complete escape for HIV_RV_, in contrast to HIV_REJO.c_. Finally, the Escape Barrier score for HIV_JR-CSF_ escaping from all bNAbs and HIV_RV_ escape from VRC07 was calculated as before (Figure S10A-D). For HIV_JR-CSF_, VRC07 exhibited the lowest score; whereas HIV_RV_ score was between HIV_REJO.c_ and HIV_JR-CSF_ (Figure 3I, Figure S10E,F).

### Increasing HIV_REJO.c_ mutation fitness cost results in complete suppression by VRC07

Given the importance of sequence context in defining the resistance benefit of mutations along the evolutionary escape path, we sought to evaluate the impact of mutation fitness cost on escape. To this end, we sought to alter the fitness of VRC07-escape mutants without affecting their resistance benefits. Given the previously described reduction in viral fitness following escape from antiretroviral therapy (ART) drug regimens^66^, we tested the ability of the Polymerase (Pol) mutations to impact the fitness of VRC07 escape mutations. We engineered Pol M184I and M184V mutations into the original HIV_REJO.c_ IMC vector and detected no changes in neutralization sensitivity to VRC07 or viral fitness (Figure S11A,B). We then evaluated both the fitness cost and resistance benefit of the previously identified VRC07-HIV_REJO.c_ escape mutations (i.e., N276D, D279A, N460D, and their combinations) in the context of the Pol M184V mutation. While we observed no changes in neutralization sensitivity (Figure 2B, Figure S11C, Table S1), we observed significant increases in the fitness cost for escape mutants containing Pol M184V relative to WT Pol (Fig 4A, Figure S11D).

**Figure 4.**
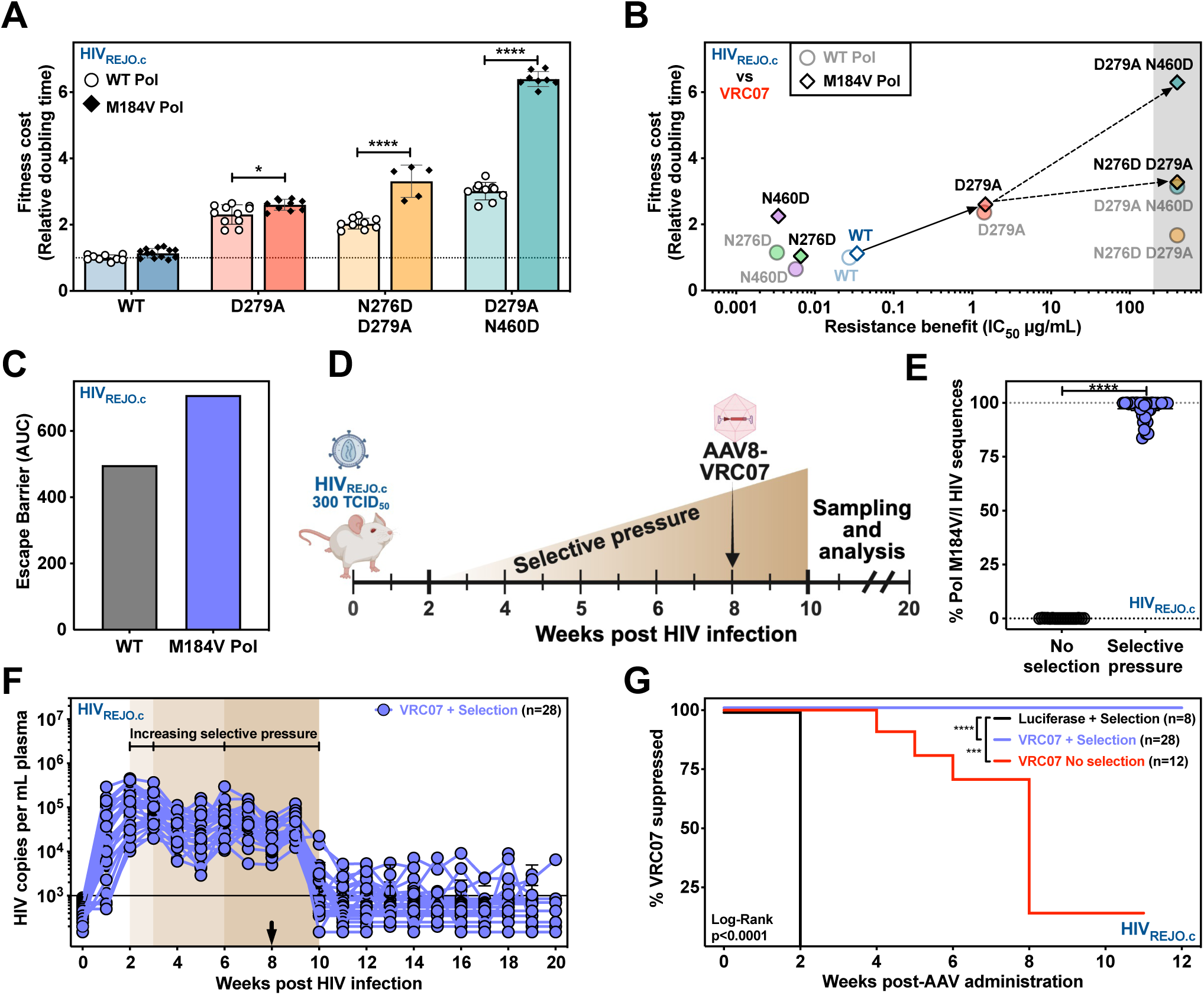
Increasing the fitness cost of HIV_REJO.c_ escape mutations enhances the efficacy of vectored VRC07. **(A)** The fitness of HIV_REJO.c_-VRC07 escape mutations with or without the Pol M184V mutation were evaluated using *QuickFit*. Data are presented as mean ± SD. Statistical differences were assessed by a Two-Way ANOVA, with a Šidák *post hoc* test to correct for multiple comparisons (*: p<0.05; ****: p<0.0001). See also Figure S9. **(B)** Escapability map of HIV_REJO.c_-VRC07 escape mutants with or without the Pol M184V mutation. Solid symbols (diamonds) represent envelope mutations on the Pol M184V mutant background, while lighter symbols (circles) represent the data for envelope mutations on the WT Pol background. Shaded areas represent the limits of detection. Solid arrows represent the likely path taken with dashed arrows representing secondary steps to escape. See also Figure S9.**(C)** Empirical Escape Barrier score for HIV_REJO.c_ escape from VRC07 as compared to the theoretical Escape Barrier score for HIV_REJO.c-PolM184V_ escape from VRC07. **(D)** Experimental setup to determine the impact of Pol M184 mutations on HIV_REJO.c_ escape from VRC07. Humanized mice were infected with HIV_REJO.c_ and then treated with a sub-optimal ART starting 2 weeks after infection, to select and maintain Pol M184 mutants. At week 8, 5×10^11^ GC of AAV8-VRC07 was injected IM, and the ART regimen was stopped at week 10. Mice were followed for 6 months, and blood samples were collected weekly. **(E)** Percentage of Pol M184V/I mutations determined from viral sequences isolated from the plasma of HIV_REJO.c_-infected mice at week 8, prior to AAV8-VRC07 administration. Statistical differences were assessed by an unpaired two-tailed Student’s t-test (****: p<0.0001). Data are presented as mean ± S.E.M. **(F)** HIV_REJO.c_ viral load in plasma of AAV8-VRC07 treated mice. Black arrows denote vector administration. Each colored line depicts an individual mouse. The qPCR lower limit of detection was 1 genome copy per µL of plasma and 5µL were used in the reaction (solid line). Data are presented as mean ± S.E.M. **(G)** Kaplan-Meier plot of viral suppression in humanized mice infected with HIV_REJO.c_ with or without ART selection and with or without AAV8-VRC07 administration. The total model significance (p<0.0001) and pairwise comparisons against the Luciferase + Selection control (***: p<0.001; ****: p<0.0001) were assessed independently with Log-rank (Mantel-Cox) tests. See also Figure S9.

The net effect of this was a shift in the escapability map for HIV_REJO.c_ with the Pol M184V mutation, such that complete escape through a second envelope mutation incurred substantially higher fitness costs than for the WT Pol strain (Figure 4B). The Escape Barrier score was calculated for Pol M184V mutants assuming the same relative escape paths frequencies as seen for the WT Pol, resulting in a 43% increase (Figure 4C), suggesting that selection of the Pol M184 mutations could improve VRC07 suppression of HIV_REJO.c_.

To evaluate the predicted reduction in HIV_REJO.c_ escape from VRC07, we intentionally drove the emergence of Pol mutations through sub-optimal dosing of ART in humanized mice. BLT mice were infected with HIV_REJO.c_ and over the course of 6 weeks, we titrated the dose of ART drug in their food, from 1% to 100% of the standard human equivalent dose. Eight weeks after infection, either AAV8-VRC07 or AAV8-Luciferase were administered. Two weeks after AAV administration, the ART treatment was interrupted (Figure 4D, Table S5). Importantly, at the time of AAV administration, no meaningful differences in viral loads were seen between the mice that had received escalating antiretroviral drugs relative to control mice (Figure S11E). As expected, HIV *pol* genes sequenced from mice receiving the suboptimal ART dosing displayed a near-complete prevalence of M184V/I at the time of VRC07 administration (Figure 4E).

As anticipated, ART drug treatment had no impact on AAV8-driven expression of VRC07 (Figure S11F). ART-treated mice receiving AAV8-Luciferase maintained steady viral loads, with minor dips at each escalation of drug concentration (Figure 4F). Administration of AAV8-VRC07 without ART treatment resulted in only partial suppression of HIV, as observed in our prior experiments (Figure S11G). Additionally, the post-escape viral load for the Luciferase-treated mice showed no difference irrespective of the presence or absence of the selective pressure (Figure S11H). However, administration of AAV8-VRC07 to ART-treated mice resulted in complete suppression (Figure 4F,G), suggesting that the additional fitness costs were specific to the VRC07 escape mutations, as predicted in our theoretical escape map and Escape Barrier scores. Taken together, these results suggest that by decreasing the overall fitness of HIV_REJO.c_, and therefore modulating the fitness cost axis of the escapability maps, resistance to VRC07 becomes significantly more difficult.

## Discussion

Antibody immunotherapy to prevent or treat HIV infection is currently being evaluated in numerous clinical trials. While bNAb combinations have shown promise^42,67–69^, individual bNAbs have demonstrated limited activity in viremic PLWH due to their half-life and the emergence of escape mutations^34,35,37,39,70,71^. Here we show that vectored delivery of a single bNAb, well-matched to the infecting strain, is capable of suppressing viremia in approximately half of HIV-infected humanized mice. Notably, this efficacy was not correlated with neutralization potency determined *in vitro* against the infecting viral stock or breadth metrics derived from global pseudovirus panels^17,62,63^. Rather, our results suggest that the *in vivo* efficacy of this treatment is best explained by the bNAb-specific dynamics of viral escape. By using the BLT humanized mouse platform and viral stocks derived from IMCs, we performed dozens of independent infections with a specific virus and AAV8-bNAb treatment. This led to the reproducible characterization of the most commonly traversed escape paths by HIV, demonstrating surprising conservation of viral escape from specific bNAbs.

Of the two CD4bs-targeting antibodies evaluated in this study, N6 was broader, neutralizing 96% of a 181-strain global panel with an IC_50_ of less than 1 μg/mL^62^. Despite its outstanding potency and breadth, we found that a single base pair change in the D-loop of HIV_REJO.c_ or HIV_JR-CSF_ yielded resistance to neutralization that was above the steady-state concentrations achieved in our experiments. Remarkably, this A281 mutation has also been previously reported for antibodies from the VRC01-class family, suggesting conserved escape paths for these antibodies^72,73^. Despite a lack of viral suppression by AAV8-N6, we observed that mice with higher antibody concentrations exhibited lower post-escape viral loads. As such, a higher steady-state concentration of N6 would likely result in improved viral suppression; however, expression was lower relative to other bNAbs despite identical vector dosage, consistent with previous reports of short N6 half-life following passive transfer^74,75^.

PGDM1400 recognizes the HIV envelope V1/V2 trimer apex and neutralizes with a median IC_50_ of 0.003 μg/mL against a 77-virus panel^63^. A single-point mutation in the V2-loop (N160 or D167) was enough for both HIV isolates to acquire complete resistance to PGDM1400, which has been previously observed for trimer apex-targeting antibodies^76^. The low fitness cost of these mutations is reflected in their high pre-existing frequency in control mice and PLWH (Table S6) (https://www.hiv.lanl.gov/). In a clinical trial using PGDM1400, multiple sequences isolated from rebound participants exhibited a loss of the N160 glycan, suggesting that glycan-based escape is a major driver of bNAb resistance in humans^68^.

VRC07 neutralizes 83% of a panel of 179 strains at less than 1 μg/mL, with a geometric mean IC_50_ of 0.11 μg/mL^17^. In contrast to N6 and PGDM1400, complete escape of HIV_REJO.c_ from VRC07 required a two-step evolutionary path. The mutations we report here include an essential D-loop mutation at D279, resulting in partial resistance with moderate fitness cost, and a second mutation in either the D- or V5-loops which confers complete resistance and further modulates the fitness of the virus. Notably, the tolerability of D- and V5-loop mutations identified through deep mutational scanning varied significantly across strains, in line with our isolate-specific fitness cost data^77^. Although it required two steps, HIV_REJO.c_ escape from VRC07 occurred in 44% of mice across our study. Interestingly, the N276D D279A escape path was observed less often than the D279A N460D escape path, despite having a lower fitness cost. This may be a result of the high frequency of pre-existing N460D (Table S6) and the large distance between these two mutations, which is known to increase the likelihood of recombination^78^. Of note, D279A, along with the paired mutations at N276 and N460, have been previously reported as escape mutations from VRC01-class antibodies in humans^36,73,79^. Finally, HIV_REJO.c_ escapes that did not acquire D279A had a significant delay in time-to-escape, suggesting that blocking a primary escape path can result in the emergence of rarer and more costly alternatives. The differences in escape paths seen for VRC07 suggest that the timing of escape, and particularly how early these mutations emerge, plays a significant role in the therapeutic success of VIT, and future analyses should try to dissect this more in depth.

The AMP trials have identified the predicted serum neutralization 80% inhibitory dilution titer (PT_80_) as a robust predictor of protection in humans and NHPs^24,80^. This biomarker is defined as the bNAb serum concentration over the *in vitro* IC_80_ value against a given HIV strain. A steady-state PT_80_ of 200 or higher correlated with a prevention efficacy of 90%^24^. Despite reaching average PT_80_ values against HIV_REJO.c_ of over 500 for VRC07 and over 400 for PGDM1400 over the four week period of increasing expression, we did not see efficacy against established infections (Figure S12). Interestingly, there were no differences in PT_80_ values between HIV_REJO.c_ suppressed or escaped mice, demonstrating that in our model, potency- and breadth-derived parameters were insufficient to predict the *in vivo* efficacy of bNAbs.

Our results show that escape from CD4bs-directed bNAbs is more likely to impact viral fitness as compared to bNAbs targeting a variable loop, likely due to the CD4-binding requirement during viral infection^36,81^. In a clinical trial evaluating 3BNC117, another CD4bs-targeting bNAb, sequences from three out of eight participants remained sensitive to the bNAb following viral rebound after antibody serum decay^34^. Additionally, patients undergoing ART interruption following 3BNC117 administration remained suppressed until antibody levels declined, suggesting a lack of pre-existing escape variants in the latent viral reservoir^72^. Recent characterization of escape from eCD4-Ig further demonstrates the difficulty of viral escape from CD4bs-targeted therapeutics^81^. Collectively, these findings highlight the advantage of targeting high fitness-cost epitopes.

In this study we project the fitness landscape of each bNAb-strain onto two-dimensional escapability maps by comparing both the fitness cost and resistance benefit of mutations for each major escape path. Using chimeric strains, we modulated the neutralization benefit of escape mutations as represented by the X-axis of the escapability maps. Despite the same neutralization sensitivity to VRC07 and no difference in fitness compared to HIV_REJO.c_, we found that the HIV_RV_ chimera exhibited complete and rapid escape from vectored VRC07, like HIV_JR-CSF_, highlighting the importance of sequence context on escape mutations^77^. In this study, we utilized two different HIV isolates and found that the diversity of the quasispecies in infected humanized mice was similar to the sequence diversity seen in the LANL database, as reported by other studies that evaluated the conservation of mutations cross-sectionally^82^. Importantly, this conservation also correlated with the fitness cost of the mutations, showing that more costly mutations were less represented in the LANL database. The relevance of the sequence context and diversification could have implications for studies performed in NHPs, given the chimeric nature of SHIV^83^.

Using M184 Pol mutations, we modulated the fitness cost of escape mutations, as represented by the Y-axis of the escapability maps. We found that M184 Pol viruses harboring VRC07 escape mutations maintained their susceptibility to VRC07 but grew more slowly than WT Pol strains *in vitro*. We predicted that this shift in fitness cost would result in a more challenging escape path from the antibody *in vivo*. Consistent with this hypothesis, AAV8-VRC07 administration to HIV_REJO.c_-infected animals harboring these Pol mutations resulted in sustained viral suppression. Together, these experiments demonstrate that the two escapability map dimensions of resistance benefit and fitness cost drive the efficacy of AAV-bNAb therapy *in vivo*.

The BLT humanized mouse model enables the systematic comparison of bNAb therapeutic efficacy and identification of escape paths taken by HIV isolates. Despite their lack of endogenous humoral immune responses, here we report the same VRC07, N6, and PGDM1400 escape mutations seen in NHPs and humans, with the benefit of a shorter time frame and lower cost^84^. Given their diverse and continuously evolving viral population, as previously reported^31^ and seen here (Figure S7), HIV-infected humanized mice represent an excellent model to evaluate bNAb escape *in vivo*.

The concept of fitness landscapes has been frequently used to understand HIV evolution, including the dynamics of intrapatient adaptation^85^ and antiretroviral escape^86^. Within this framework, it is well known that a fitness cost is typically accrued during escape from cytotoxic T lymphocytes^87–89^, antiretroviral drugs^90–93^, or antibodies^94^. Indeed, modern cART leverages this understanding to maximize their clinical efficacy^95^. Our findings suggest that bNAb escape paths are predictable and that maximizing the cost of escape *in vivo* may be the key to improving their clinical efficacy. Whether analogous improvements in the clinical efficacy of bNAbs could be achieved by reducing the fitness of viruses replicating in patients who have failed ART regimens remains to be determined. Recent efforts have focused on antibody combinations that target independent sites of vulnerability, however, future bNAb combinations with orthogonal, high cost, escape paths should be tested, as combinations whose escape mutations result in additive fitness costs may improve the therapeutic efficacy of antibody-based interventions.

## Limitations of the study

Despite our best effort to be comprehensive, our study has a number of limitations. First, as previously stated, differences in bNAb expression levels influence the resulting escape paths. Future studies could focus on dissecting the influence of bNAb expression level on escape path selection. In addition, AAV-mediated antibody delivery starts with lower serum levels of bNAb before achieving a high steady-state concentration, whereas passive transfer results in high initial concentration, which then declines over time. This difference could result in divergent outcomes, as passive transfer applies a stronger initial selection pressure, which may alter viral escape dynamics. However, subclinical viral replication has been reported as passively transferred antibody decays, even when still above the currently accepted protective levels^96^. Moreover, at steady-state, the mutations we identified were the same as those which ultimately emerged during passive transfer studies. Second, we report the results of only three antibodies targeting two distinct sites of vulnerability against two clade B isolates of HIV. However, a plethora of bNAbs have been described targeting five sites of vulnerability and many diverse HIV isolates exist ^97^. Whether our findings would extend to other bNAbs and HIV clades remains to be determined. However, several reports show that mutations and evolutionary paths are conserved and convergent across different isolates and clades^98–100^. Third, our study focuses on escape mutations present at the endpoint of each independent experiment, therefore, the dynamic events occurring earlier and over the course of escape still need to be elucidated. Fourth, HIV infection in BLT humanized mice could not be assessed beyond six months due to declining health of animals from this model, which may limit the opportunities for escape mutations to arise. Fifth, humanized mice have a relatively small blood volume and lack secondary lymphoid organs that may not completely recapitulate HIV infection in humans^101^. Moreover, BLT mice elicit a poor humoral immune response, and therefore lack this selective pressure^102^, which may influence the selection of escape paths. In the case of PGDM1400, it has been shown that the polyclonal humoral response elicited in humans is reactive against the various glycans on the Env protein^103^. Viremic patients receiving individual bNAbs have exhibited more substantial drops in viral load than what was seen in our model^34,35,41,70^. Additionally, there have been reports of synergy between passively transferred bNAbs and endogenous antibody responses in patients^104^, which are unlikely to occur in humanized mice. As such, bNAb-based suppression of HIV in humanized mice may represent a high bar for therapeutic efficacy given that suppression is completely dependent on the administered bNAb. Finally, measurements of viral growth rates were performed *in vitro* with PBMCs from a single donor. Whether the absolute growth rate of these mutants across genetically distinct patients varies remains untested, previous studies suggest that their relative fitness would remain unchanged^65^.

## Resources availability

### Lead contact

Further information and requests for resources and reagents should be directed to and will be fulfilled by the lead contact, Dr. Alejandro Balazs.

### Materials availability

All genetic constructs and unique biological materials generated in this study are available from the lead contact without restriction upon request.

### Data and code availability

A summary of the sequencing analyzed data is available online at https://github.com/Balazs-Lab/Escapability. Raw sequencing files reported in this paper will be shared by the lead contact upon request. All other data are available in the main text or as part of the supplementary data and tables. Source data referencing each figure are also provided for this study as supplementary material. Codes used for the sequencing analyses in this study, along with instructions to run them, can be found in the following link: https://github.com/Balazs-Lab/Escapability

## Acknowledgements

We wish to thank Dennis Burton (Scripps) for providing antibody proteins, Dan Barouch for providing anti-PGDM1400 idiotype antibody protein. We also thank K. L. Clayton and D. T. Claiborne for advice on experimental design and the manuscript. We thank the Ragon Institute Protein Core for providing antibody proteins. The following reagents were obtained through the NIH HIV Reagent Program, Division of AIDS, NIAID, NIH: HIV-1, Strain JR-CSF Infectious Molecular Clone (pYK-JRCSF), ARP-2708, contributed by Dr. Irvin S. Y. Chen and Dr. Yoshio Koyanagi; ARP-11746 (pREJO.c/2864), contributed by Dr. John Kappes and Dr. Christina Ochsenbauer; TZM-bl Cells, ARP-8129, contributed by Dr. John C. Kappes, Dr. Xiaoyun Wu and Tranzyme Inc. N.M.S.G. was supported by the Executive Committee on Research (ECOR) of the Mass General Hospital (MGH) Fund for Medical Discovery (FMD) Fundamental Research Fellowship Award. A.D.N. was supported by the Ruth L. Kirschstein National Research Service Award (NRSA) Postdoctoral Training Program Award T32AI007245. C.E.D. was supported by the NRSA Individual Postdoctoral Fellowship 1F32AI125096-01A1. J.M.B. was supported by the NSRA Predoctoral Award 1F31AI131747-01A1. A.B.B. was supported by the National Institute for Allergy and Infectious Disease Career Transition Award K22AI102769, Research Grants R01AI174875, R01AI174276, the National Institutes for Drug Abuse (NIDA) Avenir New Innovator Award DP2DA040254, the NIDA Avant-Garde Award 1DP1DA060607, CDC subcontract 200-2016-91773-T.O.2, the MGH Transformative Scholars Program as well as funding from the Charles H. Hood Foundation. This independent research was supported by the Gilead Sciences Research Scholars Program in HIV.

## Author contributions

C.E.D. and A.B.B. designed the experiments. N.M.S.G., A.D.N., Y.C., C.E.D., C.L.B., S.W.M., Y.E.S.A., E.C.L., M.L.S., D.P., A.L., M.J.D., S.B.Y., B.M., and C.B.B. carried out experiments and analyzed data. J.M.B., A.P., X.C., C.L., R.A.K., J.R.M. and D.L. offered suggestions for experiments and provided key materials. S.T. and V.D.V. provided humanized mice. N.M.S.G., A.D.N, Y.C., C.E.D. and A.B.B. wrote the paper with contributions from all authors. The authors of this study fulfill all the authorship criteria required by Cell Press. Roles and responsibilities were agreed among the authors ahead of the research within reason and changes on these roles were previously agreed by the authors. Declaration of interests A.B.B. is a founder of Cure Systems LLC. All other authors declare no conflicts of interest.

## Supplemental information

Document S1. Figures S1–S12 and Tables S1-S6.

Document S2. Excel file containing all source data referring to all the figures.

Document S3. Excel file presenting all the mice, their treatment, antibody concentration reported at escape, and weeks of suppression (or to escape).

## Methods

### BLT Humanized Mice

BLT humanized mice were generated by the Human Immune System Mouse Program at the Ragon Institute of MGH, MIT, and Harvard. Briefly, 6- to 8-week-old female NSG mice were transplanted with human liver and thymus tissue under the kidney capsule and injected intravenously with 100,000 CD34^+^ cells isolated from liver tissue by AutoMACS (Miltenyi Biotec, Cat#130-100-453). Mice were rested for 10 weeks after surgery to allow for recovery and engraftment. All experiments were done with approval from the Institutional Animal Care and Use Committee of the MGH and conducted in accordance with the guidelines and regulations of the American Association for the Accreditation of Laboratory Animal Care.

### HIV production

HIV was produced by transient transfection of HEK 293T/17 cells (ATCC, Cat#CRL-11268 - RRID:CVCL_1926) using 25K MW Linear Polyethyleneimine (PEI, Polysciences Inc., Cat#23966) maintained in DMEM medium (Corning, Cat# 10-013-CV) supplemented with 10% fetal bovine serum (FBS - VWR, Cat#89510-186), and 1% penicillin–streptomycin mix (Corning, Cat#30-002-CI) with infectious molecular clone (IMC) plasmids encoding for HIV_REJO.c_ or HIV_JR-CSF_ (AIDS Reagent Program NIH - BEI Resources Cat#ARP-11746 and #ARP-2708, respectively) or IMC plasmids containing the indicated mutations. After 48 hours, culture supernatants were collected, filtered through a 0.45-m filter, and titered using either an HIV-1 p24 antigen capture assay (AIDS and Cancer Virus Program, Leidos Biomedical Research, Inc., Frederick National Laboratory for Cancer Research) or a 50% tissue culture infective dose (TCID_50_) assay on TZM-bl cells (BEI Resources, Cat#ARP-8129 - RRID:CVCL_B478). TCID_50_ was calculated using the Spearman-Karber formula^105^.

### Humanized mouse HIV infection

Prior to HIV infection, blood samples were obtained from mice and subjected to flow cytometry to determine the baseline CD3^+^ (BioLegend, Cat#300434 - RRID:AB_10962690), CD4^+^ (BioLegend, Cat#300534 - RRID: AB_2563791), and CD8^+^ (BioLegend, Cat#300554 - RRID: AB_2564382) T cells engraftment levels. The next day, mice were intravenously infected with either 10ng of p24 or 300 TCID_50_ of HIV_REJO.c_ or HIV_JR-CSF_ diluted in PBS (Corning, Cat#21-031-CV) to a volume of 50 ⎧L. Blood was collected weekly to determine viral loads and bNAb levels in serum.

### AAVs vector production, quantification, validation, and administration

AAV8 vectors encoding either Luciferase, 2A10, or bNAbs were produced and validated as previously described^12^. Briefly, HEK293T/17 cells were co-transfected with the AAV backbone vector and helper vectors pHELP and pAAV 2/8 SEED using PEI. AAVs were collected over a five day period following transfection, filtered through a 0.22 µm filter (Corning, Cat#431097), and fresh media was gently added back to the cells each time. After collection, the virus was Polyethylene glycol 8,000 (PEG, VWR, Cat#JTU222-9) precipitated on ice O.N., and then pelleted at 8,000 x g for 30 min. Pellets were re-suspended in cesium chloride, split evenly into two Quick-Seal tubes (Beckman, Cat#342413) and centrifuged at 330,000 x g at 20 °C for 24 h (Beckman Coulter, Optima LE-80K, 70Ti rotor). AAV-containing fractions were determined with a refractometer, with refractive indexes between 1.3755 and 1.3655 considered positive. These were then diluted into 15l mL of Test Formulation Buffer 2 (TFB2, 100 mM sodium citrate (VWR, Cat#Cat#6132-04-3), 10 mM Tris, pH 8.00 (Fisher Scientific, Cat#S1519-500GM)), loaded in a 100 kDa MWCO centrifugal filter (Millipore, Cat#UFC910024) and centrifuged at 500 x g at 4 °C until 1 mL remained in the filter. This wash was repeated twice. Final retentate was aliquoted and stored at −80 °C. Purified AAVs were quantified by qPCR using the PerfeCTa SYBR Green SuperMix, Low ROX (Quanta Biosciences, Cat#95056-500) and primers designed against the CMV enhancer (AACGCCAATAGGGACTTTCC and GGGCGTACTTGGCATATGAT). Samples were run in duplicate on an QuantStudio 12K Flex (Applied Biosystems). To validate the functional activity of each lot, *in vitro* transduction assays were performed in HEK293T/17 cells. Six days after transduction, supernatants were recovered and quantified for total IgG production by ELISA. AAV8 IM injections were performed as previously described^12^. Briefly, aliquots of previously titered viruses were thawed on ice and diluted in PBS to achieve the predetermined dose in a 40 ⎧L volume. A single 40 ⎧L injection was administered into the gastrocnemius muscle of BLT humanized mice with a 28 G insulin syringe.

### Antibody quantification by ELISA

Plasma was used to determine bNAb concentrations. For detection of gp120-binding IgG, ELISA plates were coated with 2 ⎧g/mL of HIV gp120 protein (Novus, Cat#NBP1-76371 or Immune Technology, Cat#IT-001-0025p) per well for 1h at room temperature. For detection of PGDM1400, ELISA plates were coated with BG505 SOSIP (provided by the Vaccine Research Center) at 5 ⎧g/mL or a PGDM1400-specific idiotype (provided by Dan Barouch) at 1 ⎧g/mL for 2hr at room temperature. Plates were blocked with 1% BSA (LGC Clinical Diagnostics, Cat#50-61-00) in Tris-buffered saline (TBS - Thermo Scientific, Cat#AAJ75892AE) overnight at 4°C. Samples were incubated in TBS plus Tween 20 (Thermo Scientific, Cat#BP-337-100) containing 1% BSA for 1h at room temperature before incubation with 1:2,500 to 1:10,000-diluted horseradish peroxidase (HRP)-conjugated goat anti-human IgG-Fc antibody (Bethyl, Cat#A80-104A - RRID: AB_67064) for 30 min at room temperature. Samples were detected by the TMB Microwell Peroxidase substrate system (SeraCare, Cat#50-76-00). A standard curve was generated using purified VRC07, N6, or PGDM1400 (provided by the Vaccine Research Center) as appropriate for the sample.

### Viral load test by quantitative RT-PCR (RT-qPCR)

Viral RNA was extracted from plasma samples using the QIAamp viral RNA mini kit (Qiagen, Cat#52906)). Each RNA sample was treated with 2 U of Turbo DNase (Fischer Scientific, Cat#AM2239) at 37°C for 30 min followed by heat inactivation at 75°C for 15 min. 10 ⎧L of the treated RNA were used in a 20 ⎧L RT-qPCR reaction with the qScript XLT one-step RT-qPCR Tough Mix, low ROX mix (Quanta Biosciences, Cat#95134-500), a TaqMan probe (5’-/56- FAM/CCCACCAAC/ZEN/AGGCGGCCTTAACTG/3IABkFQ/-3’) (IDT) and primers designed targeting the Pol gene of HIV_REJO.c_ (CAATGGCCCCAATTTCATCA and GAATGCCGAATTCCTGCTTGA) or HIV_JR-CSF_ (CAATGGCAGCAATTTCACCA and GAATGCCAAATTCCTGCTTGA). Samples were run in triplicate on a QuantStudio 12K Flex (Applied Biosystems). The following cycling conditions were used: 50°C for 10 min, 95°C for 3 min followed by 55 cycles of 95°C for 3s and 60°C for 30s. Virus titer was determined by comparison with a standard curve generated using RNA extracted from serially diluted mixture of commercially titered viral stock and pure mouse serum. The limits of detection were 1,000 copies per mL for all viral strains. For the purpose of generating Kaplan-Meier curves, viral escape was defined as the first week after AAV administration in which viral load did not decrease at least 75% relative to the prior week and was above 10^4^ copies per mL, provided that a subsequent week was also above that value. Curves were analyzed for statistical significance using the Mantel-Cox Log-rank test in Graphpad Prism v10.2.2 (RRID:SCR_002798).

### Illumina deep sequencing and identification of HIV envelope mutants

Viral RNA extracted from blood samples at the conclusion of each study was used to synthesize cDNA using the SuperScript IV Reverse Transcriptase enzyme with a strain-specific 3’ primer for HIV_REJO.c_ (TTGGTACTTGTGATTGCTCCATGTCTCTCC) or HIV_JR-CSF_ (CCCTATCTGTTGCTGGCTCAGCTCGTC). cDNA was subjected to nested PCR amplification with HIV-specific envelope primers that yielded a 2.5-kb fragment. The first-round primers used for amplification from HIV_REJO.c_ were GCAATAGTAGCATTAGTAATAGCAGGAATAATAGCAATAGTTGTGTGG and CTGCTCCCACCCCCTCTG; whereas the primers used for envelope amplification from HIV_JR-CSF_ were GCAATAATTGTGTGGTCCATAGTACTCATAGAATATAGGA and CCCTATCTGTTGCTGGCTCAGCTCGTC. First-round PCR was performed with 1 to 2.5 µL from cDNA reaction using 1x Q5 reaction buffer, 5 mM dNTPs, 0.5 ⎧M of strain specific primers and 0.02U/⎧L of Q5 Hot Start High-Fidelity DNA Polymerase (NEB, Cat#M0493L) in a total reaction volume of 25 ⎧L. PCR conditions for the first round were: 98°C for 30s, followed by 30 cycles of 98°C for 10s, 70°C for 30s, 72°C for 3 min, with a final extension of 72°C for 2 min. 1⎧L of first round PCR product was then used as the template for the second round PCR with identical cycling conditions and PCR mix except for the primers. The second-round primers used for envelope amplification from HIV_REJO.c_ were AAAATAGACAGGTTAATTGATAGAATAAGAGATAGAGCAGAAGACAGTG and TCATTCTTTCCCTTACAGCAGGCCATC; whereas the primers used for envelope amplification from HIV_JR-CSF_ were AAAATAGATAGGTTAATTGATAAAATAAGAGAGAGAGCAGAAGACAG and TCATTCTTTCCCTTACAGTAGACCATCCAGGC. PCR products were gel extracted, diluted to 0.15 ng/⎧L in UV-irradiated water and subjected to Nextera XT Illumina library preparation (Illumina, Cat#FC-131-1096). PCR products were quantified using Qubit and D5000 ScreenTape System (Agilent, Cat#5067-5589). The library pool was denatured with 0.2 N NaOH, diluted to 4 nM, spiked with 10% PhiX to improve sequence heterogeneity and quality and subjected to 2 x 250 or 2 x 300 paired-end sequencing on the Illumina MiSeq (RRID:SCR_016379).

### Envelope Escape Mutant Analysis

Sequencing reads were filtered for quality using fastp (RRID: SCR_016962)^106^ and aligned to a reference sequence specific to the viral strain under analysis using Bowtie2 (RRID: SCR_016368)^107^. The alignments were sorted and indexed using samtools (RRID: SCR_002105)^108^. Amino acid changes were called using a custom codon aware variant caller, written in python (RRID: SCR_008394). For each sample, the total divergence at each amino acid site was determined by summing the frequency of every non-WT amino acid (relative to the IMC strain sequence, meaning that synonymous mutations are not considered divergent). The group amino acid divergence (plotted in the figures) was determined by averaging the amino acid divergence across each sample at every site. Each site was numbered using the HIV_HXB2_ nomenclature^109^. The analysis pipeline was run using snakemake (RRID: SCR_003475)^110^. All the bioinformatics tools used in these analyses are available online at https://github.com/Balazs-Lab/Escapability.

### Viral Diversity and Variant Conservation Analysis

Similar to the envelope escape mutation analysis, sequencing reads were filtered for quality using fastp^106^ and aligned to a reference sequence specific to the viral strain under analysis using Bowtie2^107^. The alignments were sorted and indexed using samtools^108^. SNVs were called using LoFreq^111^. The heterogeneity of each sample was determined using the SNV data to calculate the average Shannon entropy across all sites in the viral envelope, as performed previously^112,113^. The Shannon entropy at each site in the genome was calculated by taking the sum of the frequency of each SNV times the natural log of the SNV frequency (frequency * ln(frequency)), multiplied by negative one. The analysis pipeline was run using snakemake^110^ and is available online at https://github.com/Balazs-Lab/Escapability. For the comparison of viral diversification to the LANL database, the HIV1 FLT 2022 Env Protein Alignment was downloaded from LANL and filtered for subtype B sequences. All positions were numbered based on their HIV_HXB2_ alignment and only positions shared across HIV_REJO.c_, HIV_JR-CSF_, and the LANL database were analyzed. The average frequency of each HIV_REJO.c_ and HIV_JR-CSF_ amino acid was binned by rounding the log_10_ transformed frequency using the MROUND() function in excel. These grouped bins were then used to evaluate the LANL amino acid frequency data, of which the average and standard deviation were calculated using the Excel Pivot Table functions.

### Construction of HIV mutants

Individual mutations were introduced into the parental IMC vectors expressing the molecular clone using overlapping PCR with primers incorporating the desired mutations. After amplification of the *env* or *pol* gene with the mutagenesis primers using KOD Hot Start Master mix (EMD Millipore, Cat#71975-3) or Q5 Hot Start Master Mix (NEB, Cat#M0493L), the PCR product was purified by gel extraction (Promega, Cat#A9282) and cloned by homologous recombination into the appropriate recipient parental backbone vector using the In-Fusion HD cloning kit (Clontech, Cat#639650). The ligation product was transformed into DH5〈 (Zymo, Cat#T3009) or SURE2 (Agilent, Cat#200152) competent cells, and positive clones were full plasmid sequenced.

### *In vitro* neutralization assay

To compare the sensitivity of point-mutant viruses to bNAb antibody neutralization, each mutant was produced by transient transfection of HEK 293T cells as described above and viral supernatants were titered by TCID_50_ on TZM-bl cells. Then, neutralization assays were performed using a TECAN Fluent 780 liquid handler by mixing 20 µL of virus with 20µL of 2.5-fold serial dilutions of each antibody and incubating this mixture at 25°C for 1 h. After the incubation, antibody-virus mixtures were added to previously plated 6,000 TZM-bl cells with 75 ⎧g per mL of DEAE dextran (Sigma, Cat#D9885-10G) and incubated at 37°C for 48 h. Cells were then lysed using luciferin-containing buffer^114^ and Luciferase signal was quantified using a PHERAstar FSX plate reader (BMG LabTech). Percentage of infection was determined by calculating the difference in luminescence between test wells (cells with virus and antibody) and cell control wells (cells only) and dividing this value by the difference between the virus control wells (cells with virus) and the cell control wells. These values were plotted against antibody concentrations and fitted into a four-parameter nonlinear regression to calculate IC_50_ and Hill Slope using GraphPad Prism v10.2.2.

### Determination of viral fitness

*In silico* viral growth curves were generated with growth rates derived from *in vitro* QuickFit assays as described previously^65^. Briefly, commercially acquired human PBMCs (AllCells, Cat#PB004F; Lot #A2857) were thawed and CD4^+^ T cells were isolated using the EasySep™ Human CD4^+^ T Cell Isolation Kit (STEMCELL Technologies, Cat#17952). Naive CD4^+^ T cells were resuspended in complete RPMI 1640 (Corning, Cat#36750) (cRPMI; 10% FBS and 1% penicillin/streptomycin) supplemented with 10 ng/mL recombinant IL-2 (R&D Systems, Cat#202-IL-050) and 4 µg/mL of anti-CD28 antibody (Biolegend, Cat#302934 - RRID: AB_11148949), plated in 24-well plates coated with 2 µg/mL of anti-CD3 antibody (Biolegend, Cat#317326 - RRID: AB_11150592), and incubated at 37°C and 5% CO_2_ for 4 days. Cells were then pooled and incubated for another 4 days before use. Purification and activation efficiency were evaluated by flow cytometry. Previously titered viruses were three-fold serially diluted and then added to 50 ⎧L of activated CD4^+^ (1×10^5^ cells per well) plated in a 96 round-well plate. Viruses and cells were spinoculated at 1,200 RPM for 1 hour at 20°C, and then incubated for 24 hours at 37°C with 5% CO_2_. Cells were washed five times with 200⎧L of cRPMI, resuspended in 200 ⎧L of fresh cRPMI plus 10 ng/mL of IL-2 and finally transferred to a 96 flat-well plate. Plates were incubated at 37°C with 5% CO_2_ for 6 days. 32 ⎧L of supernatant were collected daily and fresh media was added to replace the volume. Collected supernatants were immediately RNA extracted using QuickExtract DNA Extraction Solution (Biosearch Technologies, Cat#QE09050)^65^. Extracted RNA was used to determine viral loads by RT-qPCR as stated above. Viral loads were used to determine growth rates and generate *in silico* growth curves using a half-maximal equation in MonolixSuite 2023R1 (Lixoft).

### Escape Barrier Analysis

Sample specific sequencing data from the Envelope Escape Mutant Analysis was used to generate sample specific escape haplotypes. Sequences from each escaped sample were filtered for a minimum mutation frequency of at least 10%, and then manually classified into an escape path using the decision tree algorithms described in Figure S3 and Figure S8. The area under the curve (AUC) for each escape path was calculated from the IC_50_ values and relative fitness cost data using the trapz function in the pracma^115^ package in R (RRID:SCR_001905)^116^. The calculation began at the WT coordinate for each virus and summed the AUC up to the position of the escaped haplotype. If the escape IC_50_ was greater than 200 µg/mL (the limit of detection), then the X-coordinate for IC_50_ was assigned a value of 200 µg/mL, otherwise the IC_50_ of the escape haplotype was used. For all samples (except the HIV_RV_ escape) only escape paths A and B were used for the Escape Barrier score analysis because they represented at least 50% of the escape paths. The raw AUC value for each path was scaled by the relative proportions of the paths (Relative Fraction value in Figure S4D and S8E). The path scaled AUC scores were then summed to create the Escape Barrier Score. The scripts and coordinate files are available at https://github.com/Balazs-Lab/Escapability.

### Passive transfer experiments

Humanized BLT mice were infected with HIV_JR-CSF_ as described above, and 4 weeks later were passively infused with 25µg per gram of body weight of a control antibody (2A10), N6 or PGDM1400, weekly for 12 weeks. Blood was collected weekly to determine viral loads and bNAb levels in serum.

### *In vivo* antiretroviral selection-pressure of HIV

To select for ART-escaping HIV mutations, individual tablets of emtricitabine (Emtriva^TM^ (FTC), Gilead Sciences) or Tenvir (tenofovir disoproxil fumarate (TDF); Cipla LTD) were crushed into a fine powder and manufactured with TestDiet 5B1Q feed (Modified LabDiet 5058 with 0.12% amoxicillin) into powder. The final concentration of these drugs in the stock food was 2.3% (4500 mg/kg TDF, 3000 mg/kg FTC)^28^. To achieve a comparable human dose (i.e., 200 mg FTC, 300 mg TDF), the Reagan-Shaw formula^117^ was used to translate the dose from human to mouse with the assumption that an average mouse weighs 20 g and a human weighs 60 kg. On average mice ate 2 g of food per day and the powdered ART-food was diluted in normal TestDiet 5B1Q food to achieve ingestion of the corresponding target human dose per day. BLT mice were infected with HIV_REJO.c_ as previously stated and over the course of 6 weeks, we titrated the dose of ART drug in their food, from 1% to 100% of the standard human equivalent dose (1% for weeks 2 and 3, 10% for weeks 4 through 6, 100% until week 10). At week 8, mice were injected with AAV expressing VRC07 or Luciferase as stated above. At week 10, ART treatment was interrupted. Blood samples were collected weekly to evaluate viral loads and antibody expression.

## References

1. Landovitz, R.J., Scott, H., and Deeks, S.G. (2023). Prevention, treatment and cure of HIV infection. Nat Rev Microbiol 21, 657–670. 10.1038/s41579-023-00914-1.

2. Bekker, L.-G., Das, M., Abdool Karim, Q., Ahmed, K., Batting, J., Brumskine, W., Gill, K., Harkoo, I., Jaggernath, M., Kigozi, G., et al. (2024). Twice-Yearly Lenacapavir or Daily F/TAF for HIV Prevention in Cisgender Women. N Engl J Med 391, 1179–1192. 10.1056/NEJMoa2407001.

3. Jewell, B.L., Mudimu, E., Stover, J., Ten Brink, D., Phillips, A.N., Smith, J.A., Martin-Hughes, R., Teng, Y., Glaubius, R., Mahiane, S.G., et al. (2020). Potential effects of disruption to HIV programmes in sub-Saharan Africa caused by COVID-19: results from multiple mathematical models. The Lancet HIV 7, e629–e640. 10.1016/S2352-3018(20)30211-3.

4. McCoy, L.E., and Burton, D.R. (2017). Identification and specificity of broadly neutralizing antibodies against HIV. Immunological reviews 275, 11–20. 10.1111/imr.12484.

5. Sajadi, M.M., Dashti, A., Rikhtegaran Tehrani, Z., Tolbert, W.D., Seaman, M.S., Ouyang, X., Gohain, N., Pazgier, M., Kim, D., Cavet, G., et al. (2018). Identification of Near-Pan-neutralizing Antibodies against HIV-1 by Deconvolution of Plasma Humoral Responses. Cell. 10.1016/j.cell.2018.03.061.

6. deCamp, A., Hraber, P., Bailer, R.T., Seaman, M.S., Ochsenbauer, C., Kappes, J., Gottardo, R., Edlefsen, P., Self, S., Tang, H., et al. (2014). Global Panel of HIV-1 Env Reference Strains for Standardized Assessments of Vaccine-Elicited Neutralizing Antibodies. J Virol 88, 2489–2507. 10.1128/JVI.02853-13.

7. Sok, D., and Burton, D.R. (2018). Recent progress in broadly neutralizing antibodies to HIV. Nat Immunol 19, 1179–1188. 10.1038/s41590-018-0235-7.

8. Wu, X., Zhang, Z., Schramm, C.A., Joyce, M.G., Kwon, Y.D., Zhou, T., Sheng, Z., Zhang, B., O’Dell, S., McKee, K., et al. (2015). Maturation and Diversity of the VRC01-Antibody Lineage over 15 Years of Chronic HIV-1 Infection. Cell 161, 470–485. 10.1016/j.cell.2015.03.004.

9. Diskin, R., Scheid, J.F., Marcovecchio, P.M., West, A.P., Klein, F., Gao, H., Gnanapragasam, P.N.P., Abadir, A., Seaman, M.S., Nussenzweig, M.C., et al. (2011). Increasing the Potency and Breadth of an HIV Antibody by Using Structure-Based Rational Design. Science. 10.1126/science.1213782.

10. Doria-Rose, N.A., Bhiman, J.N., Roark, R.S., Schramm, C.A., Gorman, J., Chuang, G.-Y., Pancera, M., Cale, E.M., Ernandes, M.J., Louder, M.K., et al. (2016). New Member of the V1V2-Directed CAP256-VRC26 Lineage That Shows Increased Breadth and Exceptional Potency. J Virol 90, 76–91. 10.1128/JVI.01791-15.

11. Balazs, A.B., Ouyang, Y., Hong, C.M., Chen, J., Nguyen, S.M., Rao, D.S., An, D.S., and Baltimore, D. (2014). Vectored immunoprophylaxis protects humanized mice from mucosal HIV transmission. Nature Medicine 20, 296–300. 10.1038/nm.3471.

12. Balazs, A.B., Chen, J., Hong, C.M., Rao, D.S., Yang, L., and Baltimore, D. (2011). Antibody-based protection against HIV infection by vectored immunoprophylaxis. Nature 481, 81–84. 10.1038/nature10660.

13. Deruaz, M., Moldt, B., Le, K.M., Power, K.A., Vrbanac, V.D., Tanno, S., Ghebremichael, M.S., Allen, T.M., Tager, A.M., Burton, D.R., et al. (2016). Protection of Humanized Mice From Repeated Intravaginal HIV Challenge by Passive Immunization: A Model for Studying the Efficacy of Neutralizing Antibodies In Vivo. The Journal of Infectious Diseases 214, 612–616. 10.1093/infdis/jiw203.

14. Stoddart, C.A., Galkina, S.A., Joshi, P., Kosikova, G., Long, B.R., Maidji, E., Moreno, M.E., Rivera, J.M., Sanford, U.R., Sloan, B., et al. (2014). Efficacy of broadly neutralizing monoclonal antibody PG16 in HIV-infected humanized mice. Virology 462-463C, 115–125. 10.1016/j.virol.2014.05.036.

15. Veselinovic, M., Neff, C.P., Mulder, L.R., and Akkina, R. (2012). Topical gel formulation of broadly neutralizing anti-HIV-1 monoclonal antibody VRC01 confers protection against HIV-1 vaginal challenge in a humanized mouse model. Virology 432, 505–510. 10.1016/j.virol.2012.06.025.

16. Brady, J.M., Phelps, M., MacDonald, S.W., Lam, E.C., Nitido, A., Parsons, D., Boutros, C.L., Deal, C.E., Garcia-Beltran, W.F., Tanno, S., et al. (2022). Antibody-mediated prevention of vaginal HIV transmission is dictated by IgG subclass in humanized mice. Sci. Transl. Med. 14, eabn9662. 10.1126/scitranslmed.abn9662.

17. Rudicell, R.S., Kwon, Y.D., Ko, S.-Y., Pegu, A., Louder, M.K., Georgiev, I.S., Wu, X., Zhu, J., Boyington, J.C., Chen, X., et al. (2014). Enhanced potency of a broadly neutralizing HIV-1 antibody in vitro improves protection against lentiviral infection in vivo. Journal of Virology 88, 12669–12682. 10.1128/JVI.02213-14.

18. Julg, B., Liu, P.-T., Wagh, K., Fischer, W.M., Abbink, P., Mercado, N.B., Whitney, J.B., Nkolola, J.P., McMahan, K., Tartaglia, L.J., et al. (2017). Protection against a mixed SHIV challenge by a broadly neutralizing antibody cocktail. Science translational medicine 9, eaao4235. 10.1126/scitranslmed.aao4235.

19. Julg, B., Tartaglia, L.J., Keele, B.F., Wagh, K., Pegu, A., Sok, D., Abbink, P., Schmidt, S.D., Wang, K., Chen, X., et al. (2017). Broadly neutralizing antibodies targeting the HIV-1 envelope V2 apex confer protection against a clade C SHIV challenge. Science translational medicine 9, eaal1321. 10.1126/scitranslmed.aal1321.

20. Julg, B., Sok, D., Schmidt, S.D., Abbink, P., Newman, R.M., Broge, T., Linde, C., Nkolola, J., Le, K., Su, D., et al. (2017). Protective Efficacy of Broadly Neutralizing Antibodies with Incomplete Neutralization Activity against Simian-Human Immunodeficiency Virus in Rhesus Monkeys. Journal of Virology 91, e01187–17. 10.1128/JVI.01187-17.

21. Moldt, B., Rakasz, E.G., Schultz, N., Chan-Hui, P.-Y., Swiderek, K., Weisgrau, K.L., Piaskowski, S.M., Bergman, Z., Watkins, D.I., Poignard, P., et al. (2012). Highly potent HIV-specific antibody neutralization in vitro translates into effective protection against mucosal SHIV challenge in vivo. Proceedings of the National Academy of Sciences of the United States of America 109, 18921–18925. 10.1073/pnas.1214785109.

22. Corey, L., Gilbert, P.B., Juraska, M., Montefiori, D.C., Morris, L., Karuna, S.T., Edupuganti, S., Mgodi, N.M., deCamp, A.C., Rudnicki, E., et al. (2021). Two Randomized Trials of Neutralizing Antibodies to Prevent HIV-1 Acquisition. N Engl J Med 384, 1003–1014. 10.1056/NEJMoa2031738.

23. Mayer, K.H., Seaton, K.E., Huang, Y., Grunenberg, N., Isaacs, A., Allen, M., Ledgerwood, J.E., Frank, I., Sobieszczyk, M.E., Baden, L.R., et al. (2017). Safety, pharmacokinetics, and immunological activities of multiple intravenous or subcutaneous doses of an anti-HIV monoclonal antibody, VRC01, administered to HIV-uninfected adults: Results of a phase 1 randomized trial. PLoS medicine 14, e1002435. 10.1371/journal.pmed.1002435.

24. Gilbert, P.B., Huang, Y., deCamp, A.C., Karuna, S., Zhang, Y., Magaret, C.A., Giorgi, E.E., Korber, B., Edlefsen, P.T., Rossenkhan, R., et al. (2022). Neutralization titer biomarker for antibody-mediated prevention of HIV-1 acquisition. Nat Med 28, 1924–1932. 10.1038/s41591-022-01953-6.

25. Badamchi-Zadeh, A., Tartaglia, L.J., Abbink, P., Bricault, C.A., Liu, P.-T., Boyd, M., Kirilova, M., Mercado, N.B., Nanayakkara, O.S., Vrbanac, V.D., et al. (2018). Therapeutic Efficacy of Vectored PGT121 Gene Delivery in HIV-1-Infected Humanized Mice. Journal of Virology 92, 118. 10.1128/JVI.01925-17.

26. Barouch, D.H., Whitney, J.B., Moldt, B., Klein, F., Oliveira, T.Y., Liu, J., Stephenson, K.E., Chang, H.-W., Shekhar, K., Gupta, S., et al. (2013). Therapeutic efficacy of potent neutralizing HIV-1-specific monoclonal antibodies in SHIV-infected rhesus monkeys. Nature, 1–16. 10.1038/nature12744.

27. Borducchi, E.N., Liu, J., Nkolola, J.P., Cadena, A.M., Yu, W.-H., Fischinger, S., Broge, T., Abbink, P., Mercado, N.B., Chandrashekar, A., et al. (2018). Antibody and TLR7 agonist delay viral rebound in SHIV-infected monkeys. Nature 278, 1295. 10.1038/s41586-018-0600-6.

28. Halper-Stromberg, A., Lu, C.-L., Klein, F., Horwitz, J.A., Bournazos, S., Nogueira, L., Eisenreich, T.R., Liu, C., Gazumyan, A., Schaefer, U., et al. (2014). Broadly Neutralizing Antibodies and Viral Inducers Decrease Rebound from HIV-1 Latent Reservoirs in Humanized Mice. Cell, 1–11. 10.1016/j.cell.2014.07.043.

29. Horwitz, J.A., Halper-Stromberg, A., Mouquet, H., Gitlin, A.D., Tretiakova, A., Eisenreich, T.R., Malbec, M., Gravemann, S., Billerbeck, E., Dorner, M., et al. (2013). HIV-1 suppression and durable control by combining single broadly neutralizing antibodies and antiretroviral drugs in humanized mice. Proceedings of the National Academy of Sciences of the United States of America. 10.1073/pnas.1315295110.

30. Julg, B., Pegu, A., Abbink, P., Liu, J., Brinkman, A., Molloy, K., Mojta, S., Chandrashekar, A., Callow, K., Wang, K., et al. (2017). Virological Control by the CD4-Binding Site Antibody N6 in Simian-Human Immunodeficiency Virus-Infected Rhesus Monkeys. Journal of Virology 91, e00498–17. 10.1128/JVI.00498-17.

31. Klein, F., Halper-Stromberg, A., Horwitz, J.A., Gruell, H., Scheid, J.F., Bournazos, S., Mouquet, H., Spatz, L.A., Diskin, R., Abadir, A., et al. (2012). HIV therapy by a combination of broadly neutralizing antibodies in humanized mice. Nature 492, 118–122. 10.1038/nature11604.

32. Shingai, M., Nishimura, Y., Klein, F., Mouquet, H., Donau, O.K., Plishka, R., Buckler-White, A., Seaman, M., Piatak, M., Lifson, J.D., et al. (2013). Antibody-mediated immunotherapy of macaques chronically infected with SHIV suppresses viraemia. Nature, 1–15. 10.1038/nature12746.

33. Bar, K.J., Sneller, M.C., Harrison, L.J., Justement, J.S., Overton, E.T., Petrone, M.E., Salantes, D.B., Seamon, C.A., Scheinfeld, B., Kwan, R.W., et al. (2016). Effect of HIV Antibody VRC01 on Viral Rebound after Treatment Interruption. The New England journal of medicine 375, 2037–2050. 10.1056/NEJMoa1608243.

34. Caskey, M., Klein, F., Lorenzi, J.C.C., Seaman, M.S., West, A.P., Buckley, N., Kremer, G., Nogueira, L., Braunschweig, M., Scheid, J.F., et al. (2015). Viraemia suppressed in HIV-1-infected humans by broadly neutralizing antibody 3BNC117. Nature. 10.1038/nature14411.

35. Caskey, M., Schoofs, T., Gruell, H., Settler, A., Karagounis, T., Kreider, E.F., Murrell, B., Pfeifer, N., Nogueira, L., Oliveira, T.Y., et al. (2017). Antibody 10-1074 suppresses viremia in HIV-1-infected individuals. Nature Medicine 23, 185–191. 10.1038/nm.4268.

36. Lynch, R.M., Wong, P., Tran, L., O’Dell, S., Nason, M.C., Li, Y., Wu, X., and Mascola, J.R. (2015). HIV-1 fitness cost associated with escape from the VRC01 class of CD4 binding site neutralizing antibodies. Journal of Virology 89, 4201–4213. 10.1128/JVI.03608-14.

37. Mehandru, S., Vcelar, B., Wrin, T., Stiegler, G., Joos, B., Mohri, H., Boden, D., Galovich, J., Tenner-Racz, K., Racz, P., et al. (2007). Adjunctive passive immunotherapy in human immunodeficiency virus type 1-infected individuals treated with antiviral therapy during acute and early infection. Journal of Virology 81, 11016–11031. 10.1128/JVI.01340-07.

38. Mendoza, P., Gruell, H., Nogueira, L., Pai, J.A., Butler, A.L., Millard, K., Lehmann, C., Suárez, I., Oliveira, T.Y., Lorenzi, J.C.C., et al. (2018). Combination therapy with anti-HIV-1 antibodies maintains viral suppression. Nature, 1–21. 10.1038/s41586-018-0531-2.

39. Trkola, A., Kuster, H., Rusert, P., Joos, B., Fischer, M., Leemann, C., Manrique, A., Huber, M., Rehr, M., Oxenius, A., et al. (2005). Delay of HIV-1 rebound after cessation of antiretroviral therapy through passive transfer of human neutralizing antibodies. Nature Medicine 11, 615–622. 10.1038/nm1244.

40. Happe, M., Lynch, R.M., Fichtenbaum, C.J., Heath, S.L., Koletar, S.L., Landovitz, R.J., Presti, R.M., Santana-Bagur, J.L., Tressler, R.L., Holman, L.A., et al. (2025). Virologic effects of broadly neutralizing antibodies VRC01LS and VRC07-523LS on chronic HIV-1 infection. JCI Insight 10, e181496. 10.1172/jci.insight.181496.

41. Bar-On, Y., Gruell, H., Schoofs, T., Pai, J.A., Nogueira, L., Butler, A.L., Millard, K., Lehmann, C., rez, I.S. x000E1, Oliveira, T.Y., et al. (2018). Safety and antiviral activity of combination HIV-1 broadly neutralizing antibodies in viremic individuals. Nature Medicine, 1–13. 10.1038/s41591-018-0186-4.

42. Julg, B., Walker-Sperling, V.E.K., Wagh, K., Aid, M., Stephenson, K.E., Zash, R., Liu, J., Nkolola, J.P., Hoyt, A., Castro, M., et al. (2024). Safety and antiviral effect of a triple combination of HIV-1 broadly neutralizing antibodies: a phase 1/2a trial. Nat Med. 10.1038/s41591-024-03247-5.

43. Ledgerwood, J.E., Coates, E.E., Yamshchikov, G., Saunders, J.G., Holman, L., Enama, M.E., DeZure, A., Lynch, R., Gordon, I., Plummer, S., et al. (2015). Safety, Pharmacokinetics, and Neutralization of the Broadly Neutralizing HIV-1 Human Monoclonal Antibody VRC01 in Healthy Adults. Clinical and experimental immunology. 10.1111/cei.12692.

44. Lewis, A.D., Chen, R., Montefiori, D.C., Johnson, P.R., and Clark, K.R. (2002). Generation of Neutralizing Activity against Human Immunodeficiency Virus Type 1 in Serum by Antibody Gene Transfer. Journal of Virology 76, 8769–8775. 10.1128/JVI.76.17.8769-8775.2002.

45. Fang, J., Qian, J.-J., Yi, S., Harding, T.C., Tu, G.H., VanRoey, M., and Jooss, K. (2005). Stable antibody expression at therapeutic levels using the 2A peptide. Nature Biotechnology 23, 584–590. 10.1038/nbt1087.

46. Fuchs, S.P., Martinez-Navio, J.M., Piatak, M., Lifson, J.D., Gao, G., and Desrosiers, R.C. (2015). AAV-Delivered Antibody Mediates Significant Protective Effects against SIVmac239 Challenge in the Absence of Neutralizing Activity. PLoS Pathogens 11, e1005090. 10.1371/journal.ppat.1005090.

47. Gardner, M.R., Kattenhorn, L.M., Kondur, H.R., von Schaewen, M., Dorfman, T., Chiang, J.J., Haworth, K.G., Decker, J.M., Alpert, M.D., Bailey, C.C., et al. (2015). AAV-expressed eCD4-Ig provides durable protection from multiple SHIV challenges. Nature, 1–16. 10.1038/nature14264.

48. Brady, J.M., Baltimore, D., and Balazs, A.B. (2017). Antibody gene transfer with adeno-associated viral vectors as a method for HIV prevention. Immunological reviews 275, 324–333. 10.1111/imr.12478.

49. Saunders, K.O., Wang, L., Joyce, M.G., Yang, Z.-Y., Balazs, A.B., Cheng, C., Ko, S.-Y., Kong, W.-P., Rudicell, R.S., Georgiev, I.S., et al. (2015). Broadly Neutralizing Human Immunodeficiency Virus Type 1 Antibody Gene Transfer Protects Nonhuman Primates from Mucosal Simian-Human Immunodeficiency Virus Infection. Journal of Virology 89, 8334–8345. 10.1128/JVI.00908-15.

50. Welles, H.C., Jennewein, M.F., Mason, R.D., Narpala, S., Wang, L., Cheng, C., Zhang, Y., Todd, J.-P., Lifson, J.D., Balazs, A.B., et al. (2018). Vectored delivery of anti-SIV envelope targeting mAb via AAV8 protects rhesus macaques from repeated limiting dose intrarectal swarm SIVsmE660 challenge. PLoS Pathogens 14, e1007395. 10.1371/journal.ppat.1007395.

51. Casazza, J.P., Cale, E.M., Narpala, S., Yamshchikov, G.V., Coates, E.E., Hendel, C.S., Novik, L., Holman, L.A., Widge, A.T., Apte, P., et al. (2022). Safety and tolerability of AAV8 delivery of a broadly neutralizing antibody in adults living with HIV: a phase 1, dose-escalation trial. Nat Med. 10.1038/s41591-022-01762-x.

52. Fuchs, S.P., Mondragon, P.G., Zabizhin, R., Tomer, S., Wang, L., Cook, E., Dudley, D.M., Weisgrau, K.L., Furlott, J., Coonen, J., et al. (2025). Transient rapamycin treatment avoids unwanted host immune responses toward AAV-delivered anti-HIV antibodies. Nat Commun 16, 8906. 10.1038/s41467-025-63970-6.

53. Ardeshir, A., O’Hagan, D., Mehta, I., Shandilya, S., Hopkins, L.L.J., Adamson, L., Kuroda, M.J., Hahn, P.A., da Costa, L.A.B., Fuchs, S.P., et al. (2025). Determinants of successful AAV-vectored delivery of HIV-1 bNAbs in early life. Nature 645, 1020–1028. 10.1038/s41586-025-09330-2.

54. Thavarajah, J.J., Hønge, B.L., and Wejse, C.M. (2024). The Use of Broadly Neutralizing Antibodies (bNAbs) in HIV-1 Treatment and Prevention. Viruses 16, 911. 10.3390/v16060911.

55. Koyanagi, Y., Miles, S., Mitsuyasu, R.T., Merrill, J.E., Vinters, H.V., and Chen, I.S. (1987). Dual infection of the central nervous system by AIDS viruses with distinct cellular tropisms. Science 236, 819–822.

56. Ochsenbauer, C., Edmonds, T.G., Ding, H., Keele, B.F., Decker, J., Salazar, M.G., Salazar-Gonzalez, J.F., Shattock, R., Haynes, B.F., Shaw, G.M., et al. (2012). Generation of Transmitted/Founder HIV-1 Infectious Molecular Clones and Characterization of Their Replication Capacity in CD4 T Lymphocytes and Monocyte-Derived Macrophages. J Virol 86, 2715–2728. 10.1128/JVI.06157-11.

57. McCarthy, M., He, J., and Wood, C. (1998). HIV-1 Strain-Associated Variability in Infection of Primary Neuroglia. J Neurovirol 4, 80–89. 10.3109/13550289809113484.

58. Wright, S. (1932). The roles of mutation, inbreeding, crossbreeding, and selection in evolution. Proc. In Sixth Int. Cong. Genet 1, 356–366.

59. Melkus, M.W., Estes, J.D., Padgett-Thomas, A., Gatlin, J., Denton, P.W., Othieno, F.A., Wege, A.K., Haase, A.T., and Garcia, J.V. (2006). Humanized mice mount specific adaptive and innate immune responses to EBV and TSST-1. Nature Medicine 12, 1316–1322. 10.1038/nm1431.

60. Baenziger, S., Tussiwand, R., Schlaepfer, E., Mazzucchelli, L., Heikenwalder, M., Kurrer, M.O., Behnke, S., Frey, J., Oxenius, A., Joller, H., et al. (2006). Disseminated and sustained HIV infection in CD34+ cord blood cell-transplanted Rag2-/-gamma c-/- mice. Proceedings of the National Academy of Sciences of the United States of America 103, 15951–15956. 10.1073/pnas.0604493103.

61. Sun, Z., Denton, P.W., Estes, J.D., Othieno, F.A., Wei, B.L., Wege, A.K., Melkus, M.W., Padgett-Thomas, A., Zupancic, M., Haase, A.T., et al. (2007). Intrarectal transmission, systemic infection, and CD4+ T cell depletion in humanized mice infected with HIV-1. The Journal of experimental medicine 204, 705–714. 10.1084/jem.20062411.

62. Huang, J., Kang, B.H., Ishida, E., Zhou, T., Griesman, T., Sheng, Z., Wu, F., Doria-Rose, N.A., Zhang, B., McKee, K., et al. (2016). Identification of a CD4-Binding-Site Antibody to HIV that Evolved Near-Pan Neutralization Breadth. Immunity 45, 1108–1121. 10.1016/j.immuni.2016.10.027.

63. Sok, D., van Gils, M.J., Pauthner, M., Julien, J.-P., Saye-Francisco, K.L., Hsueh, J., Briney, B., Lee, J.H., Le, K.M., Lee, P.S., et al. (2014). Recombinant HIV envelope trimer selects for quaternary-dependent antibodies targeting the trimer apex. Proceedings of the National Academy of Sciences of the United States of America 111, 17624–17629. 10.1073/pnas.1415789111.

64. Webb, N.E., Montefiori, D.C., and Lee, B. (2015). Dose–response curve slope helps predict therapeutic potency and breadth of HIV broadly neutralizing antibodies. Nat Commun 6, 8443. 10.1038/ncomms9443.

65. Galvez, N.M.S., Sheehan, M.L., Lin, A.Z., Cao, Y., Lam, E.C., Jackson, A.M., and Balazs, A.B. (2024). QuickFit: A High-Throughput RT-qPCR-Based Assay to Quantify Viral Growth and Fitness In Vitro. Viruses 16, 1320. 10.3390/v16081320.

66. Wainburg, M.A. (2004). The impact of the M184V substitution on drug resistance and viral fitness. Expert Review of Anti-infective Therapy 2, 147–151. 10.1586/14787210.2.1.147.

67. Mendoza, P., Gruell, H., Nogueira, L., Pai, J.A., Butler, A.L., Millard, K., Lehmann, C., Suárez, I., Oliveira, T.Y., Lorenzi, J.C.C., et al. (2018). Combination therapy with anti-HIV-1 antibodies maintains viral suppression. Nature, 1–21. 10.1038/s41586-018-0531-2.

68. Julg, B., Stephenson, K.E., Wagh, K., Tan, S.C., Zash, R., Walsh, S., Ansel, J., Kanjilal, D., Nkolola, J., Walker-Sperling, V.E.K., et al. (2022). Safety and antiviral activity of triple combination broadly neutralizing monoclonal antibody therapy against HIV-1: a phase 1 clinical trial. Nat Med 28, 1288–1296. 10.1038/s41591-022-01815-1.

69. Pegu, A., Xu, L., DeMouth, M.E., Fabozzi, G., March, K., Almasri, C.G., Cully, M.D., Wang, K., Yang, E.S., Dias, J., et al. (2022). Potent anti-viral activity of a trispecific HIV neutralizing antibody in SHIV-infected monkeys. Cell Reports 38, 110199. 10.1016/j.celrep.2021.110199.

70. Lynch, R.M., Boritz, E., Coates, E.E., DeZure, A., Madden, P., Costner, P., Enama, M.E., Plummer, S., Holman, L., Hendel, C.S., et al. (2015). Virologic effects of broadly neutralizing antibody VRC01 administration during chronic HIV-1 infection. Science translational medicine 7, 319ra206–319ra206. 10.1126/scitranslmed.aad5752.

71. Schoofs, T., Klein, F., Braunschweig, M., Kreider, E.F., Feldmann, A., Nogueira, L., Oliveira, T., Lorenzi, J.C.C., Parrish, E.H., Learn, G.H., et al. (2016). HIV-1 therapy with monoclonal antibody 3BNC117 elicits host immune responses against HIV-1. Science. 10.1126/science.aaf0972.

72. Scheid, J.F., Horwitz, J.A., Bar-On, Y., Kreider, E.F., Lu, C.-L., Lorenzi, J.C.C., Feldmann, A., Braunschweig, M., Nogueira, L., Oliveira, T., et al. (2016). HIV-1 antibody 3BNC117 suppresses viral rebound in humans during treatment interruption. Nature. 10.1038/nature18929.

73. Otsuka, Y., Schmitt, K., Quinlan, B.D., Gardner, M.R., Alfant, B., Reich, A., Farzan, M., and Choe, H. (2018). Diverse pathways of escape from all well-characterized VRC01-class broadly neutralizing HIV-1 antibodies. PLoS Pathog 14, e1007238. 10.1371/journal.ppat.1007238.

74. Wise, M.C., Xu, Z., Tello-Ruiz, E., Beck, C., Trautz, A., Patel, A., Elliott, S.T.C., Chokkalingam, N., Kim, S., Kerkau, M.G., et al. (2020). In vivo delivery of synthetic DNA–encoded antibodies induces broad HIV-1–neutralizing activity. Journal of Clinical Investigation 130, 827–837. 10.1172/JCI132779.

75. Wu, R.L., Houser, K.V., Gaudinski, M.R., Widge, A.T., Awan, S.F., Carter, C.A., Holman, L.A., Saunders, J., Hendel, C.S., Eshun, A., et al. (2025). Safety and pharmacokinetics of N6LS, a broadly neutralising monoclonal antibody for HIV: a phase 1, open-label, dose-escalation study in healthy adults. The Lancet HIV. 10.1016/s2352-3018(25)00041-4.

76. Moore, P.L., Gray, E.S., Sheward, D., Madiga, M., Ranchobe, N., Lai, Z., Honnen, W.J., Nonyane, M., Tumba, N., Hermanus, T., et al. (2011). Potent and broad neutralization of HIV-1 subtype C by plasma antibodies targeting a quaternary epitope including residues in the V2 loop. J Virol 85, 3128–3141. 10.1128/JVI.02658-10.

77. Radford, C.E., and Bloom, J.D. (2025). Comprehensive maps of escape mutations from antibodies 10-1074 and 3BNC117 for Envs from two divergent HIV strains. bioRxiv, 2025.01.30.635715. 10.1101/2025.01.30.635715.

78. Neher, R.A., and Leitner, T. (2010). Recombination Rate and Selection Strength in HIV Intra-patient Evolution. PLoS Comput Biol 6, e1000660. 10.1371/journal.pcbi.1000660.

79. Li, Y., O’Dell, S., Walker, L.M., Wu, X., Guenaga, J., Feng, Y., Schmidt, S.D., McKee, K., Louder, M.K., Ledgerwood, J.E., et al. (2011). Mechanism of neutralization by the broadly neutralizing HIV-1 monoclonal antibody VRC01. J Virol 85, 8954–8967. 10.1128/JVI.00754-11.

80. Pegu, A., Borate, B., Huang, Y., Pauthner, M.G., Hessell, A.J., Julg, B., Doria-Rose, N.A., Schmidt, S.D., Carpp, L.N., Cully, M.D., et al. (2019). A Meta-analysis of Passive Immunization Studies Shows that Serum-Neutralizing Antibody Titer Associates with Protection against SHIV Challenge. Cell Host & Microbe 26, 336–346.e3. 10.1016/j.chom.2019.08.014.

81. Fellinger, C.H., Gardner, M.R., Weber, J.A., Alfant, B., Zhou, A.S., and Farzan, M. (2019). eCD4-Ig limits HIV-1 escape more effectively than CD4-Ig or a broadly neutralizing antibody. Journal of Virology. 10.1128/JVI.00443-19.

82. Bricault, C.A., Yusim, K., Seaman, M.S., Yoon, H., Theiler, J., Giorgi, E.E., Wagh, K., Theiler, M., Hraber, P., Macke, J.P., et al. (2019). HIV-1 Neutralizing Antibody Signatures and Application to Epitope-Targeted Vaccine Design. Cell Host & Microbe 25, 59–72.e8. 10.1016/j.chom.2018.12.001.

83. Asmal, M., Luedemann, C., Lavine, C.L., Mach, L.V., Balachandran, H., Brinkley, C., Denny, T.N., Lewis, M.G., Anderson, H., Pal, R., et al. (2015). Infection of monkeys by simian-human immunodeficiency viruses with transmitted/founder clade C HIV-1 envelopes. Virology 475, 37–45. 10.1016/j.virol.2014.10.032.

84. Zhang, C., Zaman, L.A., Poluektova, L.Y., Gorantla, S., Gendelman, H.E., and Dash, P.K. (2023). Humanized Mice for Studies of HIV-1 Persistence and Elimination. Pathogens 12, 879. 10.3390/pathogens12070879.

85. Wodarz, D., and Nowak, M.A. (1998). The effect of different immune responses on the evolution of virulent CXCR4–tropic HIV. Proc. R. Soc. Lond. B 265, 2149–2158. 10.1098/rspb.1998.0552.

86. Theys, K., Deforche, K., Beheydt, G., Moreau, Y., Van Laethem, K., Lemey, P., Camacho, R.J., Rhee, S.-Y., Shafer, R.W., Van Wijngaerden, E., et al. (2010). Estimating the individualized HIV-1 genetic barrier to resistance using a nelfinavir fitness landscape. BMC Bioinformatics 11, 409. 10.1186/1471-2105-11-409.

87. Ganusov, V.V., and De Boer, R.J. (2006). Estimating Costs and Benefits of CTL Escape Mutations in SIV/HIV Infection. PLoS Comput Biol 2, e24. 10.1371/journal.pcbi.0020024.

88. Read, E.L., Tovo-Dwyer, A.A., and Chakraborty, A.K. (2012). Stochastic effects are important in intrahost HIV evolution even when viral loads are high. Proc. Natl. Acad. Sci. U.S.A. 109, 19727–19732. 10.1073/pnas.1206940109.

89. Batorsky, R., Sergeev, R.A., and Rouzine, I.M. (2014). The Route of HIV Escape from Immune Response Targeting Multiple Sites Is Determined by the Cost-Benefit Tradeoff of Escape Mutations. PLoS Comput Biol 10, e1003878. 10.1371/journal.pcbi.1003878.

90. Gardner, E.M., Burman, W.J., Steiner, J.F., Anderson, P.L., and Bangsberg, D.R. (2009). Antiretroviral medication adherence and the development of class-specific antiretroviral resistance. AIDS 23, 1035–1046. 10.1097/QAD.0b013e32832ba8ec.

91. Armstrong, K.L., Lee, T.-H., and Essex, M. (2011). Replicative Fitness Costs of Nonnucleoside Reverse Transcriptase Inhibitor Drug Resistance Mutations on HIV Subtype C. Antimicrob Agents Chemother 55, 2146–2153. 10.1128/AAC.01505-10.

92. Hill, A.L., Rosenbloom, D.I.S., and Nowak, M.A. (2012). Evolutionary dynamics of HIV at multiple spatial and temporal scales. J Mol Med 90, 543–561. 10.1007/s00109-012-0892-1.

93. Zanini, F., Puller, V., Brodin, J., Albert, J., and Neher, R.A. (2017). In vivo mutation rates and the landscape of fitness costs of HIV-1. Virus Evolution 3, vex003. 10.1093/ve/vex003.

94. Meijers, M., Vanshylla, K., Gruell, H., Klein, F., and Lässig, M. (2021). Predicting in vivo escape dynamics of HIV-1 from a broadly neutralizing antibody. Proc. Natl. Acad. Sci. U.S.A. 118, e2104651118. 10.1073/pnas.2104651118.

95. Cong, M., Heneine, W., and García-Lerma, J.G. (2007). The Fitness Cost of Mutations Associated with Human Immunodeficiency Virus Type 1 Drug Resistance Is Modulated by Mutational Interactions. J Virol 81, 3037–3041. 10.1128/jvi.02712-06.

96. Gonelli, C.A., King, H.A.D., Ko, S., Fennessey, C.M., Iwamoto, N., Mason, R.D., Heimann, A., Flebbe, D.R., Todd, J.-P., Foulds, K.E., et al. (2025). Antibody prophylaxis may mask subclinical SIV infections in macaques. Nature 639, 205–213. 10.1038/s41586-024-08500-y.

97. Caskey, M. (2020). Broadly neutralizing antibodies for the treatment and prevention of HIV infection. Curr Opin HIV AIDS 15, 49–55. 10.1097/COH.0000000000000600.

98. Han, C., Johnson, J., Dong, R., Kandula, R., Kort, A., Wong, M., Yang, T., Breheny, P.J., Brown, G.D., and Haim, H. (2020). Key Positions of HIV-1 Env and Signatures of Vaccine Efficacy Show Gradual Reduction of Population Founder Effects at the Clade and Regional Levels. mBio 11, e00126–20. 10.1128/mBio.00126-20.

99. Haddox, H.K., Dingens, A.S., Hilton, S.K., Overbaugh, J., and Bloom, J.D. (2018). Mapping mutational effects along the evolutionary landscape of HIV envelope. eLife 7. 10.7554/eLife.34420.

100. DeLeon, O., Hodis, H., O’Malley, Y., Johnson, J., Salimi, H., Zhai, Y., Winter, E., Remec, C., Eichelberger, N., Van Cleave, B., et al. (2017). Accurate predictions of population-level changes in sequence and structural properties of HIV-1 Env using a volatility-controlled diffusion model. PLoS Biol 15, e2001549. 10.1371/journal.pbio.2001549.

101. Karpel, M.E., Boutwell, C.L., and Allen, T.M. (2015). BLT humanized mice as a small animal model of HIV infection. Curr Opin Virol 13, 75–80. 10.1016/j.coviro.2015.05.002.

102. Seung, E., and Tager, A.M. (2013). Humoral immunity in humanized mice: a work in progress. J Infect Dis 208 Suppl 2, S155–159. 10.1093/infdis/jit448.

103. McCoy, L.E., van Gils, M.J., Ozorowski, G., Messmer, T., Briney, B., Voss, J.E., Kulp, D.W., Macauley, M.S., Sok, D., Pauthner, M., et al. (2016). Holes in the Glycan Shield of the Native HIV Envelope Are a Target of Trimer-Elicited Neutralizing Antibodies. Cell Reports 16, 2327–2338. 10.1016/j.celrep.2016.07.074.

104. Freund, N.T., Wang, H., Scharf, L., Nogueira, L., Horwitz, J.A., Bar-On, Y., Golijanin, J., Sievers, S.A., Sok, D., Cai, H., et al. (2017). Coexistence of potent HIV-1 broadly neutralizing antibodies and antibody-sensitive viruses in a viremic controller. Science translational medicine 9, eaal2144. 10.1126/scitranslmed.aal2144.

105. Ramakrishnan, M.A. (2016). Determination of 50% endpoint titer using a simple formula. World journal of virology 5, 85–86. 10.5501/wjv.v5.i2.85.

106. Chen, S., Zhou, Y., Chen, Y., and Gu, J. (2018). fastp: an ultra-fast all-in-one FASTQ preprocessor. Bioinformatics (Oxford, England) 34, i884–i890. 10.1093/bioinformatics/bty560.

107. Langmead, B., and Salzberg, S.L. (2012). Fast gapped-read alignment with Bowtie 2. Nature Methods 9, 357–359. 10.1038/nmeth.1923.

108. Li, H., Handsaker, B., Wysoker, A., Fennell, T., Ruan, J., Homer, N., Marth, G., Abecasis, G., Durbin, R., and 1000 Genome Project Data Processing Subgroup (2009). The Sequence Alignment/Map format and SAMtools. Bioinformatics (Oxford, England) 25, 2078–2079. 10.1093/bioinformatics/btp352.

109. Korber, B.T., Foley, B.T., Kuiken, C.L., Pillai, S.K., and Sodroski, J.G. (2014). Numbering Positions in HIV Relative to HXB2CG. LANL. https://www.hiv.lanl.gov/content/sequence/HIV/REVIEWS/HXB2.html.

110. Köster, J., and Rahmann, S. (2012). Snakemake–a scalable bioinformatics workflow engine. Bioinformatics (Oxford, England) 28, 2520–2522. 10.1093/bioinformatics/bts480.

111. Wilm, A., Aw, P.P.K., Bertrand, D., Yeo, G.H.T., Ong, S.H., Wong, C.H., Khor, C.C., Petric, R., Hibberd, M.L., and Nagarajan, N. (2012). LoFreq: a sequence-quality aware, ultra-sensitive variant caller for uncovering cell-population heterogeneity from high-throughput sequencing datasets. Nucleic Acids Research 40, 11189–11201. 10.1093/nar/gks918.

112. Piantadosi, A., Freije, C.A., Gosmann, C., Ye, S., Park, D., Schaffner, S.F., Tully, D.C., Allen, T.M., Dong, K.L., Sabeti, P.C., et al. (2019). Metagenomic Sequencing of HIV-1 in the Blood and Female Genital Tract Reveals Little Quasispecies Diversity during Acute Infection. Journal of Virology 93. 10.1128/JVI.00804-18.

113. McCrone, J.T., and Lauring, A.S. (2016). Measurements of Intrahost Viral Diversity Are Extremely Sensitive to Systematic Errors in Variant Calling. Journal of Virology 90, 6884–6895. 10.1128/JVI.00667-16.

114. Siebring-van Olst, E., Vermeulen, C., de Menezes, R.X., Howell, M., Smit, E.F., and van Beusechem, V.W. (2013). Affordable Luciferase Reporter Assay for Cell-Based High-Throughput Screening. J Biomol Screen 18, 453–461. 10.1177/1087057112465184.

115. Borchers, H.W. (2023). pracma: Practical Numerical Math Functions. Version R package version 2.4.4.

116. R Core Team (2024). R: A Language and Environment for Statistical Computing. (R Foundation for Statistical Computing).

117. Reagan-Shaw, S., Nihal, M., and Ahmad, N. (2008). Dose translation from animal to human studies revisited. The FASEB journal1: official publication of the Federation of American Societies for Experimental Biology 22, 659–661. 10.1096/fj.07-9574LSF.

