## Supplemental Figures for "HIV broadly neutralizing antibody escape dynamics drive the outcome of AAV vectored immunotherapy in humanized mice"

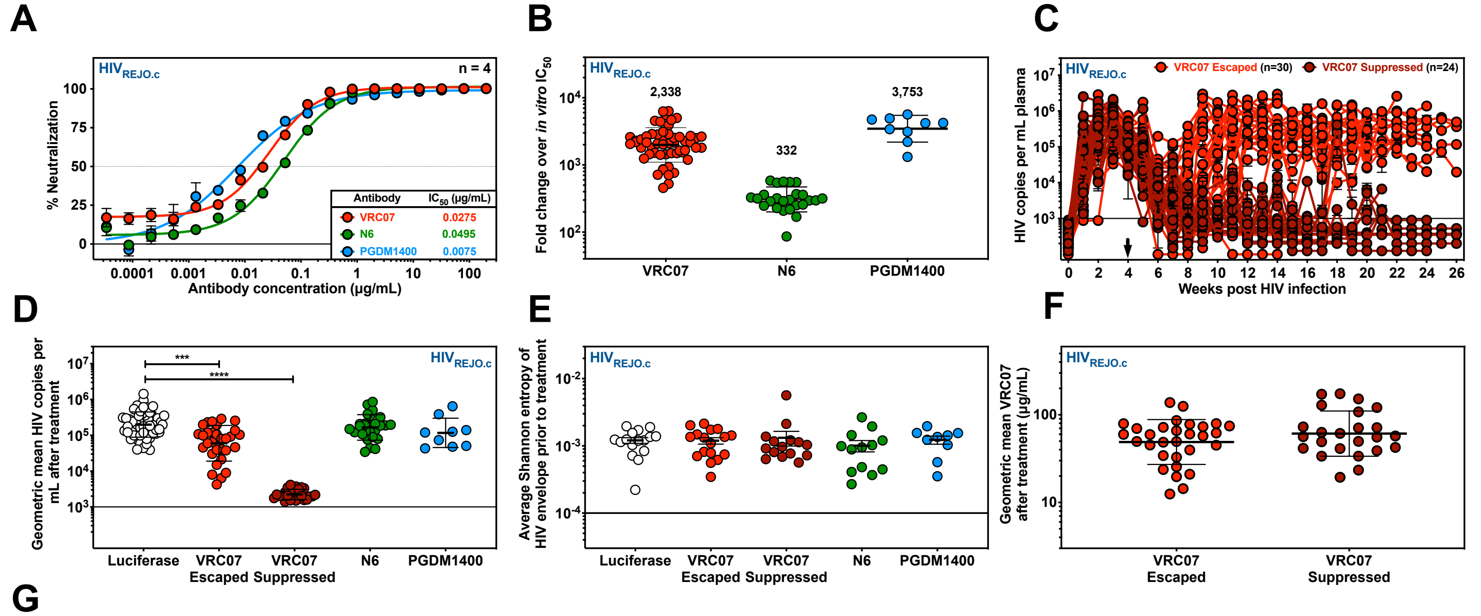

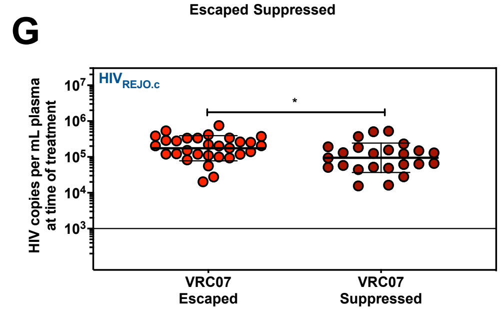


**Figure S1. AAV-VRC07 can suppress $HIV_{REJO.c}$ infection in BLT humanized mice (Related to Figure 1).**

**(A)** *In vitro* neutralization of $HIV_{REJO.c}$ by VRC07, N6, or PGDM1400 bNAbs. Data are plotted as mean ± S.E.M. Each datapoint was evaluated in quadruplicate.

**(B)** The fold change above *in vitro* $IC_{50}$ was calculated as the geometric mean of bNAb expression after AAV administration for each mouse, divided by the $IC_{50}$ value determined *in vitro* for the corresponding bNAb against $HIV_{REJO.c}$. Numbers above each group represent the average fold change for each bNAb-$IC_{50}$ pair. Error bars indicate geometric SD.

**(C)** HIV viral load in plasma of $HIV_{REJO.c}$-infected mice treated with AAV-VRC07, as seen in **Figure 1 (D)**. Each colored line depicts an individual mouse. Black arrow denotes vector administration. The qPCR lower limit of detection was 1 genome copy per µL of plasma, and 5µL were used in the reaction (solid line). Data are presented as mean ± S.E.M. Individual AAV-VRC07-treated mice that escaped or were suppressed are denoted by light or dark red colors, respectively.

**(D)** Geometric mean viral load of mice in **C** following vector administration over the period of observation. (***:p<0.005; ****:p<0.001; One-way ANOVA with Dunnett's *post hoc* test to correct for multiple comparisons). Error bars indicate geometric SD.

**(E)** HIV viral diversity prior to vector administration as quantified by average Shannon entropy of envelope-amplicons measured by Illumina Deep Sequencing (p=0.8755, One-way ANOVA with Tukey's *post hoc* test). Data are presented as mean ± S.E.M.

**(F)** Geometric mean of VRC07 antibody concentration in plasma following AAV-VRC07 vector administration in mice that were escaped or suppressed (p=0.1305, unpaired two-tailed Student's t test). Error bars indicate geometric SD.

**(G)** HIV viral load in plasma of $HIV_{REJO.c}$-infected mice at the time of vector administration, grouped by eventual suppression or escape. Statistical differences were assessed by an unpaired two-tailed Student's t-test (*: p<0.05). Data are geometric mean ± geometric SD.

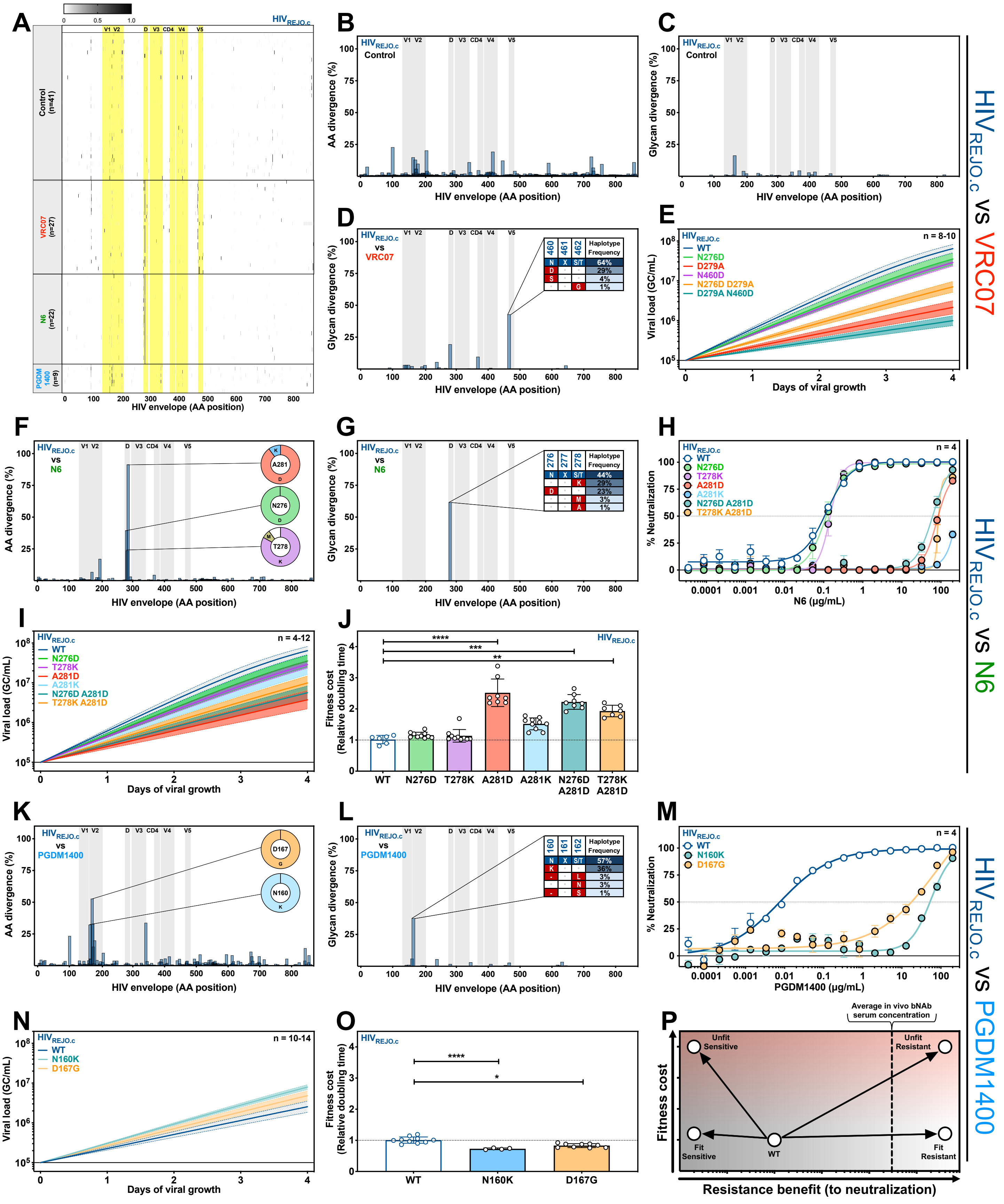


**Figure S2. Detailed escape data for $HIV_{REJO.c}$-infected mice treated with VRC07, N6, and PGDM1400 (Related to Figure 2).**

**Figure S2. Detailed escape data for $HIV_{REJO.c}$-infected mice treated with VRC07, N6, and PGDM1400 (Related to Figure 2).**

**(A)** Haplotypes of individual mice infected with $HIV_{REJO.c}$ treated with vectored VRC07, N6, or PGDM1400 as determined by Illumina Deep Sequencing of the viral envelope isolated from plasma at the final experimental timepoint. Each row represents a single mouse, with the color intensity of each cell denoting mutational frequency at each site. The X-axis represents the HIV envelope amino acid position relative to $HIV_{HXB2}$ numbering. Yellow shading denotes the position of individual loops within the HIV envelope (as indicated at the top).

**(B)** Amino acid divergence from the envelope gene of the $HIV_{REJO.c}$ parental strain across all control-treated mice. Sequences were determined by Illumina Deep Sequencing of the viral envelope isolated from plasma at the final experimental time point. The X-axis represents the envelope protein amino acid position relative to $HIV_{HXB2}$ numbering. The Y-axis represents the percentage of average amino acid divergence from the parental strain. Shaded areas denote the position of individual loops within the HIV envelope (as indicated at the top).

**(C-D)** Glycosylation site divergence from the reference strain for $HIV_{REJO.c}$ across all control animals **(C)** and all VRC07-treated mice **(D)**, as determined by Illumina Deep Sequencing of the viral envelope isolated from plasma at the final experimental timepoint. The X-axis represents the envelope protein amino acid position of each potential N-linked glycosylation site relative to $HIV_{HXB2}$ numbering. The Y-axis represents the average divergence from the parental strain envelope glycosylation sites. **(D)** is corrected for the glycan divergence seen in control mice. Potential N-linked glycosylation (PNGs) sites in the sequences were defined as Asn-X-Thr/Ser (NXT/S, with X being any amino acid but Pro), and the reference sequences with their respective haplotype frequencies are shown at the top of each plot legend. Shaded areas denote the position of individual loops within the HIV envelope (as indicated at the top).

**(E)** Viral growth curves of $HIV_{REJO.c}$ mutants identified as potential VRC07 escapes in activated $CD4^+$ T cells generated using growth rates derived from the *QuickFit* assay. Data are plotted as mean ± 95% C.I.

**(F-G)** Amino acid divergence **(F)** and Glycosylation site divergence **(G)** from the envelope gene of the $HIV_{REJO.c}$ parental strain across all N6-treated mice. Pie charts represent the most common amino acid mutations for sites with the highest divergence, and tables show the reference glycan-related sequences with their respective haplotype frequencies.

**(H)** *In vitro* neutralization assay of $HIV_{REJO.c}$ mutants identified as potential N6 escapes against N6. Data are plotted as mean ± S.E.M. Each datapoint was evaluated in quadruplicate.

**(I)** Viral growth curves of $HIV_{REJO.c}$ mutants identified as potential N6 escapes in activated $CD4^+$ T cells generated using growth rates derived from the *QuickFit* assay. Data are plotted as mean ± 95% C.I.

**(J)** Relative viral growth of $HIV_{REJO.c}$ mutants identified as potential N6 escapes. Growth rates were determined by the *QuickFit* assay and normalized to the parental strain. Data are plotted as mean ± S.E.M., and statistical differences were assessed by a Kruskal-Wallis non-parametric ANOVA with Dunn's *post hoc* test to correct for multiple comparisons (**: $p<0.01$; ***: $p<0.001$; ****: $p<0.0001$).

**(K-L)** Amino acid divergence **(K)** and Glycosylation site divergence **(L)** from the envelope gene of the $HIV_{REJO.c}$ parental strain across all PGDM1400-treated mice.

**(M)** *In vitro* neutralization assay of $HIV_{REJO.c}$ mutants identified as potential PGDM1400 escapes against PGDM1400. Data are plotted as mean ± S.E.M. Each datapoint was evaluated in quadruplicate.

**(N)** Viral growth curves of $HIV_{REJO.c}$ mutants identified as potential PGDM1400 escapes using the *QuickFit* assay. Data are plotted as mean ± 95% C.I.

**(O)** Relative viral growth of $HIV_{REJO.c}$ mutants identified as potential PGDM1400 escapes using the *QuickFit* assay. Data are plotted as mean ± S.E.M., and statistical differences were assessed by a Kruskal-Wallis non-parametric ANOVA with Dunn's *post hoc* test to correct for multiple comparisons (*: $p<0.05$; ****: $p<0.0001$).

**(P)** Theoretical escapability maps demonstrating the potential fitness cost (Y-axis) and resistance benefits (X-axis) for WT HIV and hypothetical mutants that could arise during antibody selective pressure. The dashed vertical line denotes the hypothetical geometric mean serum concentration of the evaluated antibody. Arrows represent potential paths taken during selection.

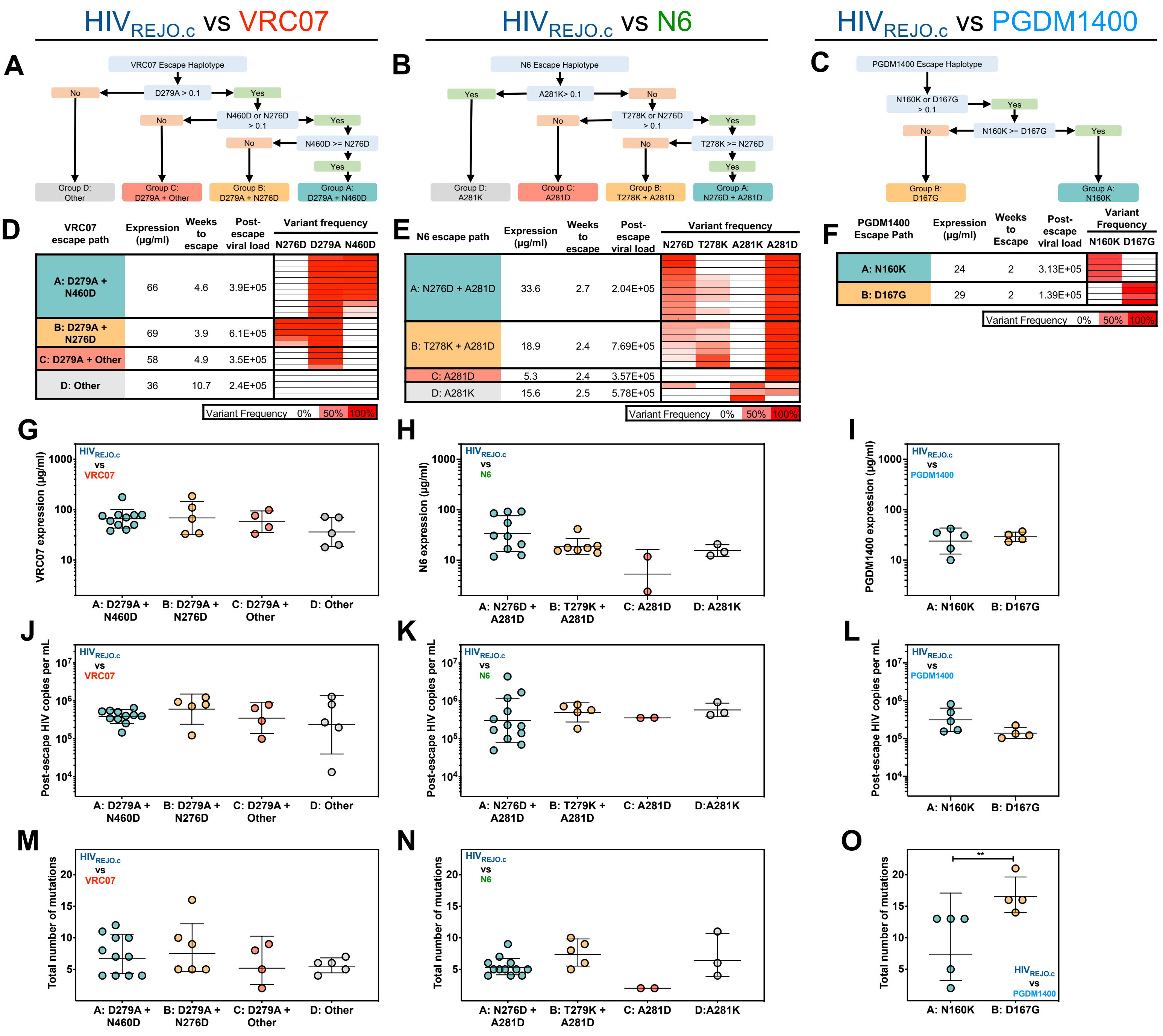


**Figure S3. Escape path analysis for HIV$_{REJO.c}$-infected mice treated with various bNAbs (Related to Figure 2).**
**(A-C)** Escape path haplotype categorization algorithms for HIV$_{REJO.c}$ escapes from VRC07 **(A)**, N6 **(B)**, and PGDM1400 **(C)**. Sample haplotypes were generated by evaluating the frequency of variants greater than 10%.
**(D-F)** Escape path categories and metadata for each HIV$_{REJO.c}$ escape sample from VRC07 **(D)**, N6 **(E)**, and PGDM1400 **(F)**. The escape paths were assigned using the antibody-specific algorithms described in **(A-C).** The geometric mean of antibody expression, weeks to viral escape, and post-escape viral load are grouped by escape path, and within each group, the escape variant frequency is reported for each mouse (row).
**(G-I)** Serum bNAb expression of VRC07 **(G)**, N6 **(H)**, and PGDM1400 **(I)** grouped by escape path. Data are geometric mean ± geometric SD, and each point represents one mouse. To compare the levels of bNAb expression across various escape paths, a Kruskal-Wallis non-parametric ANOVA test with Dunn's *post hoc* test to correct for multiple comparisons was performed for **(G)** and **(H);** while a Mann-Whitney test was performed for **(I).** No statistical differences were found ($p>0.05$).
**(J-L)** Post-escape viral load for VRC07 **(J)**, N6 **(K)**, and PGDM1400 **(L)** grouped by escape path. The post-escape viral load was determined by calculating the geometric mean of the viral load from 2 weeks post-escape until either the mouse died, or the experiment ended. Data are geometric mean ± geometric SD, and each point represents one individual mouse sample. To compare the post-escape viral load levels across various escape paths, a Kruskal-Wallis non-parametric ANOVA test with Dunn's *post hoc* test to correct for multiple comparisons was performed for **(J)** and **(K);** while a Mann-Whitney test was performed for **(L).** No statistical differences were found for any of the comparisons performed ($p>0.05$).
**(M-O)** The total number of mutations post-escape for VRC07 **(M)**, N6 **(N)**, and PGDM1400 **(O)** grouped by escape path. This accounts for the total number of variants with a frequency of at least 10%, including the variants used to categorize the haplotypes. To compare the various escape paths, for **(M)** and **(N)**, a Kruskal-Wallis non-parametric ANOVA test with Dunn's *post hoc* test to correct for multiple comparisons was performed. For **(O),** a Mann-Whitney test was performed. Only comparisons with a significant p-value are indicated on the plots (**: $p<0.01$).

A B C

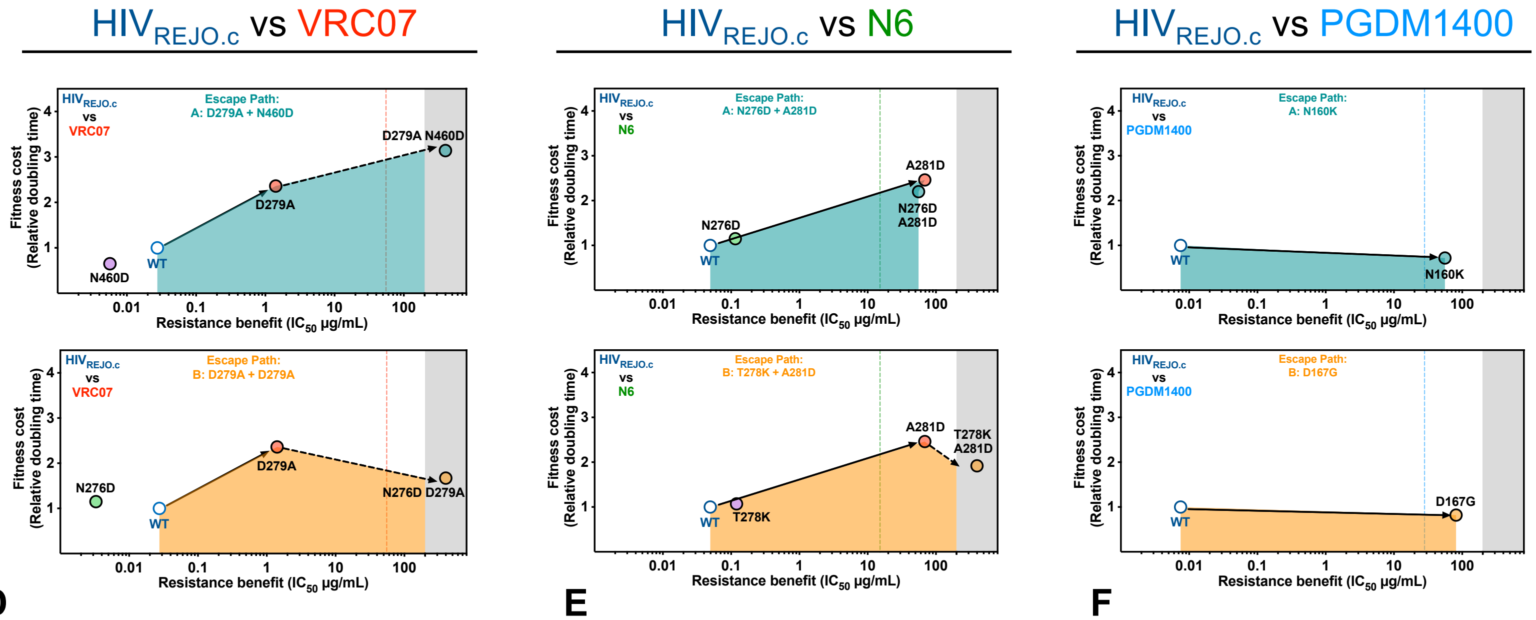


D

| Antibody | Fraction of Total Escapes | Escape Path | Raw AUC | Relative Fraction | Scaled AUC | Escape Barrier Score |
|---|---|---|---|---|---|---|
| VRC07 | 65% | A | 458 | 65% | 355 | 497 |
| | | B | 402 | 35% | 142 | |
| N6 | 77% | A | 88 | 59% | 52 | 219 |
| | | B | 406 | 41% | 167 | |
| PGDM1400 | 100% | A | 48 | 56% | 27 | 59 |
| | | B | 74 | 44% | 32 | |

E

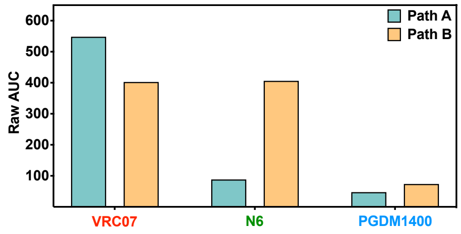


F

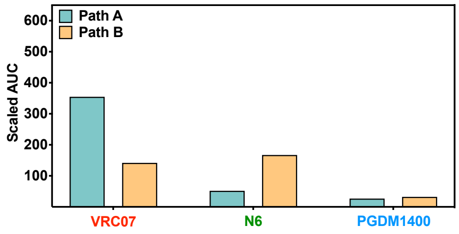


**Figure S4. Escape Barrier score calculation for the primary $HIV_{REJO.c}$ escape paths from various bNAbs (Related to Figure 2).**

**(A-C)** Visualized Escape Barrier score calculation for $HIV_{REJO.c}$ escape paths A (**upper**) and B (**lower**) against VRC07 **(A)**, N6 **(B)**, and PGDM1400 **(C)**. For each escape path, the path-specific barrier to escape is calculated as the area under the curve (AUC) across the accumulated resistance benefit from the WT $HIV_{REJO.c}$ position to the haplotype position.

**(D)** Escape Barrier score calculations for $HIV_{REJO.c}$ escape from VRC07, N6, and PGDM1400. For each antibody, the fraction of all escapes that were categorized as paths A or B were calculated. To compute the Escape Barrier score for each antibody, the AUC was calculated for each path and was then scaled by the relative path frequency, resulting in a scaled AUC. The scaled AUC values are then summed to create the Escape Barrier scores.

**(E)** Raw AUC Escape Barrier scores for paths A and B for $HIV_{REJO.c}$ escapes from VRC07, N6, and PGDM1400.

**(F)** Scaled AUC Escape Barrier scores for paths A and B for $HIV_{REJO.c}$ escapes from VRC07, N6, and PGDM1400.

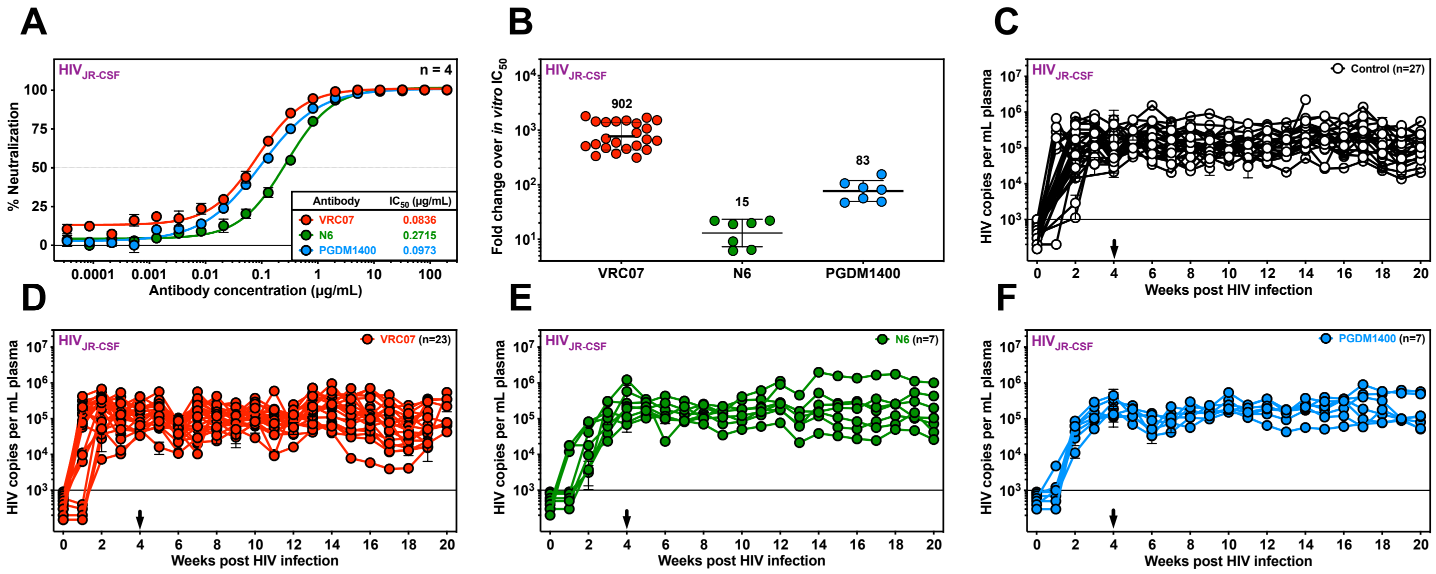


**Figure S5. Vectored delivery of VRC07, N6, or PGDM1400 bNAbs fails to suppress $HIV_{JR\text{-}CSF}$ in BLT humanized mice (Related to Figure 3).**

**(A)** *In vitro* neutralization of $HIV_{JR\text{-}CSF}$ by VRC07, PGDM1400, or N6. Data are plotted as mean ± S.E.M. Each datapoint was evaluated in quadruplicate.

**(B)** The fold change above *in vitro* $IC_{50}$ was calculated as the geometric mean of bNAb expression after AAV administration for each mouse, over the $IC_{50}$ *in vitro* value determined for the corresponding bNAb against $HIV_{JR\text{-}CSF}$. Numbers above each group represent the average fold change for each bNAb-$IC_{50}$ pair. Error bars indicate geometric SD.

**(C-F)** HIV viral load in plasma of $HIV_{JR\text{-}CSF}$-infected BLT mice following administration of AAV-Luciferase or AAV-2A10 controls **(C)**, AAV-VRC07 **(D)**, AAV-N6 **(E)**, or AAV-PGDM1400 **(F)**. Black arrows denote vector administration. Each colored line depicts an individual mouse tracked over time. Data are presented as mean ± S.E.M.

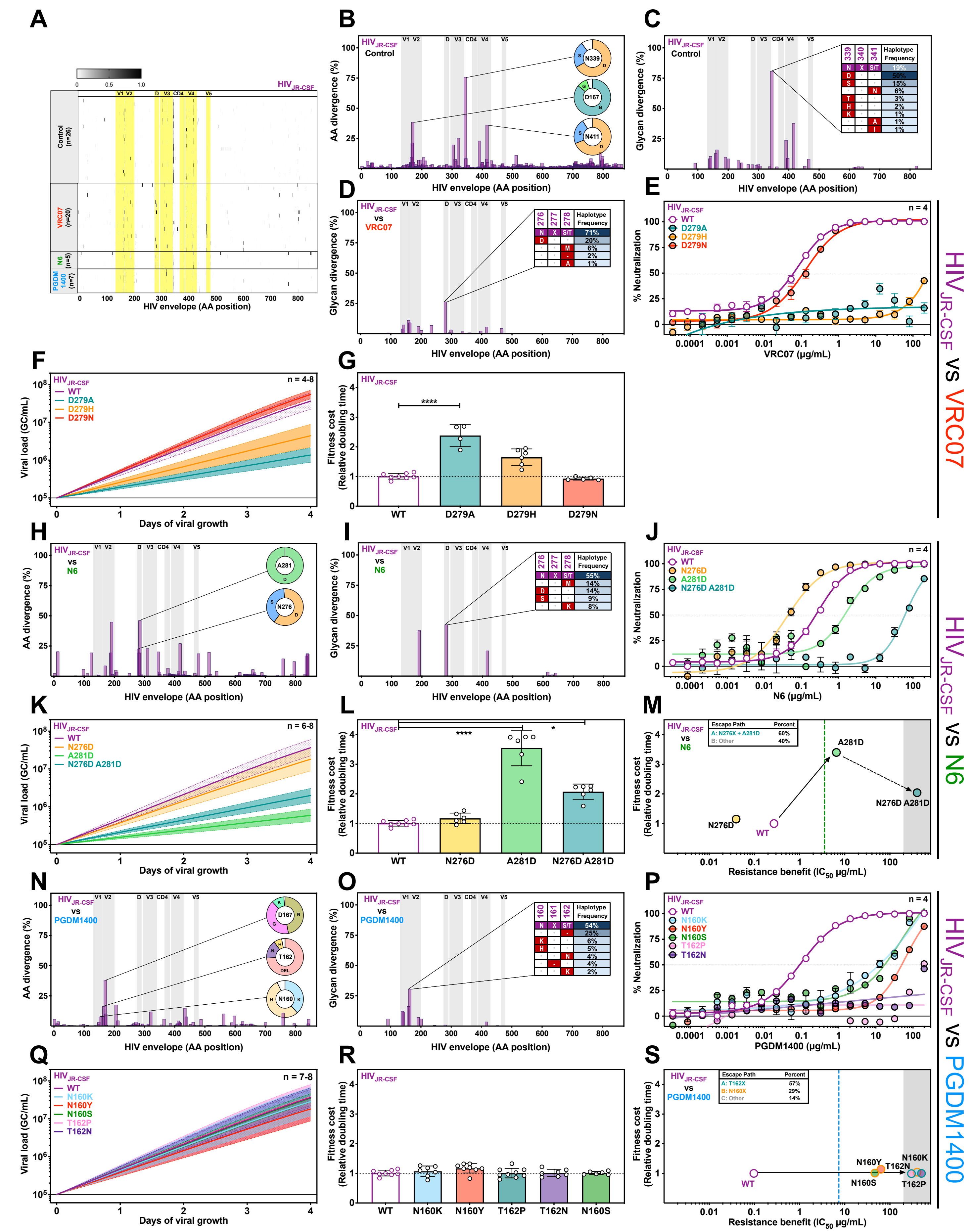


**Figure S6. Detailed escape data for HIV$_{JR\text{-}CSF}$-infected mice treated with VRC07, N6, and PGDM1400 (Related to Figure 3).**

**Figure S6. Detailed escape data for $HIV_{JR\text{-}CSF}$-infected mice treated with VRC07, N6, and PGDM1400 (Related to Figure 3).**

**(A)** Haplotypes of individual mice infected with $HIV_{JR\text{-}CSF}$ treated with vectored VRC07, N6, or PGDM1400.

**(B)** Amino acid divergence from the envelope gene of the $HIV_{JR\text{-}CSF}$ parental strain across all control-treated mice.

**(C-D)** Glycosylation site divergence from the reference strain for $HIV_{JR\text{-}CSF}$ across all control animals **(C)** and all VRC07-treated **(D)** mice.

**(E)** *In vitro* neutralization assay of $HIV_{JR\text{-}CSF}$ mutants identified as potential VRC07 escapes against VRC07. Data are plotted as mean ± S.E.M. Each datapoint was evaluated in quadruplicate.

**(F)** Viral growth curves of $HIV_{JR\text{-}CSF}$ mutants identified as potential VRC07 escapes using the *QuickFit* assay. Data are plotted as mean ± 95% C.I.

**(G)** Relative viral growth of $HIV_{JR\text{-}CSF}$ mutants identified as potential VRC07 escapes using the *QuickFit* assay and normalized to the parental strain. Data are plotted as mean ± S.E.M. and statistical differences were assessed by a Kruskal-Wallis non-parametric ANOVA with Dunn's *post hoc* test to correct for multiple comparisons (****: $p<0.0001$).

**(H-I)** Amino acid divergence **(H)** and Glycosylation site divergence **(I)** from the envelope gene of the $HIV_{JR\text{-}CSF}$ parental strain across all N6-treated mice.

**(J)** *In vitro* neutralization assay of $HIV_{JR\text{-}CSF}$ mutants identified as potential N6 escapes against N6. Data are plotted as mean ± S.E.M. Each datapoint was evaluated in quadruplicate.

**(K)** Viral growth curves of $HIV_{JR\text{-}CSF}$ mutants identified as potential N6 escapes using growth rates derived from the *QuickFit* assay. Data are plotted as mean ± 95% C.I.

**(L)** Relative viral growth of $HIV_{JR\text{-}CSF}$ mutants identified as potential N6 escapes using the *QuickFit* assay and normalized to the parental strain. Data are plotted as mean ± S.E.M. and statistical differences were assessed by a Kruskal-Wallis non-parametric ANOVA with Dunn's *post hoc* test to correct for multiple comparisons (*: $p<0.05$: ****: $p<0.0001$).

**(M)** Escapability map denoting fitness cost and resistance benefits for each $HIV_{JR\text{-}CSF}$ mutant observed during escape from N6.

**(N-O)** Amino acid divergence **(N)** and Glycosylation site divergence **(O)** from the envelope gene of the $HIV_{JR\text{-}CSF}$ parental strain across all PGDM1400-treated mice.

**(P)** *In vitro* neutralization assay of $HIV_{JR\text{-}CSF}$ mutants identified as potential PGDM1400 escapes against PGDM1400. Data are plotted as mean ± S.E.M. Each datapoint was evaluated in quadruplicate.

**(Q)** Viral growth curves of $HIV_{JR\text{-}CSF}$ mutants identified as potential PGDM1400 escapes using the *QuickFit* assay. Data are plotted as mean ± 95% C.I.

**(R)** Relative viral growth of $HIV_{JR\text{-}CSF}$ mutants identified as potential PGDM1400 escapes using the *QuickFit* assay and normalized to the parental strain. Data are plotted as mean ± S.E.M., and statistical differences were assessed by a Kruskal-Wallis non-parametric ANOVA with Dunn's *post hoc* test to correct for multiple comparisons.

**(S)** Escapability map denoting fitness cost and resistance benefits for each $HIV_{JR\text{-}CSF}$ mutant observed during escape from PGDM1400.

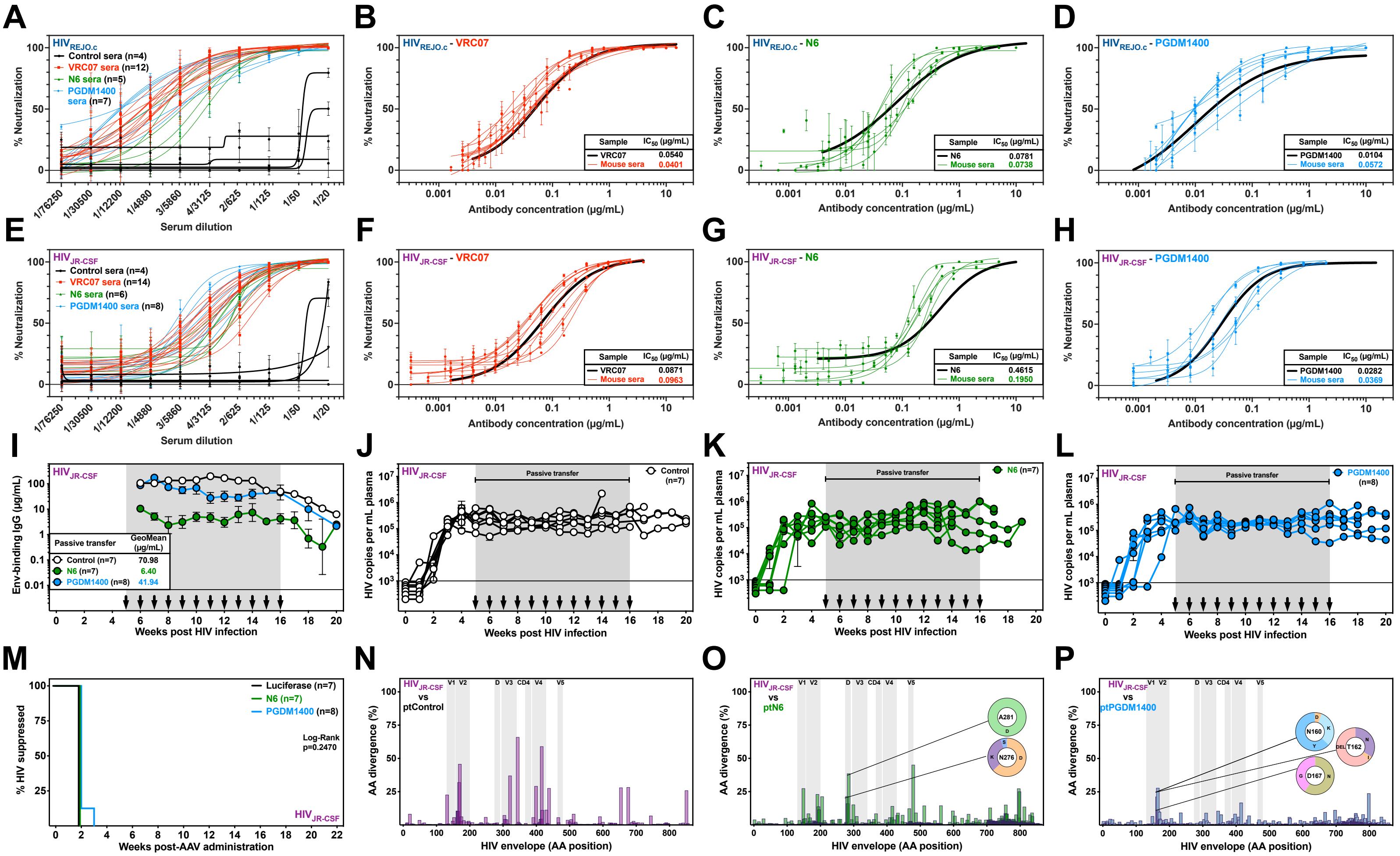


**Figure S7. Vectored antibodies retain neutralizing activity *in vivo* and passively transferred bNAbs fail to suppress HIV$_{JR\text{-}CSF}$ in the BLT model due to escape.**

**(A-H)** Sera from the terminal timepoint in mice from Figures 1 and 3 (colored lines) were tested for neutralization activity against HIV$_{REJO.c}$ (A-D) and HIV$_{JR\text{-}CSF}$ (E-H). Dilutions of sera retained potent neutralizing activity as compared to control animals (A, E) and this activity was equivalent to that seen for purified proteins (bold black lines) (B-D, F-H).

**(I)** ELISA-based quantitation of gp120-binding antibodies in the serum of HIV$_{JR\text{-}CSF}$-infected humanized mice following passive transfer of control antibody, N6 and PGDM1400. Black arrows denote weekly antibody administration. Data are plotted as geometric mean ± geometric SD.

**(J-L)** HIV viral load in plasma of HIV$_{JR\text{-}CSF}$-infected mice during passive transfer of control **(J)**, N6 **(K)**, or PGDM1400 **(L)**. Black arrows denote weekly protein administration. Each colored line depicts an individual mouse tracked over time. The sensitivity of qPCR was 1 genome copy per μL of plasma, and 5 μL were used in the reaction, resulting in a 1000 copy per mL limit of detection (solid line). Data are presented as mean ± S.E.M.

**(M)** Kaplan-Meier plot of viral suppression in BLT humanized mice infected with HIV$_{JR\text{-}CSF}$ given the indicated antibody. The total model was not significant (p=0.247) as assessed using a Log-rank (Mantel-Cox) test. The percentage of HIV suppressed was defined as the fraction of mice that were not escaped as described in the methods.

**(N-P)** Amino acid divergence from the envelope gene of the HIV$_{JR\text{-}CSF}$ parental strain across all mice passively transferred with control **(N)**, N6 **(O)**, or PGDM1400 **(P)** antibodies. Sequences were determined by Illumina Deep Sequencing of the viral envelope isolated from plasma at the final experimental time point. The X-axis represents the envelope protein amino acid position relative to HIV$_{HXB2}$ numbering. The Y-axis represents the percentage of average amino acid divergence from the parental strain. Shaded areas denote the position of individual loops within the HIV envelope (as indicated at the top).

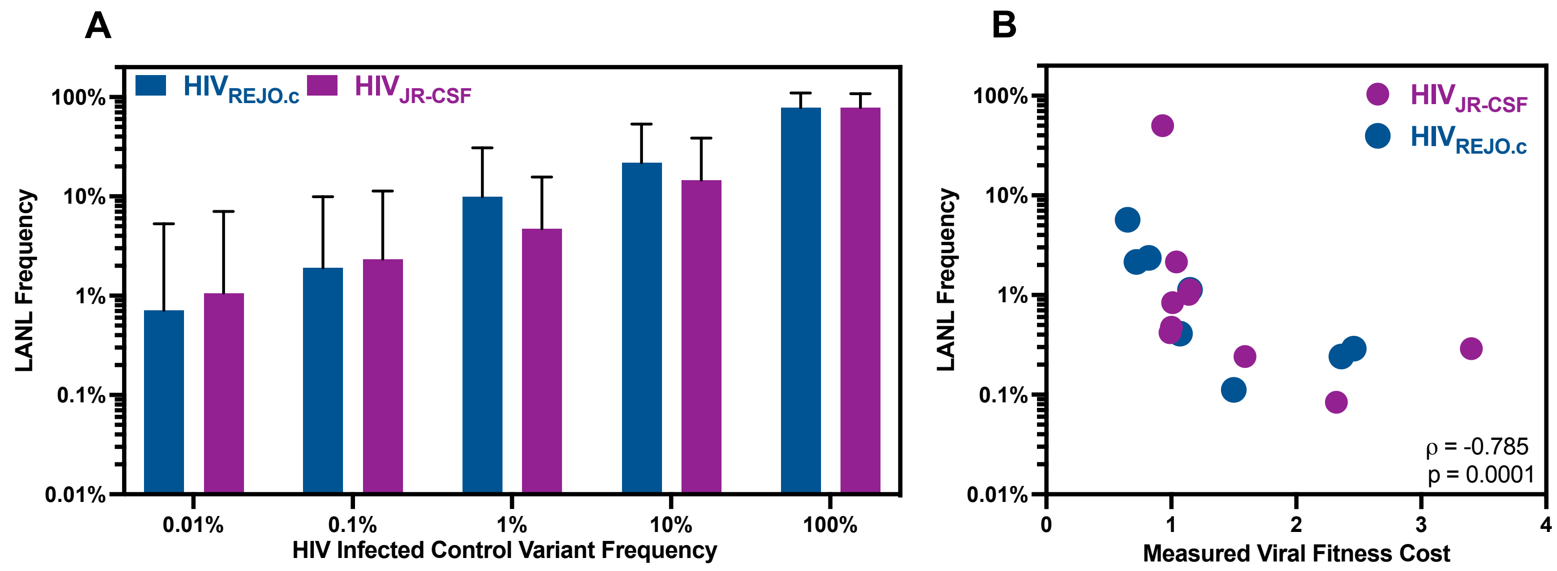


**Figure S8. Comparative analysis of amino acid frequencies in control mice and LANL database (Related to Figures 1 and 3).**

**(A)** Frequency of amino acid changes in HIV infected control mice compared to the same amino acid changes in the LANL database. Briefly, the frequency of each amino acid observed in $HIV_{REJO.c}$- or $HIV_{JR\text{-}CSF}$-infected mice with no antibody treatment (as seen in Figures S2B and S6B, respectively), were binned into five groups based on their frequency. Then the frequency of the same amino acid seen in LANL subtype B sequences were averaged for each of the five groups. Infrequent amino acids in the infected mice were also infrequent in the LANL database.

**(B)** Measured fitness cost of primary escape mutations profiled in this study compared to their frequency in the LANL database. Briefly, the fitness cost of primary single escape mutations (Table S1) as measured using the *QuickFit* assay were compared to frequency in LANL subtype B sequences (Table S6). Spearman correlation analysis demonstrated a strong and significant negative correlation between the measured viral fitness cost and the frequency in the LANL database.

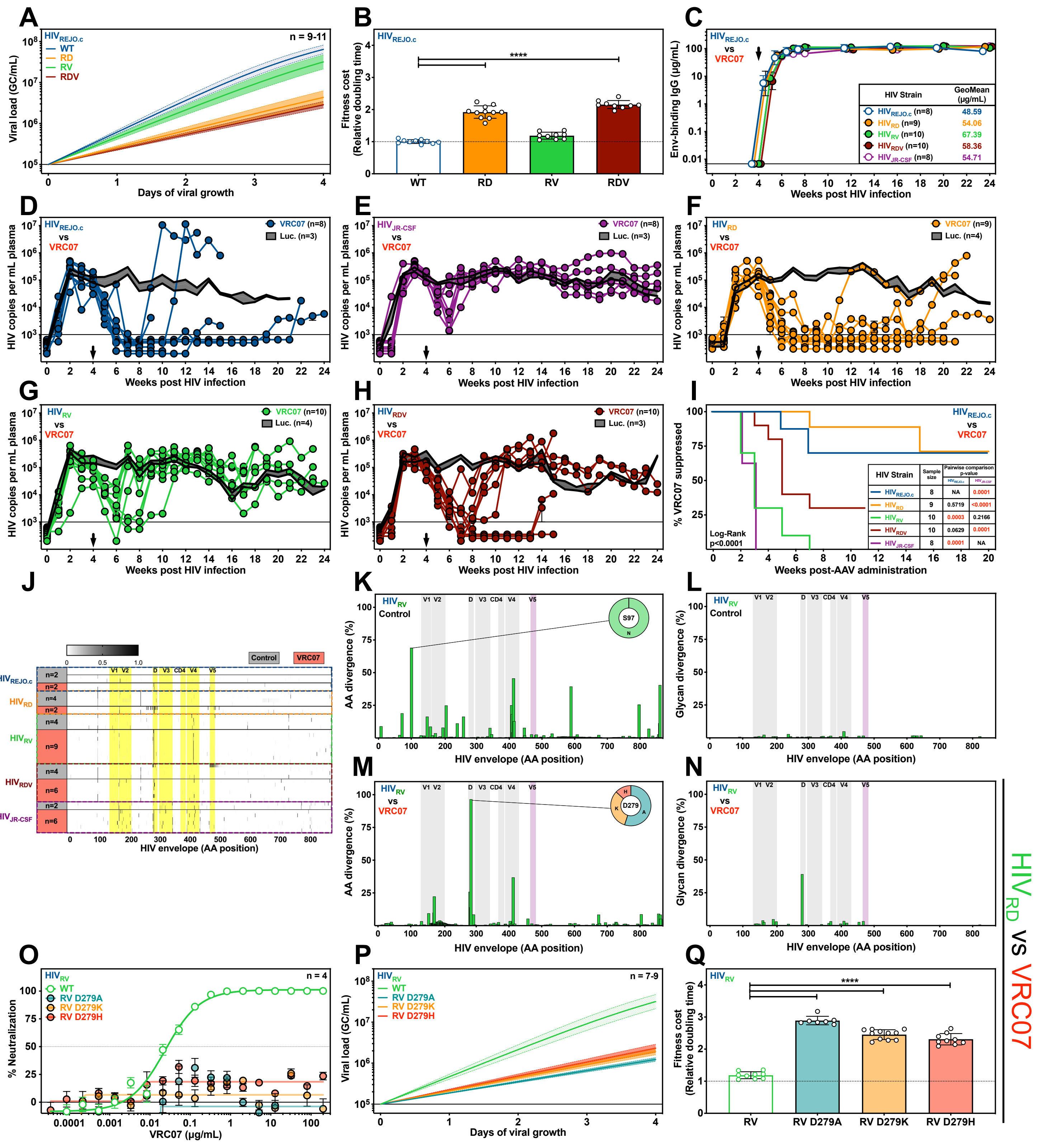


**Figure S9. Neutralization, growth rate, and infectivity of chimeric $HIV_{REJO.c}$ viruses (Related to Figure 3).**

**Figure S9. Neutralization, growth rate, and infectivity of chimeric $HIV_{REJO.c}$ viruses (Related to Figure 3).**

**(A)** Viral growth curves of $HIV_{REJO.c}$, $HIV_{RD}$, $HIV_{RV}$, and $HIV_{RDV}$ using the *QuickFit* assay. Data are plotted as mean ± 95% C.I.

**(B)** Relative viral growth of $HIV_{REJO.c}$, $HIV_{RD}$, $HIV_{RV}$, and $HIV_{RDV}$ using the *QuickFit* assay and normalized to the parental strain. Data are plotted as mean ± S.E.M., and statistical differences were assessed by a Kruskal-Wallis non-parametric ANOVA with Dunn's *post hoc* test to correct for multiple comparisons (****: p<0.0001).

**(C)** ELISA-based quantitation of gp120-binding antibodies in the serum of infected humanized mice following administration of $5x10^{11}$ genome copies (GC) of AAV-Luciferase or AAV-VRC07. Black arrow denotes vector administration. Data are plotted as geometric mean ± geometric SD.

**(D-H)** HIV viral load in plasma following intravenous infection of $HIV_{REJO.c}$ (**D**), $HIV_{JR-CSF}$ (**E**), $HIV_{RD}$ **(F)**, $HIV_{RV}$ **(G)**, and $HIV_{RDV}$ **(H)** following AAV-VRC07 administration. Data are presented as mean ± S.E.M.

**(I)** Kaplan-Meier plot of viral suppression in humanized mice infected with $HIV_{REJO.c}$, $HIV_{JR-CSF}$, $HIV_{RD}$, $HIV_{RV}$, and $HIV_{RDV}$ following AAV-VRC07 administration. The total model significance (p<0.0001) and pairwise comparisons against the $HIV_{REJO.c}$ or $HIV_{JR-CSF}$ were assessed independently using Log-rank (Mantel-Cox) tests. The p-value for each pairwise comparison is shown in the table.

**(J)** Haplotypes of individual mice infected with $HIV_{REJO.c}$, $HIV_{JR-CSF}$, $HIV_{RD}$, $HIV_{RV}$, and $HIV_{RDV}$ following either AAV-Luciferase (grey) or AAV-VRC07 (red) administration.

**(K-L)** Amino acid divergence **(K)** and Glycosylation site divergence **(L)** from the envelope gene of the $HIV_{RV}$ parental strain across all control-treated mice.

**(M-N)** Amino acid divergence **(M)** and Glycosylation site divergence **(N)** from the envelope gene of the $HIV_{RV}$ parental strain across all VRC07-treated mice.

**(O)** *In vitro* neutralization assay of $HIV_{RV}$ mutants identified as potential VRC07 escapes against VRC07. Data are plotted as mean ± S.E.M. Each data point was evaluated in quadruplicate.

**(P)** Viral growth curves of $HIV_{RV}$ mutants identified as potential VRC07 escapes using the *QuickFit* assay. Data are plotted as mean ± 95% C.I.

**(Q)** Relative viral growth of $HIV_{RV}$ mutants identified as potential VRC07 escapes using the *QuickFit* assay. Data are plotted as mean ± S.E.M. and statistical differences were assessed by a Kruskal-Wallis non-parametric ANOVA with Dunn's *post hoc* test to correct for multiple comparisons (****: p<0.0001).

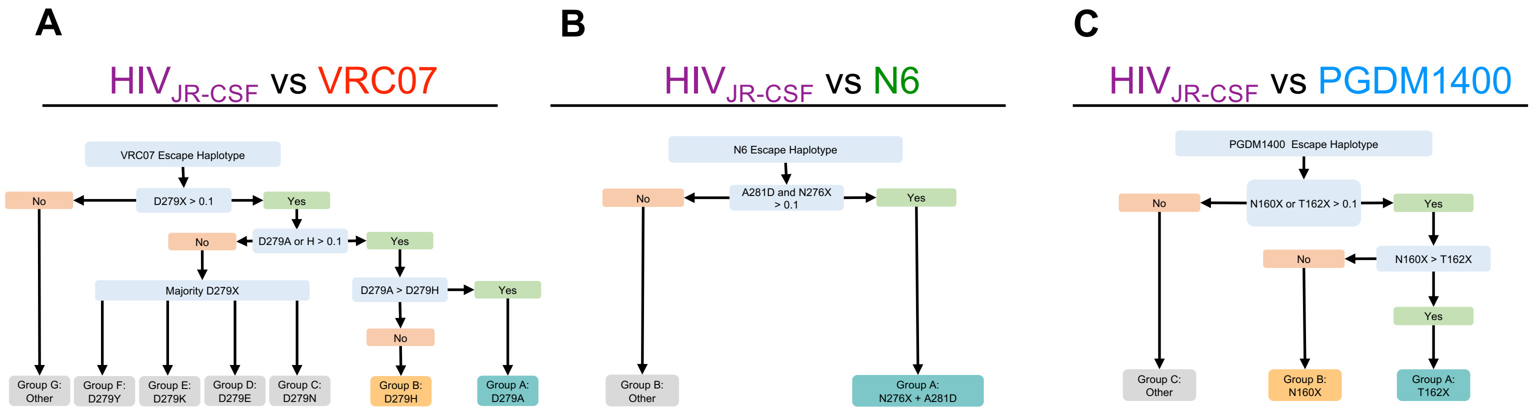


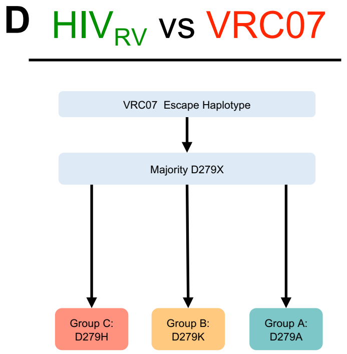


**E**

| Virus | Antibody | Fraction of Total Escapes | Escape Path | Raw AUC | Relative Fraction | Scaled AUC | Escape Barrier Score |
|---|---|---|---|---|---|---|---|
| HIV$_{JR\text{-}CSF}$ | VRC07 | 54% | A | 332 | 64% | 213 | 305 |
| | | | B | 259 | 36% | 92 | |
| HIV$_{JR\text{-}CSF}$ | N6 | 60% | A | 14 | 100% | 14 | 14 |
| HIV$_{JR\text{-}CSF}$ | PGDM1400 | 86% | A | 200 | 67% | 134 | 169 |
| | | | B | 106 | 33% | 35 | |
| HIV$_{RV}$ | VRC07 | 100% | A | 406 | 56% | 226 | 385 |
| | | | B | 360 | 33% | 120 | |
| | | | C | 347 | 11% | 39 | |

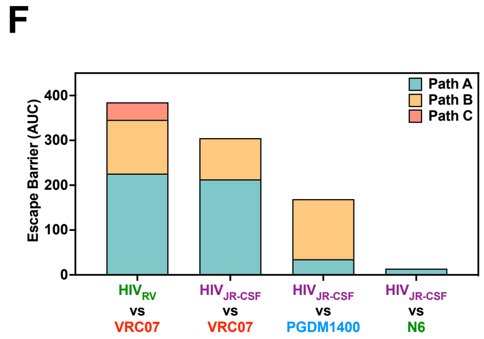


**Figure S10. Escape Barrier scores for HIV$_{JR\text{-}CSF}$ escape paths from various bNAbs and HIV$_{RV}$ escape paths from VRC07 (Related to Figure 3).**

**(A-D)** Escape path haplotype categorization algorithm for HIV$_{JR\text{-}CSF}$ escapes from VRC07 **(A)**, N6 **(B)**, PGDM1400 **(C),** and for HIV$_{RV}$ escapes from VRC07 **(D)**.

**(E)** Escape Barrier score calculations using paths A and B for HIV$_{JR\text{-}CSF}$ escapes from VRC07, N6, and PGDM1400, and for HIV$_{RV}$ escapes paths A, B, and C from VRC07.

**(F)** Escape Barrier (AUC) score denoting the aggregate fitness cost for each escape path as determined in the escapability maps. This score was calculated by adding the area under the escapability plot for each path A and path B viral escapes from VRC07, N6**,** and PGDM1400 and for paths A, B, and C for HIV$_{RV}$ escape from VRC07.

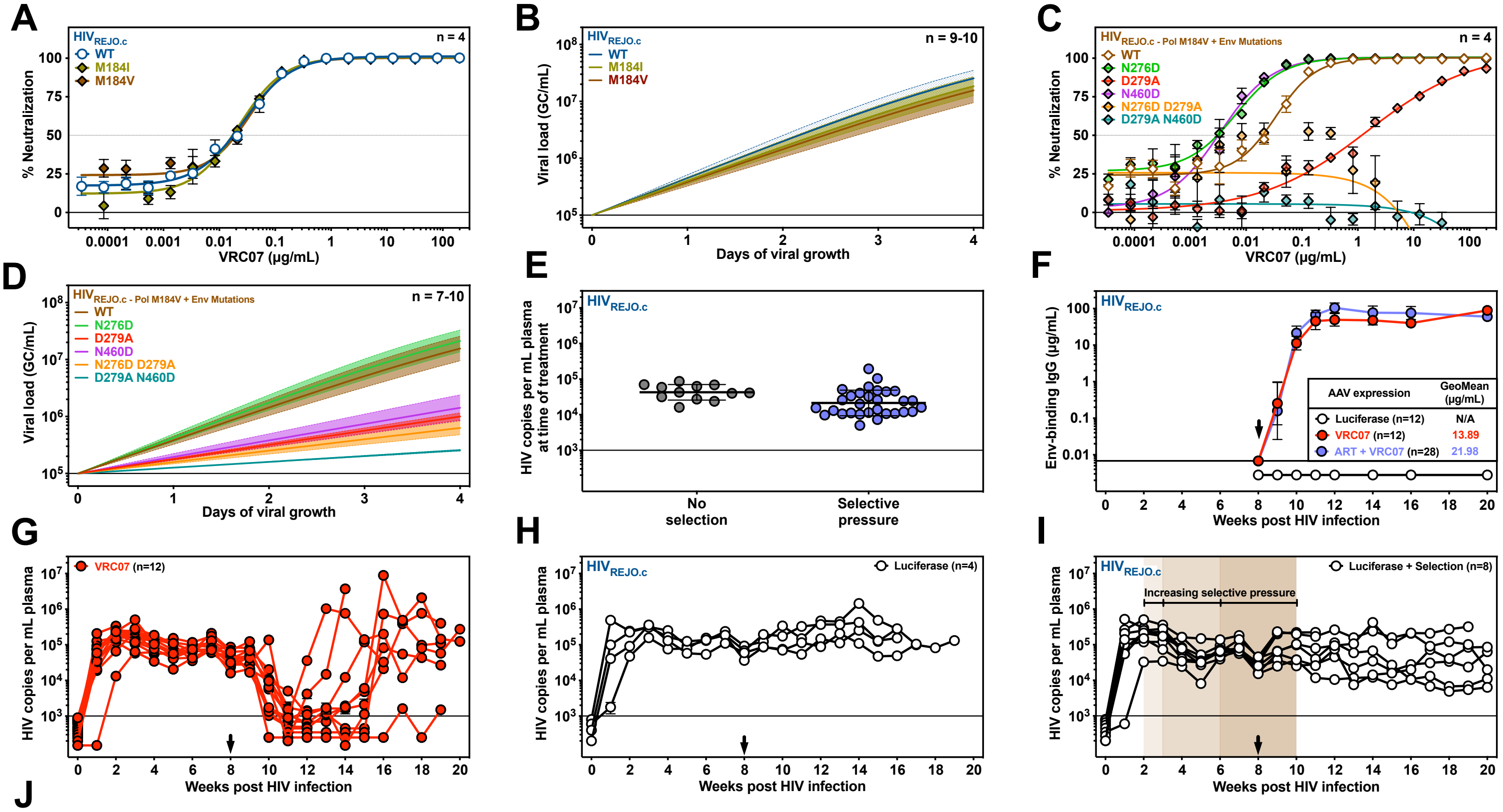


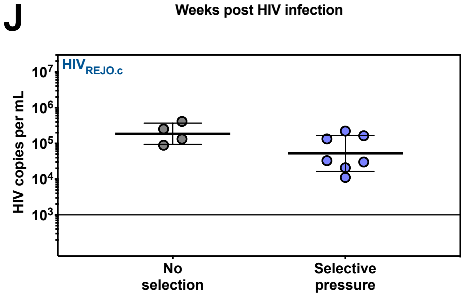


**Figure S11. Characterization of HIV$_{REJO.c}$ strains harboring ART-escape mutations during VRC07 escape (Related to Figure 4).**

**(A)** *In vitro* neutralization assay of HIV$_{REJO.c}$ Pol mutants against VRC07. Data are plotted as mean ± S.E.M. Each datapoint was evaluated in quadruplicate.

**(B)** Viral growth curves of HIV$_{REJO.c}$ Pol mutants determined using the *QuickFit* assay. Data are plotted as mean ± 95% C.I.

**(C)** *In vitro* neutralization assay of HIV$_{REJO.c}$ Pol mutants in combination with VRC07 escape mutants against VRC07. Data are plotted as mean ± S.E.M. Each datapoint was evaluated in quadruplicate.

**(D)** Viral growth curves of HIV$_{REJO.c}$ Pol mutants in combination with VRC07 escape mutants determined using *the QuickFit* assay. Data are plotted as mean ± 95% C.I.

**(E)** Geometric mean viral load of HIV$_{REJO.c}$-infected humanized mice that did (Selective pressure) or did not (No selection) receive ART prior to vectored VRC07 administration (p=0.1741, unpaired two-tailed Student's t test). Error bars indicate geometric SD.

**(F)** ELISA-based quantitation of gp120-binding human IgG antibodies in serum of HIV$_{REJO.c}$-infected humanized mice following injection of $5x10^{11}$ GC of AAV-Luciferase or VRC07. Black arrow denotes vector administration. Data are plotted as geometric mean ± geometric SD.

**(G-I)** HIV$_{REJO.c}$ viral load in plasma of humanized mice that did not receive ART before administration of AAV-VRC07 (**G**) or AAV-Luciferase (**H**) or that did receive ART before administration of AAV-Luciferase (**I**). Black arrows denote vector administration. Data are presented as mean ± S.E.M.

**(J)** Viral load of humanized mice that did (Selective pressure) or did not (No selection) receive ART after AAV-Luciferase treatment. The viral load was determined by calculating the geometric mean from 2 weeks post-treatment until either the mouse died or the experiment ended. Data are geometric mean ± geometric SD. (p=0.0795, unpaired two-tailed Student's t test).

**A**

| | Strain | | | |
|---|---|---|---|---|
| | $HIV_{REJO.c}$ | | $HIV_{JR-CSF}$ | |
| **Antibody** | **Average fold change over $IC_{50}$** | **Average $PT_{80}$** | **Average fold change over $IC_{50}$** | **Average $PT_{80}$** |
| **VRC07** | 2,338 | 611 | 901 | 238 |
| **N6** | 332 | 76 | 14 | 3 |
| **PGDM1400** | 3,753 | 454 | 83 | 15 |

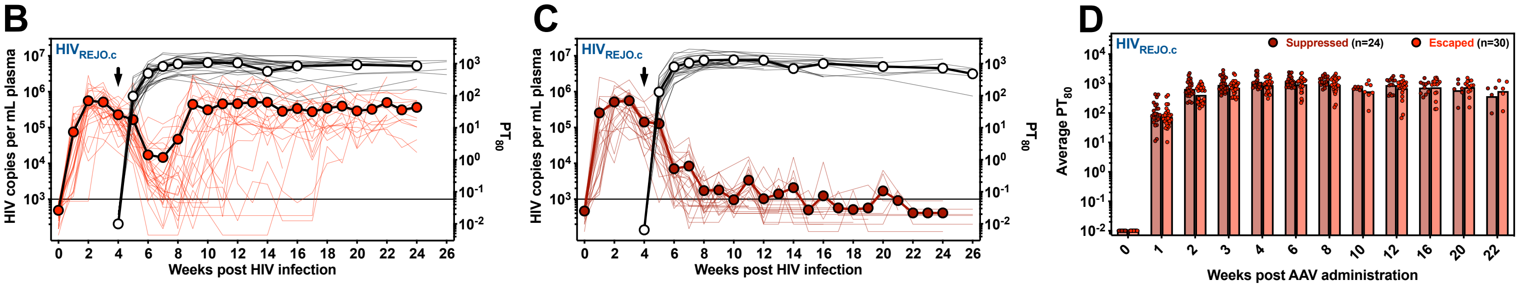


**Figure S12. Fold change over $IC_{50}$ and calculated $PT_{80}$ values for $HIV_{REJO.c}$- and $HIV_{JR-CSF}$-infected humanized mice expressing vectored bNAbs (Related to Figures 1 and 3).**

**(A)** Table describing the mean fold change above the $IC_{50}$ measured *in vitro*, and the corresponding mean $PT_{80}$ value achieved for each experimental condition.

**(B-C)** HIV viral load in plasma of $HIV_{REJO.c}$-infected mice (red lines) that escaped **(B)** or were suppressed **(C)** overlaid with the calculated $PT_{80}$ at each time point (black lines). Thin lines represent individual mice, and bold lines represent the average values for each group tracked over time. The sensitivity of qPCR was 1 genome copy per µL of plasma, and 5 µL were used in the reaction, resulting in a 1000 copy per mL limit of detection (solid line). Data are presented as mean ± S.E.M.

**(D)** Average $PT_{80}$ value for each independent mouse that was either suppressed (left bars) or escaped (right bars) tracked over time. No statistical differences in $PT_{80}$ were found at any time points between the groups ($p=0.1667$, Two-way ANOVA for Mixed effects model).

**Table S1. Neutralization resistance, dose-response curve slope, and relative fitness of HIV escape mutations from bNAbs.**

| Strain | Mutations | Antibody | $IC_{50}$ (μg/mL) | Dose-Response Curve Slope | Relative fitness |
|---|---|---|---|---|---|
| $HIV_{REJO.c}$ | Env WT | VRC07 | 0.0275 | 1.0330 | 1.00 |
| $HIV_{REJO.c}$ | Env WT | N6 | 0.0495 | 0.9405 | 1.00 |
| $HIV_{REJO.c}$ | Env WT | PGDM1400 | 0.0075 | 0.5638 | 1.00 |
| $HIV_{REJO.c}$ | Env N160K | PGDM1400 | 55.7800 | 1.6500 | 0.72 |
| $HIV_{REJO.c}$ | Env D167G | PGDM1400 | 81.4100 | 0.5694 | 0.82 |
| $HIV_{REJO.c}$ | Env N276D | VRC07 | 0.0033 | 0.8393 | 1.15 |
| $HIV_{REJO.c}$ | Env N276D | N6 | 0.1152 | 0.8393 | 1.15 |
| $HIV_{REJO.c}$ | Env T278K | N6 | 0.1201 | 4.0610 | 1.07 |
| $HIV_{REJO.c}$ | Env D279A | VRC07 | 1.4140 | 0.5011 | 2.36 |
| $HIV_{REJO.c}$ | Env A281D | N6 | 68.6800 | 1.7550 | 2.46 |
| $HIV_{REJO.c}$ | Env A281K | N6 | >200 | ND | 1.50 |
| $HIV_{REJO.c}$ | Env N460D | VRC07 | 0.0057 | 0.9295 | 0.65 |
| $HIV_{REJO.c}$ | Env N276D D279A | VRC07 | >200 | ND | 1.67 |
| $HIV_{REJO.c}$ | Env N276D A281D | N6 | 55.6600 | 1.3900 | 2.20 |
| $HIV_{REJO.c}$ | Env T278K A281D | N6 | >200 | ND | 1.92 |
| $HIV_{REJO.c}$ | Env D279A N460D | VRC07 | >200 | ND | 3.14 |
| $HIV_{JR\text{-}CSF}$ | Env WT | VRC07 | 0.0836 | 1.0440 | 1.00 |
| $HIV_{JR\text{-}CSF}$ | Env WT | N6 | 0.2715 | 1.0540 | 1.00 |
| $HIV_{JR\text{-}CSF}$ | Env WT | PGDM1400 | 0.0973 | 0.8354 | 1.00 |
| $HIV_{JR\text{-}CSF}$ | Env N160K | PGDM1400 | >200 | ND | 1.04 |
| $HIV_{JR\text{-}CSF}$ | Env N160Y | PGDM1400 | 63.3500 | 1.1730 | 1.14 |
| $HIV_{JR\text{-}CSF}$ | Env N160S | PGDM1400 | 46.5200 | 0.4690 | 1.01 |
| $HIV_{JR\text{-}CSF}$ | Env T162P | PGDM1400 | >200 | ND | 0.99 |
| $HIV_{JR\text{-}CSF}$ | Env T162N | PGDM1400 | >200 | ND | 1.00 |
| $HIV_{JR\text{-}CSF}$ | Env N276D | N6 | 0.0395 | 0.8759 | 1.15 |
| $HIV_{JR\text{-}CSF}$ | Env D279N | VRC07 | 0.1262 | 0.9310 | 0.93 |
| $HIV_{JR\text{-}CSF}$ | Env D279A | VRC07 | >200 | ND | 1.59 |
| $HIV_{JR\text{-}CSF}$ | Env D279H | VRC07 | >200 | ND | 2.32 |
| $HIV_{JR\text{-}CSF}$ | Env A281D | N6 | 6.5360 | 1.3900 | 3.40 |
| $HIV_{JR\text{-}CSF}$ | Env N276D A281D | N6 | >200 | ND | 2.04 |
| $HIV_{RD}$ | Env WT | VRC07 | 0.0355 | 1.1430 | 1.90 |
| $HIV_{RV}$ | Env WT | VRC07 | 0.0237 | 1.0050 | 1.17 |
| $HIV_{RV}$ | Env D279A | VRC07 | >200 | ND | 2.89 |
| $HIV_{RV}$ | Env D279H | VRC07 | >200 | ND | 2.30 |
| $HIV_{RV}$ | Env D279K | VRC07 | >200 | ND | 2.43 |
| $HIV_{RDV}$ | Env WT | VRC07 | 0.0332 | 0.8763 | 2.14 |
| $HIV_{REJO.c}$ | Pol M184I | VRC07 | 0.0226 | 1.0860 | 1.08 |
| $HIV_{REJO.c}$ | Pol M184V | VRC07 | 0.0343 | 1.2120 | 1.12 |
| $HIV_{REJO.c}$ | Pol M184V Env N276D | VRC07 | 0.0066 | 1.0220 | 1.04 |
| $HIV_{REJO.c}$ | Pol M184V Env D279A | VRC07 | 1.4650 | 0.4609 | 2.60 |
| $HIV_{REJO.c}$ | Pol M184V Env N460D | VRC07 | 0.0007 | 0.5980 | 2.25 |
| $HIV_{REJO.c}$ | Pol M184V Env N276D D279A | VRC07 | >200 | ND | 3.28 |
| $HIV_{REJO.c}$ | Pol M184V Env D279A N460D | VRC07 | >200 | ND | 6.29 |

### Table S2: Independent experiments performed for the data shown on Figures 1 and 2.

| | | | $HIV_{REJO.c}$-infected mice per treatment | | | | $HIV_{JR\text{-}CSF}$-infected mice per treatment | | | |
|---|---|---|---|---|---|---|---|---|---|---|
| **Experiment #** | **BLT donor** | **Total mice** | **Control** | **VRC07** | **N6** | **PGDM1400** | **Control** | **VRC07** | **N6** | **PGDM1400** |
| Experiment 1 | #1 | 33 | 17 | 16 | - | - | - | - | - | - |
| Experiment 2 | #2 | 28 | 10 | 9 | - | 9 | - | - | - | - |
| Experiment 3 | #3 | 42 | 15 | 12 | 15 | - | - | - | - | |
| Experiment 4 | #4 | 33 | 7 | 9 | - | - | 8 | 9 | - | - |
| Experiment 5 | #5 | 20 | - | - | - | - | 6 | - | 7 | 7 |
| Experiment 6 | #6 | 27 | - | - | - | - | 13 | 14 | - | - |
| Experiment 7 | #7 | 15 | 5 | - | 10 | - | - | - | - | - |
| Experiment 8 | #8 | 11 | 3 | 8 | - | - | - | - | - | - |
| **Total:** | | 209 | 57 | 54 | 25 | 9 | 27 | 23 | 7 | 7 |

### Table S3: Independent experiments performed for the data shown in Figure 3.

| | | | $HIV_{JR\text{-}CSF}$-infected mice per treatment | | |
|---|---|---|---|---|---|
| **Experiment #** | **BLT donor** | **Total mice** | **Control** | **ptN6** | **ptPGDM1400** |
| Experiment 11 | #11 | 22 | 7 | 7 | 8 |

### Table S4: Independent experiments performed for the data shown in Figure 3.

| **Experiment #** | Experiment 9 | **HIV Strain** | | | | |
|---|---|---|---|---|---|---|
| **BLT donor** | #9 | **$HIV_{REJO.c}$-infected mice** | **$HIV_{RD}$-infected mice** | **$HIV_{RV}$-infected mice** | **$HIV_{RDV}$-infected mice** | **$HIV_{JR\text{-}CSF}$-infected mice** |
| **Treatment** | **Control** | 3 | 4 | 4 | 3 | 3 |
| | **VRC07** | 8 | 9 | 10 | 10 | 8 |
| | **Total** | 11 | 13 | 14 | 13 | 11 |

### Table S5: Independent experiments performed for the data shown in Figure 4.

| | | | $HIV_{REJO.c}$-infected mice per treatment | | | |
|---|---|---|---|---|---|---|
| **Experiment #** | **BLT donor** | **Total mice** | **Control** | **Control+ART** | **VRC07** | **VRC07+ART** |
| Experiment 10 | #10 | 59 | 4 | 8 | 12 | 28 |

### Table S6: Frequency of primary escape mutations determined in control mice.

| **Mutation** | **Frequency** | | |
|---|---|---|---|
| | **$HIV_{REJO.c}$-infected control mice** | **$HIV_{JR\text{-}CSF}$-infected control mice** | **LANL HIV Sequence Database** |
| N160K | 14.320% | 0.932% | 2.143% |
| N160Y | 0.102% | 1.734% | 1.018% |
| N160S | 0.018% | 0.017% | 0.839% |
| T162P | 0.064% | 0.019% | 0.419% |
| T162N | 0.928% | 0.588% | 0.475% |
| D167G | 8.513% | 3.641% | 2.357% |
| N276D | 0.141% | 0.016% | 1.127% |
| T278K | 0.123% | 0.090% | 0.410% |
| D279A | 0.096% | 0.027% | 0.242% |
| D279N | 0.519% | 1.540% | 50.200% |
| D279H | 0.011% | 0.020% | 0.084% |
| D279K | 0.040% | 0.004% | 0.503% |
| A281D | 0.167% | 0.096% | 0.289% |
| A281K | 0.022% | 0.000% | 0.112% |
| N460D | 0.074% | 0.001% | 5.682% |
